# Phylogenomics unravels speciation patterns in temperate-montane plant species: a case study on the recently radiating *Ranunculus auricomus* species complex

**DOI:** 10.1101/2020.01.06.895904

**Authors:** Salvatore Tomasello, Kevin Karbstein, Ladislav Hodač, Claudia Paetzold, Elvira Hörandl

## Abstract

The time frame and geographical patterns of diversification processes in European temperate-montane herbs are still not well understood. We used the sexual species of the *Ranunculus auricomus* complex as a model system to understand how vicariance vs. dispersal processes in the context of Pleistocene climatic fluctuations have triggered speciation in temperate-montane plant species. We employed Target Enrichment sequence data from about 600 nuclear genes and coalescent-based species tree inference methods to resolve phylogenetic relationships among the sexual taxa of the complex. We estimated absolute divergence times and, using ancestral range reconstruction, we tested if speciation was rather enhanced by vicariance or dispersal processes.

Phylogenetic relationships among taxa were fully resolved. Incongruence among species trees mainly concerned the intraspecific relationships in *R. notabilis* s.l., *R. cassubicifolius* s.l., and the position of the tetraploid *R. marsicus*. Speciation events took place in a very short time at the end of the Mid-Pleistocene Transition (830-580 ka). A second wave of intraspecific geographical differentiation within and around the European mountain systems happened between 200-100 ka. Ancestral range reconstruction supports the existence of a widespread European ancestor of the *R. auricomus* complex. Vicariance processes have triggered allopatric speciation in temperate-montane plant species during the climatic deterioration occurred in the last phase of the Mid-Pleistocene Transition. Vegetation restructuring from forest into tundra could have confined these forest species into isolated glacial refugia. During subsequent warming periods, range expansions of these locally distributed species could have been hampered by congeneric competitors in the same habitat.

## 1 INTRODUCTION

Pleistocene climatic fluctuations have caused periodical range shifts (e.g., north-south) and/or range expansions and contractions in northern hemisphere plant species. The alternation of cold and warm periods has characterized the entire Quaternary (the last 1.8 Ma). In Europe, cold periods have produced southward shifts in the distribution ranges of temperate species (in the so-called “Mediterranean refugia”; Taberlet, Fumagalli, Wust-Saucy, & Cosson, 1998; Brewer et al., 2002) and/or survival of these in few, isolated central European refugial areas (e.g., Magri et al., 2006; Naydenov, Senneville, Beaulieu, Tremblay, & Bousquet, 2007). On the other hand, warmer interglacials have favoured range expansion towards north and the formation of contact zones among diverged populations of the same species (Hewitt, 1999; Magri, 2008). As a result, the survival of formerly coherent population groups in isolated refugia promoted allopatric speciation. Range expansions and the consequent formation of secondary contact zones resulted in hybridization, and eventually in the formation of new taxa through allopolyploidization (Stebbins, 1984; Abbott et al., 2013).

In the last three decades, many studies have elucidated distribution and diversification in European high alpine plant species (see Vargas, 2003; Schönswetter, Stehlik, Holderegger, & Tribsch, 2005). Although many studies have highlighted the importance of vicariance and postglacial recolonization for explaining phylogeographical patterns in alpine plant species (Taberlet et al., 1998; Comes & Kadereit, 1998; Widmer & Lexer, 2001; Kropf, Kadereit, & Comes, 2003), sometimes the situation is rather complicated and not easily explained by either vicariance or dispersal (Tribsch, Schönswetter, & Stuessy., 2002; Schönswetter & Tribsch, 2005; Reisch, 2008; Sanz, Schönswetter, Vallés, Scheeweiss, & Vilartesana, 2014; Tomasello & Oberprieler, 2017). Cold periods have not exclusively influenced the actual species distribution and intraspecific genetic patterns of alpine species, they have also actively promoted divergence and speciation, as documented in studies on animals (Knowles, 2001) and plants (Kadereit, Griebeler, & Comes, 2004). In the latter study, the authors observed that speciation processes in *Primula* sect. *Auricula* occurred principally during glacial periods in geographically isolated refugia.

Much less information is available on montane and/or subalpine plant species of the northern hemisphere. Research of forest plants focused on the dominating tree species, such as beech, oaks and pine (e.g., Magri, 2008; Brewer et al., 2002; Naydenov et al., 2007). However, herbaceous plants inhabiting the understory of cool and deciduous mountain forests of the temperate biome have hardly been investigated (Després, Loriot, & Gaudeul, 2002; Alsos, 2005). For the boreo-montane species *Polygonatum verticillatum* putative glacial refugia might have existed in the foothills of heavily glaciated mountain ranges, such as the Alps and the Carpathians (Kramp, Huck, Niketić, Tomović, & Schmitt, 2008).

The cosmopolitan genus *Ranunculus* L. with ca. 600 species originated and started diversifying in the early Miocene (Emadzade & Hörandl, 2011). Emadzade, Gehrke, Linder, & Hörandl (2011) have elucidated how both vicariance and long-distance dispersal have to be invoked to explain present distribution patters of lowland and montane species in the genus. Intercontinental and transoceanic dispersal of forest species happened mostly in the *R. polyanthemos* clade (Emadzade et al., 2011). Other clades, like *R*. sect. *Auricomus*, were so far not fully resolved. The Eurasian *Ranunculus auricomus* complex is nested as a monophyletic group within a big clade (*R*. sect. *Auricomus*) that comprises otherwise Asian, Arctic and North American species (Emadzade et al., 2015). Whereas apomictic polyploids of the *R. auricomus* complex usually occupy lowlands of the temperate and boreal biomes, sexuals are rather montane and subalpine species of Mediterranean and temperate European mountains, inhabiting deciduous forest ecosystems or natural and anthropogenic meadows. These taxa have restricted or disjunct distribution ranges, and usually occur at the foothills of the main European mountain ranges (Figure 1).

**Figure 1.**
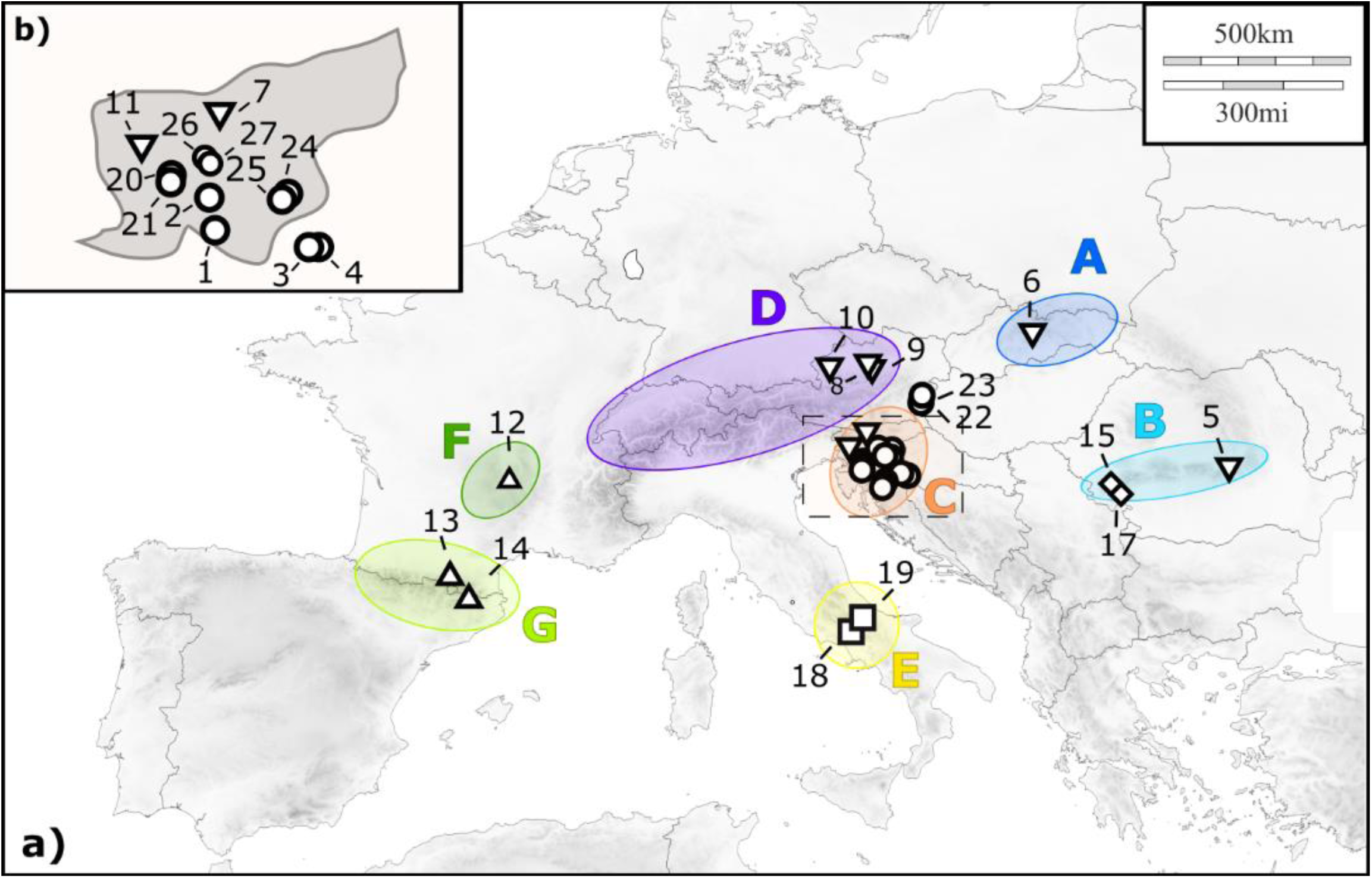
Distribution map of the accessions included in the present study. Numbers correspond to the map IDs in Table 1. Different symbols are for different species, following the treatment given in Karbstein et al. (2019): Circles are for *R. notabilis* s.l., diamonds for *R. flabellifolius*, squares for *R. marsicus*, triangles for *R. envalirensis* s.l. and overturned triangles for *R. cassubicifolius* s.l.. Coloured ellipses refer to the areas defined for the ancestral range reconstruction analysis: A) Northern Carpathians; B) Southern Carpathians; C) Illyrian region; D) Northern Prealps; E) Central Apennines; F) Massif Central; and G) Pyrenees.

**Table 1.**
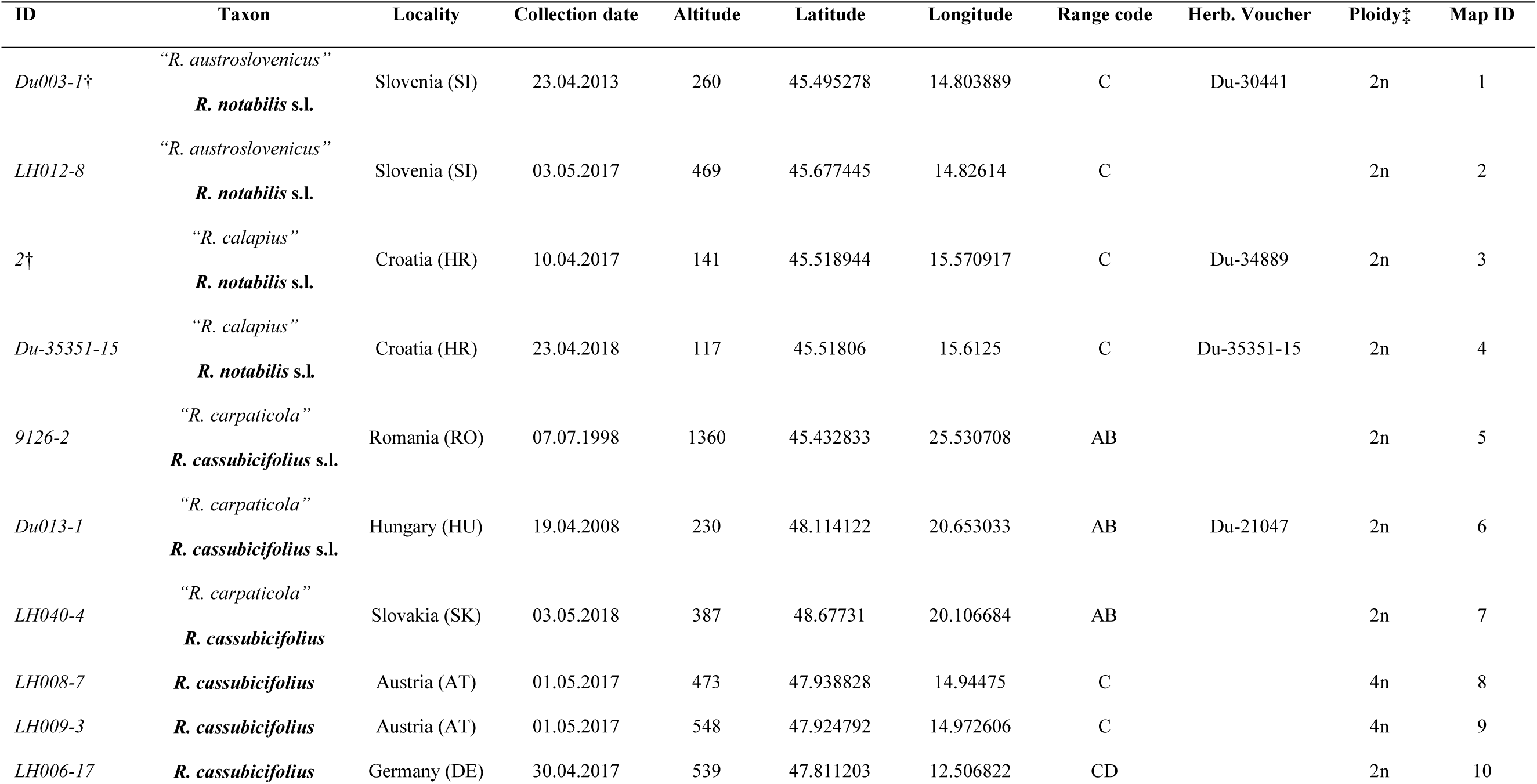
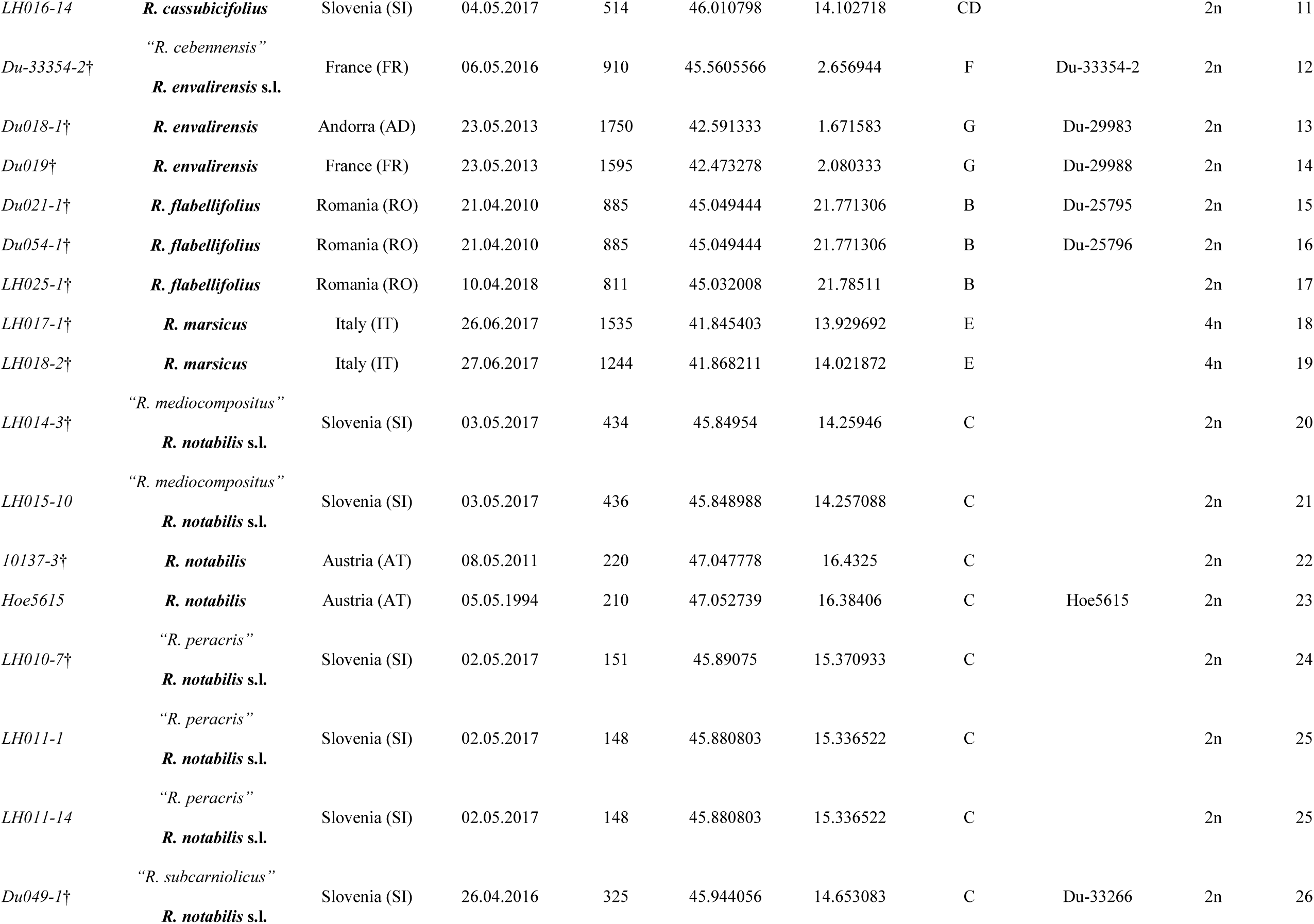
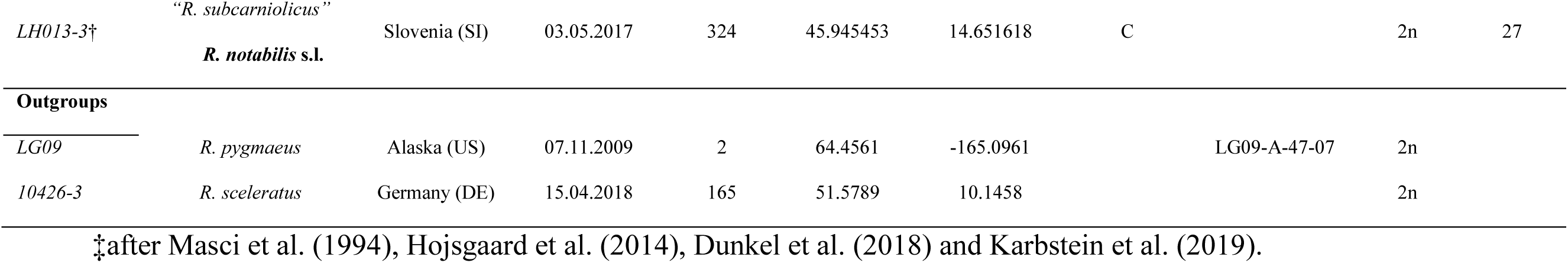
Information about the 28 accessions belonging to sexual taxa of the *R. auricomus* complex, and the two outgroup samples. Accepted names following Karbstein et al. (2019) in boldface, younger synonyms in inverted commas. Countries are abbreviated according to the ISO code 3166-2. Altitude is given in meter above sea level (m.a.s.l.). † indicates samples from type material or collected from “locus classicus” and nearby. Range codes are those used in BioGeoBEARS for the ancestral range reconstruction and correspond to those in Figure 1. Map IDs refer to the samples IDs given in Figure 1.

All taxa included herein are diploid or tetraploid sexuals (Masci, Miho, & Marchi, 1994; Hojsgaard et al. 2014; Dunkel, Gregor, & Paule, 2018). Hörandl (2004) estimated the divergence time between *R. notabilis* and *R. carpaticola* using isozyme data and Nei’s genetic distances (Nei, 1975) as 914,000 years ago. Apomictic hybrid taxa are hypothesized to be even younger (<100,000 years; Pellino et al. 2013). Up to now, traditional markers have failed in resolving the phylogenetic relationships among taxa of the complex (Hörandl et al., 2009). Karbstein, Tomasello, Hodač, Daubert, & Hörandl (2019) applied phylogenomic methods (RADseq and Target Capture), together with geometric morphometrics to investigate species boundaries among the sexual taxa of the complex. The authors concluded that five species are recognisable within the complex (*R. cassubicifolius* s.l., *R. envalirensis* s.l., *R. flabellifolius, R. marsicus* and *R. notabilis* s.l.; Table 1).

In the present study, we address two questions concerning the spatio-temporal diversification of temperate-montane species in Europe, using as study object the sexual species of the *R. auricomus* complex. (1) Did colder periods during glaciation cycles enhance speciation processes in temperate montane species? (2) Is vicariance or dispersal the most probable scenario to explain the present distribution of sexual taxa of *R. auricomus*? Therefore, we applied target enrichment of nuclear genes to: (i) resolve phylogenetic relationships among taxa of the complex; (ii) use coalescent-based methods to estimate the species divergence times in the context of past climatic oscillations; (iii) apply ancestral range reconstruction methods to test vicariance vs.dispersal speciation scenarios.

## 2 MATERIALS AND METHODS

### 2.1 Plant material

We included samples from all the described sexual species of the *R. auricomus* complex. Plant material used for DNA extraction consisted of silica-gel dried leaves or herbarium specimens (Table 1). We follow the taxonomy of Karbstein et al. (2019; see synonymy in Table 1). Two additional *Ranunculus* species not belonging to the *R. auricomus* complex were included in the analyses as outgroup: *R. pygmaeus*, from the clade of *Ranunculus* sect. *Auricomus*, and *R. sceleratus* from the next sister clade (Emadzade et al. 2011).

### 2.2 Gene selection, probe design

To find single-copy genes and select target regions for phylogenetic analyses, we used transcriptomes from two diploid species (*R. notabilis* and “*R. carpaticola*”), a tetraploid accession (*R. cassubicifolius*) (Pellino et al. 2013), and *R. brotherusii* (Chen, Zhao, Wang, & Moody, 2015), an Asian species of *R*. sect. *Auricomus*. MarkerMiner v1.0 (Chamala et al., 2015) was used to identify putative single-copy orthologous loci for probe design with *Arabidopsis thaliana* as reference. We selected exons found in at least two of the four transcriptomes, with lengths ranging from 120-960 bp and a minimum variability of two SNPs/120 bp, using a custom python script (available at: https://github.com/ClaudiaPaetzold/MarkerMinerFilter.git). We identified 2628 exonic regions belonging to 736 target genes (Supplemental Table S1). The selected target regions (genes) ranged from 121 to 5291 bp and included at least one exonic fragment each.

Arbor Biosciences (Ann Arbor, Michigan, USA) produced a MYbaits target enrichment kit with 20,000, 120 bp long in-solution biotinylated baits based on target sequence information. The final bait panel consisted of 17,988 probes, 14,632 of which unique, and tiling at 2 x density.

### 2.3 Sequencing, reads processing and alignment

Sequencing libraries were prepared using either the “NEBNext Ultra II DNA Library Prep Kit for Illumina^®^” (E7645) or the “NEBNext Ultra II FS DNA Library Prep Kit for Illumina^®^” (E7805) (New England BioLabs, Ipswich, USA). Sequencing was conducted on an Illumina MiSeq System (Illumina Inc., San Diego, USA) at the Transcriptome and Genome Analysis Laboratory (University of Goettingen, Göttingen, Germany). Pools were mixed equimolarly and sequenced in two different paired-end runs (6 pools, 24 samples each) with a 2x 250 bp (500 cycles) v2 kit.

Processing of the obtained raw reads was done using the pipeline HybPhyloMaker (Fér & Schmickl, 2018). We produced different sets of alignments by performing phasing or not, and by applying different filtering schemes for paralogs and alignment positions (see Supporting Information for details on library preparation and read processing).

### 2.4 Gene tree and species tree analyses

For all different datasets, species trees were estimated (i) using Maximum Likelihood (ML) as implemented in RAxML v8.2.4 (Stamatakis, 2014) on the concatenated datasets (concatML) and (ii) applying the coalescent-based method ASTRAL III (Chao, Rabiee, Sayyari, & Mirarab, 2018). For the concatML analyses, concatenated alignments were produced in HybPhyloMaker using AMAS and saved in the phylip format. Analyses were run on the Scientific Computer Cluster of the GWDG (https://www.gwdg.de/) using RAxML, with alignments partitioned by genes, the GTRGAMMA model and 100 bootstrap replicates. Concatenated analyses were only performed for the consensus alignments, since there was no way of knowing how the phased alleles of a sample combined across loci.

For the coalescent-based analyses, firstly gene trees (or exon trees in case of the allele alignments) were reconstructed with RAxML. Analyses were run with 100 standard bootstrap replicates, with the GTRGAMMA model and partitioning by exons. Gene tree characteristics (average bootstrap support, average branch length, etc.) and correlations among alignments and gene tree characteristics were calculated and plotted in *R* v3.5.2 (R Core Team, 2018), using modified scripts from Borowiec et al. (2016) as implemented in HybPhyloMaker. Gene trees were rooted, collapsing branches with bootstrap values <50, and combined into single NEWICK files using Newick Utilities (Junier & Zdobnov, 2010). Species trees were inferred applying the coalescent-based algorithm implemented in ASTRAL-III v5.6.3 (Chao et al., 2018) with 100 multi-locus bootstrap replicates. To assess the amount of gene tree conflict on branches, we measured the quartet support on the ASTRAL trees (Sayyari & Mirarab, 2016) for the main, the first alternative and the second alternative topologies (“-t 8” option in ASTRAL). In addition, we also quantified branch support and conflict on the concatML trees with the Quartet Sampling method (Pease, Brown, Walker, Hinchliff, & Smith, 2018).

### 2.5 Divergence time estimation

To obtain absolute divergence times, we used two secondary calibration points for the analyses. The crown age of the *R. auricomus* complex was set according to Hörandl (2004). In this study, the divergence time between *R. notabilis* and *R. carpaticola* was estimated as 914,000 years, whereas the split between *R. notabilis* and *R. cassubicifolius* as 535,000 years. Consequently, we applied a normally distributed tmrca prior to the crown age of the complex, with prior distribution ranging between the two age estimates (0.7245, ±0.189 Ma). The root of the tree (only analysis including the outgroup; see below) was calibrated according to the dated phylogeny published in Emadzade & Hörandl (2011). Therefore, we used a normally distributed tmrca prior for the root centred at 12.53 Ma (± 2.3 Ma).

Divergence times were estimated with BEAST v2.5.2 (Bouckaert et al., 2019) using the 50 gene alignments with the highest number of parsimony informative sites (PI) from those of the consensus dataset, without paralog filtering and cleaned from “phantom spikes”. We estimated the best fitting substitution model for each of the 50 loci (Table S2) using the Bayesian Information Criterion (BIC) in jModeltest v2.1.10 (Darriba, Taboada, Doallo, & Posada, 2012). Input files for the BEAST analysis were prepared in BEAUti v2.5.2 (Bouckaert et al., 2019). We used the *BEAST template and unlinked substitution models, clock models and gene trees for all loci. The strict clock was enforced and the average clock rate for one random locus was fixed to 1.66×10^-6^ while estimating all other clock rates in relation to this locus. Formerly described species were treated as independent lineages. Tetraploid accessions of *R. cassubicifolius* were treated separately from the diploid representatives of the species.

**Table 2.**
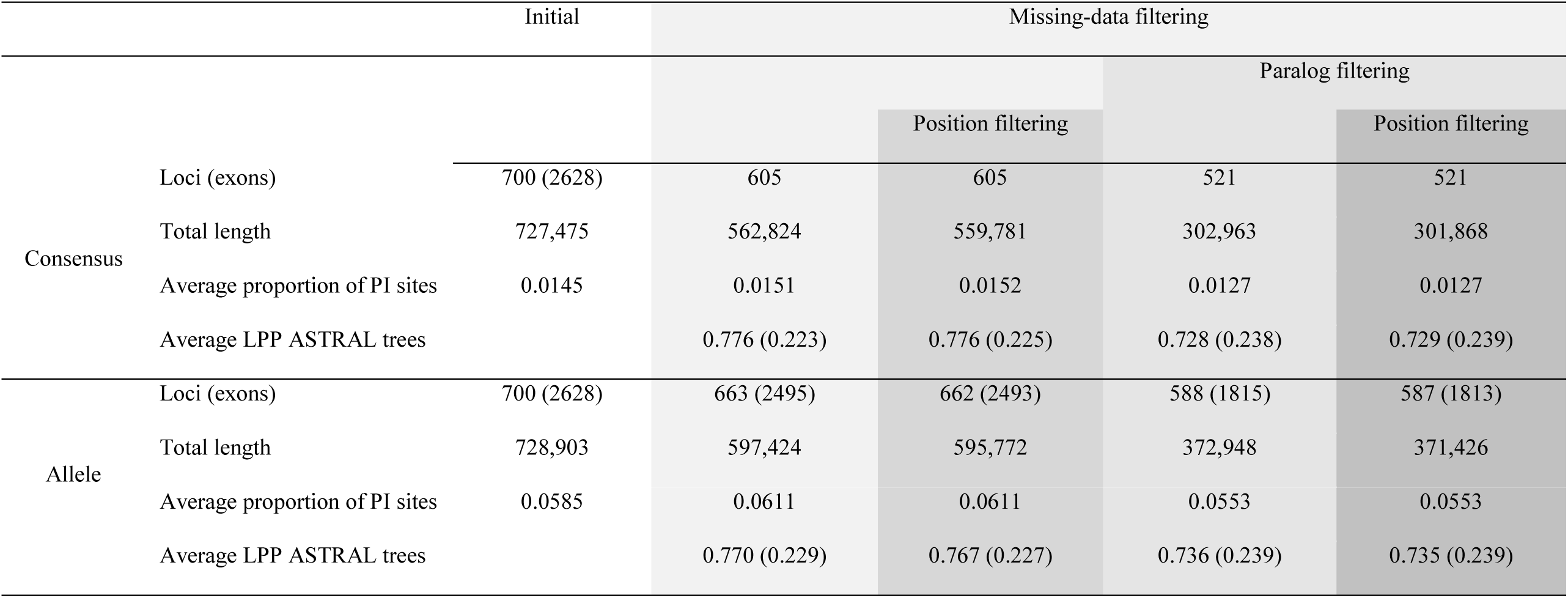
Alignment characteristics and performance of different filtering scenarios for the consensus and the allele dataset. For the Local Posterior Probability (LPP) of the ASTRAL bootstrap trees, mean values are given with standard deviation in brackets.

Since the main goal of the analyses was to estimate the time frame of speciation events, and since these analyses were based only on a reduced number of loci, we used a soft-constrained approach on the species tree topology in order to reach faster convergence and to make the topology of the *BEAST tree similar to the topology of the species trees obtained with the other methods. In particular, the monophyly of the clade with the Illyrian taxa was enforced.

Substitution models for each locus were adjusted to those found with jModeltest, and the Birth-Death model was used as prior on the species tree. Two independent analyses were run for 2×10^9^ generations, sampling every 100,000^th^ generation. ESS values and convergence between independent analyses were checked in Tracer v1.6 (Rambaut & Drummond, 2007). Results of the two analyses were merged using LogCombiner v2.5.2 (Bouckaert et al., 2019) applying a burn-in of 10%. Finally, the remaining 18,000 trees were used to construct a maximum clade credibility tree with a posterior probability limit set to 0.5 and “Mean Heights” for node heights using TreeAnnotator v2.5.2 (Bouckaert et al., 2019). Another set of analyses was performed as above-described, but including only the ingroup taxa, since many authors highlighted that using outgroups in coalescent-based analyses violates some of the model assumptions (Drummond & Bouckaert, 2015, Agudo, Tomasello, Alvarez, & Oberprieler, 2017).

### 2.6 Ancestral area reconstruction

To evaluate hypotheses about past distribution patterns and dispersal and/or vicariance events in the evolutionary history of the sexual representatives of the *R. auricomus* complex, we performed an ancestral area reconstruction with the *R* package BioGeoBEARS v1.1.2 (Matzke, 2018). This approach allows several models of biogeographic evolution to be compared via likelihood-based model selection (Matzke, 2013). Since sexual *R. auricomus* species grow within or near the main European mountain ranges, we assigned them to the seven following areas: Northern Carpathian (A), Southern Carpathian (B), Illyrian region (at the edge of the south-eastern Alps border; C), Northern Prealps (D), Central Apennines (E), Massif Central (F) and Pyrenees (G; Figure 1). As tree input, we used the *BEAST tree without the outgroup taxa, since the employment of widely distributed and non-uniformly sampled outgroups would have been counterproductive for the analyses. We specified a distance matrix (including scaled geographic distances among predefined areas), and activated the *x* parameter, which can be used to estimate dispersal probability as a function of distance (Van Dam & Matzke, 2016). Ancestral areas were estimated under all six available models DEC, DIVA-LIKE and BAYAREA-LIKE, with and without the jump-dispersal parameter *j*. We compared the different nested models (e.g., DEC+*x* against DEC) with a likelihood ratio test (LRT) and selected the overall best-fitting model using the Akaike Information Criterion (AIC).

## 3 RESULTS

### 3.1 Gene Capture and Sequencing

The average number of obtained reads per sample was 1,373,158 (867,000 to 2,178,952). Read quality was relatively high, and on average only 3.27% of reads per sample were discarded after quality check (Table S3). The average percent of mapped reads was 70.14%, with mapping success ranging from 53% to 82%, and only one sample (*R. notabilis* s.l.; 2) exhibiting a considerably lower number of mapped reads (26.89%; Table S4). Mapping success was not taxon dependent. The sample with the highest mapping percent belongs to the outgroup (*R. pygmaeus*, 82.48%). From the initial 736 target loci, 700 were successfully captured. In average, 612 loci per sample were captured, with LH008-7 (*R. cassubicifolius*) capturing best (621 loci), and *R. sceleratus* lowest (595 loci).

**Table 3.** Prior and posterior distributions of age estimates for the most important nodes of the *BEAST chronogram (Figure 4). Estimates are expressed in ka.

| Node description | Prior distribution |  | Posterior distribution |  |
| --- | --- | --- | --- | --- |
|  | Median age | 95% HPD interval | Mean age | 95% HPD interval |
| root | 724.5 | 914 -535 | 733.5 | 829.1-639.2 |
| *R. flabellifolius* stem age |  |  | 697.2 | 793.9-596.8 |
| *R. envalirensis* clade stem age |  |  | 683 | 780.6-577.1 |
| *R. marsicus*-*R. notabilis* s.l. split |  |  | 182.5 | 266.3-106.7 |
| *R. notabilis* s.l. crown age |  |  | 175.9 | 257.5-103.1 |

**Table 4.** Summary of data likelihoods under each model, and results of statistical model choice. Estimates for parameters dispersal parameter (*d*), extinction (*e*), the *x* (dispersal probability as a function of distance) and *j* (jump speciation events at cladogenesis) parameters. In bold the favoured model.

| Model | Max. No. areas | LnL | No. Parameters | $d$ | $e$ | $x$ | $j$ | AIC |
| --- | --- | --- | --- | --- | --- | --- | --- | --- |
| DEC | 7 | -26.228 | 2 | 0.210 | 0.497 | - | - | 56.457 |
| DEC+x | 7 | -22.757 | 3 | 2.275 | 1E-12 | -2.020 | - | 51.513 |
| DEC+j | 7 | -24.274 | 3 | 0.063 | 1.0E-12 | - | 0.054 | 54.548 |
| DEC+x+j | 7 | -22.597 | 4 | 1.213 | 1.0E-12 | -1.736 | 0.210 | 53.195 |
| DIVA | 7 | -21.475 | 2 | 0.123 | 1.0E-12 | - | - | 46.950 |
| **DIVA+x** | **7** | **-18.307** | **3** | **0.890** | **1.0E-12** | **-1.522** | **-** | **42.614** |
| DIVA+j | 7 | -21.473 | 3 | 0.121 | 1.0E-12 | - | 0.002 | 48.946 |
| DIVA+x+j | 7 | 17.718 | 4 | 2.450 | 1.0E-12 | -2.243 | 1.0E-05 | 43.437 |
| BAYAREA | 7 | -31.872 | 2 | 0.278 | 2.791 | - | - | 67.744 |
| BAYAREA+x | 7 | -29.506 | 3 | 2.976 | 2.911 | -1.771 | - | 65.012 |
| BAYAREA+j | 7 | -26.982 | 3 | 0.059 | 0.536 | - | 0.063 | 59.963 |
| BAYAREA+x+j | 7 | -25.354 | 4 | 0.336 | 0.218 | -1.253 | 0.302 | 58.709 |

### 3.2 Alignments and the impact of the different filtering schemes

The number of alignments used for the phylogenetic analyses varied according to the filtering scheme applied (Table 2). For the “consensus dataset”, about 15% of loci were excluded after missing data filtering and more than 25% after paralog filtering (more than 50% of the total alignment length). Concerning the “allele alignments”, working with exons made it possible to retrieve more loci (more than 80% and 50% of the total alignment length before and after paralog filtering, respectively). More details are given in Table 2.

Allele phasing generated alignments with a considerably higher proportion of PI sites compared to the consensus alignments (5.82% and 1.4% on average, respectively). Paralog filtering resulted in a loss of a fraction of PI sites in both the consensus and the allele datasets. When selecting the 50 loci for the BEAST analyses, 14.02% (78,458 bp) of the total matrix length was retrieved, comprising >23% of total PI sites (1,971). Lists of the alignments used in the different filtering schemes, their characteristics and substitution models of the loci selected for the BEAST analyses are given in Table S2.

### 3.3 Phylogenetic analyses and species trees

Species trees obtained from the concatML and coalescent-based analyses of the consensus dataset showed topological incongruences across most of the filtering scenarios. When using allele alignments, coalescent-based species trees obtained from different filtering treatments showed always the same topology, differing only slightly in support values and branch lengths. When taking the average local posterior probability (LPP) of the Astral bootstrap trees as criterion to compare different treatments, paralog filtering significantly worsened the results (in a post hoc Tukey’s test only differences between treatments with and without paralog filtering were significant; Table S5), whereas position filtering improved support values slightly in some cases (Figure S1). No significant differences were noticed between “consensus” and “allele” datasets. Accordingly, and as a matter of simplicity, only results of the “consensus” and “allele” analyses, without paralog filtering and with position filtering are shown in Figures 2 and 3. The rest of the trees are available as Supporting Information (Figures S2-S7).

**Figure 2.**
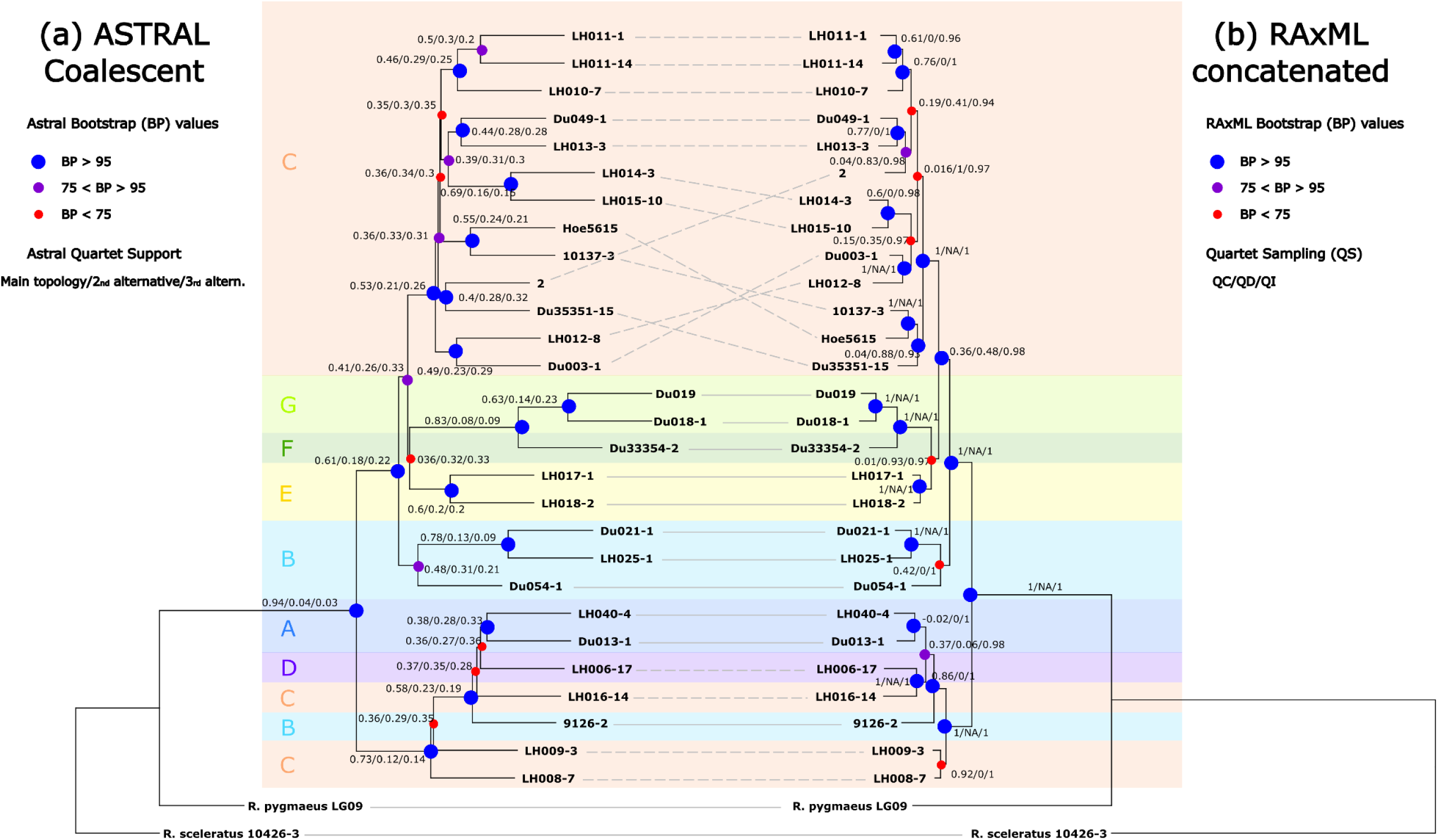
Tanglegram showing the coalescent-based Astral species tree (a) and concatenated maximum likelihood tree (b), based on the consensus dataset after applying position filtering and without paralog filtering. Continuous lines indicate accession with a congruent position, dashed lines indicate incongruence. Circles on the nodes refer to the bootstrap support values. Blue big circles are for support values above 95, purples medium-sized circles are for values between 75 and 95, and small red circles are for bootstraps below 75. Numbers at nodes indicate Astral quartet supports (a) and quartet sampling results on the maximum likelihood tree (b). Colours and letters refer to the areas defined for the ancestral range reconstruction, as indicated in Figure 1.

**Figure 3.**
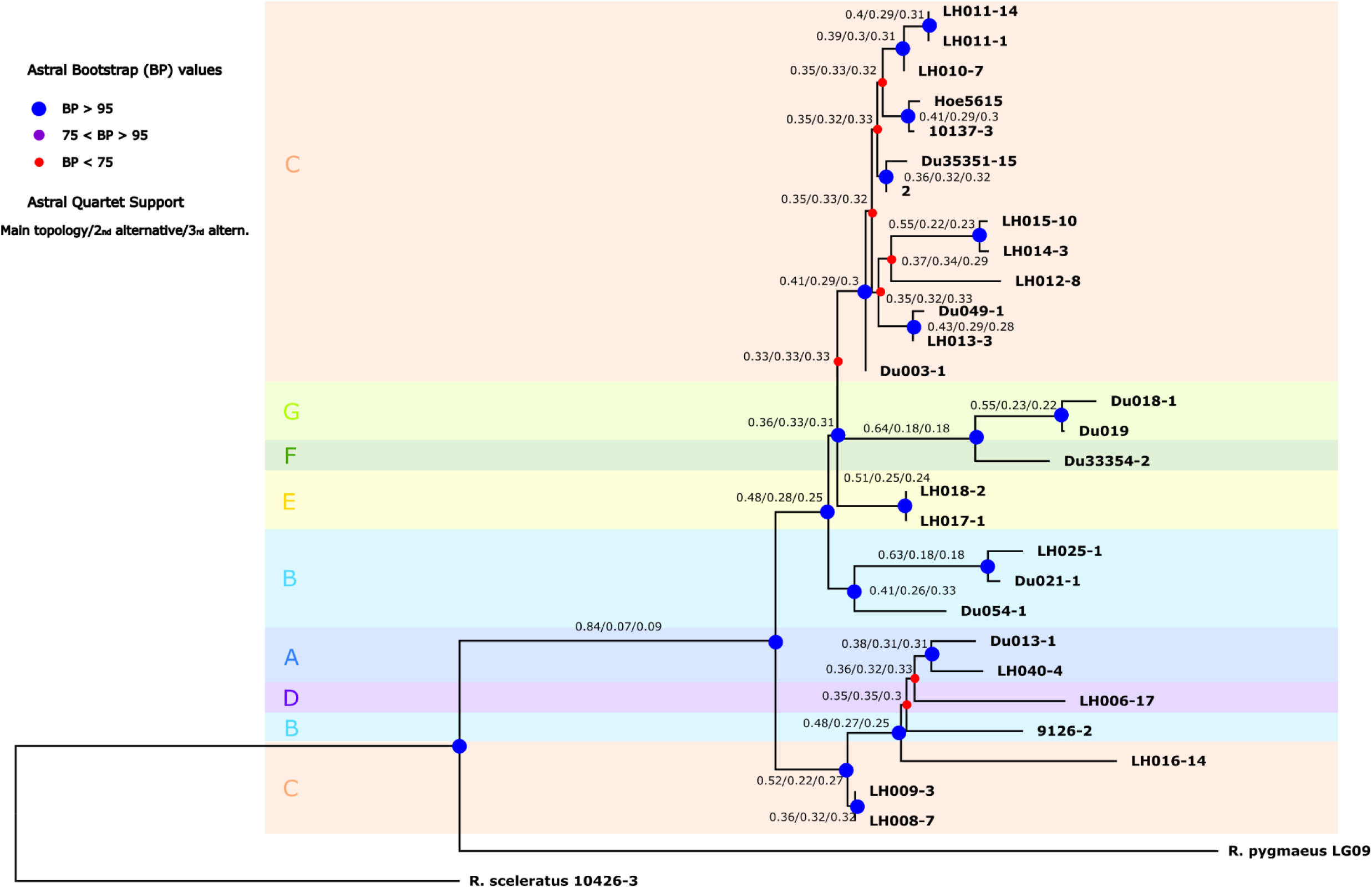
Astral species tree based on the allele dataset after applying position filtering and without paralog filtering. Circles on the nodes of the trees refer to the bootstrap support values. Blue big circles are for support values above 95, purples medium-sized circles are for values between 75 and 95, and small red circles are for bootstraps below 75. Numbers at nodes indicate Astral quartet support. Colours and letters refer to the areas defined for the ancestral range reconstruction, as indicated in Figure 1.

We detected two main incongruences among species trees. (i) The position of the taxa within *R. notabilis* s.l.. This clade is always highly supported, and in general samples cluster according to the taxonomy proposed by Dunkel et al. (2018), with exception of “*R. austroslovenicus”* and “*R. calapius”* that in some cases are found to be polyphyletic. However, support values are usually low within the clade and phylogenetic relationships among taxa change consistently across analyses. (ii) The position of *R. marsicus*, which often forms a clade with *R. envalirensis* s.l. but is resolved as sister to *R. flabellifolius* in some analyses. Support values of these phylogenetic relationships are generally low and branches very short.

Apart from these considerations, relationships among the main clades in the *R. auricomus* group are constant across analyses. The first split separates *R. cassubicifolius* s.l. from the rest of the taxa. The tetraploid *R. cassubicifolius* was well separated from the diploid accessions of the same species. In general, samples do not seem to cluster according to geographic origin within this clade. In the other main clade, samples cluster according to geographic distribution. *Ranunculus flabellifolius* (Southern Carpathians) occupies a basal position, whereas the clade with samples from the Massif Central and from the Pyrenees (*R. envalirensis* s.l.) is sister to the Illyrian clade. The position of the Apenninian *R. marsicus*, as mentioned above, is equivocal.

### 3.4 Age estimation

Results from the two *BEAST analyses (including and excluding the outgroup) were congruent to a large extent, varying only slightly on the age estimates. As for the concatML and the ASTRAL species tree, relationships within *R. notabilis* s.l. and within *R. cassubicifolius* s.l. are blurred and phylogenetic relationships receive low support (Figures 4 and S8).

**Figure 4.**
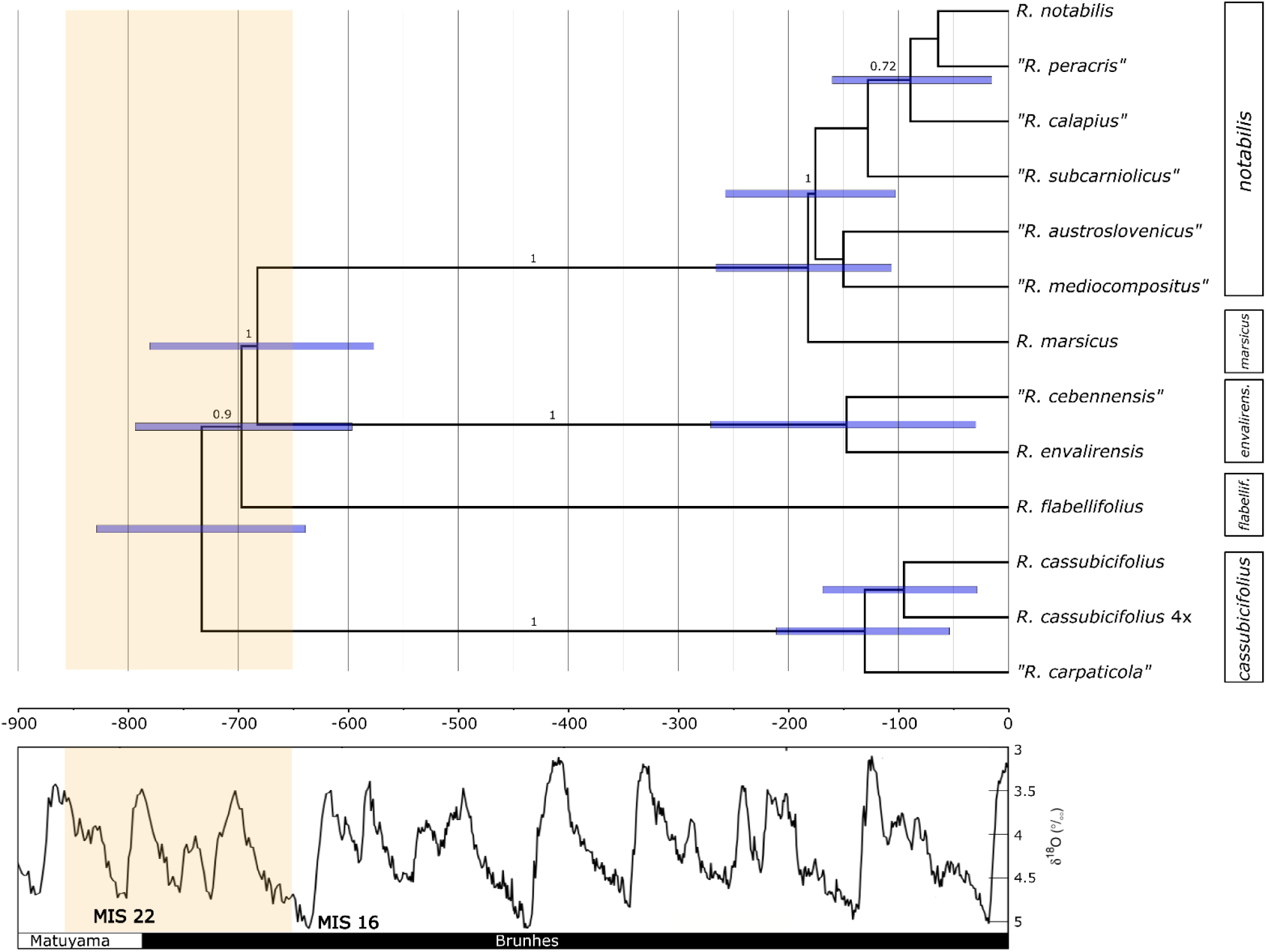
Maximum clade credibility tree estimated in *BEAST using the 50 most informative loci from the consensus dataset. Blue bars indicate 95% highest posterior density (HPD) intervals of the age estimate (see Table 3). Numbers above branches indicate Bayesian posterior probabilities. Only values above 0.7 are shown. On the right, bars indicate the current nomenclature according to Karbstein et al. (2019). Below the tree, the curve with the δ^18^O modified from Lisiecki & Raymo (2005). Valleys correlate to cold periods (glaciations) and peaks to warm periods (interglacials). Marine Isotope Stages (MIS) 22 and 16, representing the two most severe glaciations during the Mid-Pleistocene Transition (MPT), are also indicated. The orang region corresponds to the final phase of the MPT.

The crown age of the *R. auricomus* complex is estimated to be approximately 733.5 ka (Table 3), somewhat older (∼750 Ma) in the analyses including outgroups (Figure S8). Most of the speciation events occurred between 830 and 580 ka. The only exception is the Apenninian *R. marsicus*, which diverged from *R. notabilis* s.l. more recently (266-107 ka). However, the latter result does not seem to be supported by the concatML and by the species tree analyses, in which *R. marsicus* is rather found to be related to *R. envalirensis* (but see support values in Figures 2, 3), and its divergence seems to have occurred much earlier.

Differentiation within *R. notabilis* s.l., *R. envalirensis* s.l. and *R. cassubicifolius* s.l. occurred in the last 200 ka, with the crown ages of these three species estimated as 176, 147 and 130 ka, respectively (166, 128 and 119 ka in the analysis including outgroups; Figure S8).

### 3.5 Ancestral area

The Likelihood Ratio Test (LRT) shows that results of models including the *x* parameter were significantly better (*p* <0.05) than those without this parameter (Table S6). The *j* parameter did not improve the model fit for DEC and DIVA-LIKE, whereas it considerably ameliorated the BAYAREA-LIKE model fit. Overall, the best-fitting model was the DIVA-like+*x* (Table 4).

Results of the DIVA-LIKE+*x* model support a vicariance scenario, with the existence of a widespread ancestor for the *R. auricomus* complex (Figure 5a). The most probable ancestral range of the *R. cassubicifolius* s.l. was the Northern Carpathians and the Northern Prealps, while the ancestor of the remaining species most probably had a southern range, being distributed in the Southern Carpathians, Apennines and Pyrenees. However, some uncertainty is present in the estimation of the ancestral areas for the deepest nodes of the phylogeny, and statistical differences between most probable and alternative ancestral ranges are relatively low for some nodes (Figure 5b).

**Figure 5.**
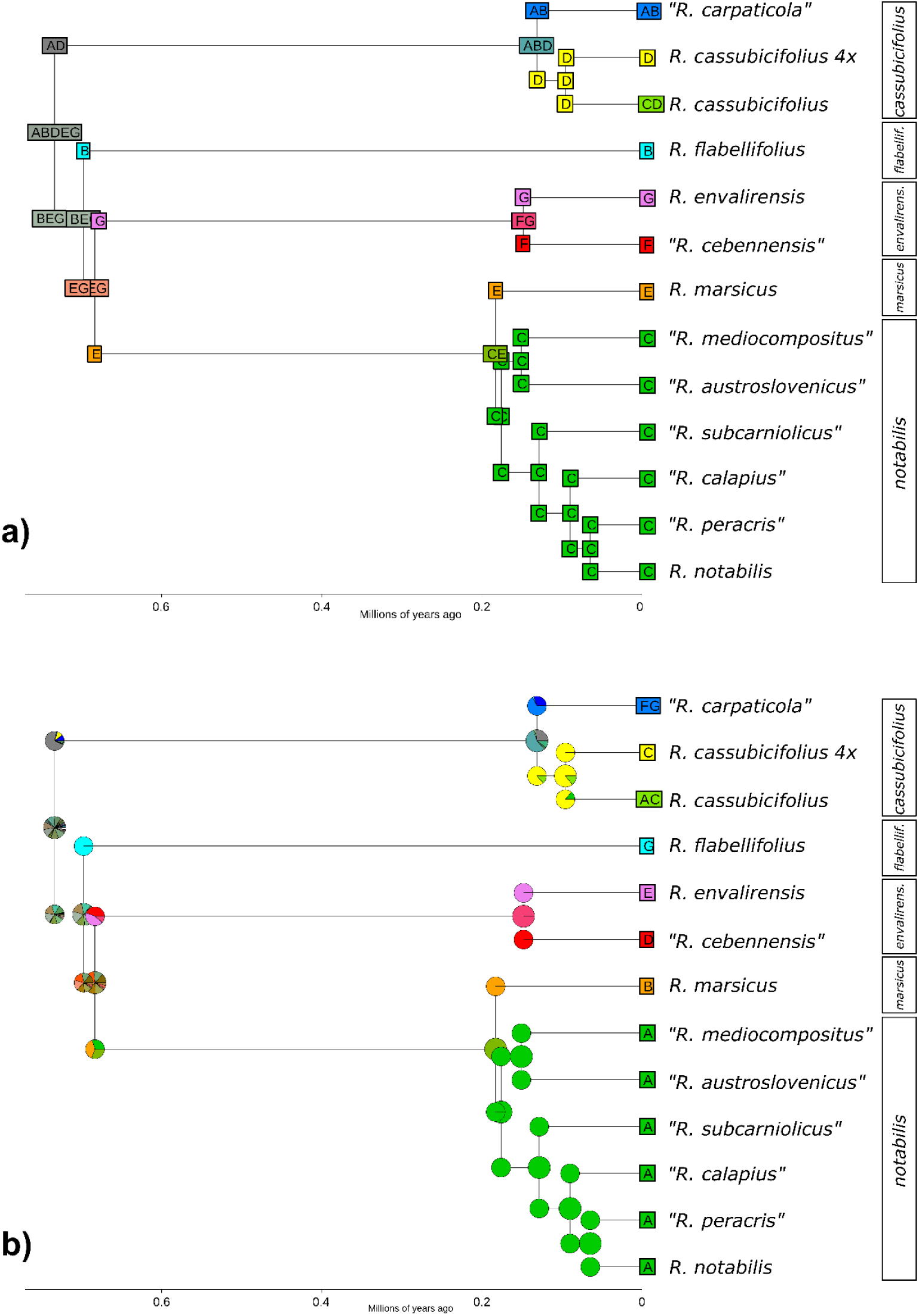
Ancestral state reconstruction for the sexual representatives of the *Ranunculus auricomus* complex, as estimated in BioGeoBEARS and using the favoured DIVALIKE+*x* model (a). Plot of the single-most-probable geographical range at each node (just before speciation) and post-split (just after speciation) (b). Pie charts representing the probabilities of each possible geographical range per node. In capital letters the areas defined for the analysis: A) Northern Carpathians; B) Southern Carpathians; C) Illyrian region; D) Northern Prealps; E) Central Apennines; F) Massif Central; and G) Pyrenees. Bars in the right part of the diagrams indicate the species delimitation according to Karbstein et al. (2019).

## 4 DISCUSSION

### 4.1 Phylogenetic relationships among sexual species of the *R. auricomus* complex

Disentangling phylogenetic relationships among recently and quickly diverged lineages remains a challenging task (Knowles & Chan, 2008). In the present paper, we tried to resolve the phylogeny of the sexual representatives of the less than one million-year-old *R. auricomus* complex. To this end, we analysed target enrichment data using concatenated ML and coalescent-based species-tree inference methods. Target enrichment is considered to be a very suitable approach for resolving deep phylogenies, as the baits are usually designed for highly conserved regions (McCormack et al., 2013; McKain, Johnson, Uribe-Convers, Eaton, & Yang, 2018). Nevertheless, the potential of this method for resolving shallow phylogenies and application in population genetic studies has been highlighted recently (Harvey, Smith, Glenn, Faircloth & Brumfield, 2016). Up to now, this sequencing approach has been applied a few times to resolve phylogenetic relationship in recently radiating plant and animal groups (e.g., the 7.2 Ma old genus *Heuchera* (Folk, Mandel, & Freudenstein, 2015); the ∼3 Ma old carnivorous plant *Sarracenia* (Stephens et al., 2015); the rapid island radiation of the Philippine Shrews (<4.8 Ma; Giarla & Esselstyn, 2015); and the lizard species group *Liolaemus fitzingerii* (2.6 Ma; Grummer, Morando, Avila, Sites Jr., & Leaché, 2018)). The crown age of the *R. auricomus* complex was estimated to be 733.5 ka (829-639 ka) herein. In this sense, our study represents the most outstanding example of employing target enrichment data in recently and rapidly radiating species complexes.

The main problems when working with rapidly radiating groups are: (i) rampant incomplete lineage sorting (ILS) due to the fact that ancestral polymorphisms are carried through a series of nearly simultaneous speciation events; and (ii) the lack of information due to the slow mutation rate of the (most often) coding regions used in studies employing a target enrichment approach (Giarla & Esselstyn, 2015). Using information from several (hundreds of) independent loci from multiple individuals per species, and applying coalescent-based species-tree methods should improve phylogenetic inference and overcome the above-mentioned problems (Kumar, Filipski, Battistuzzi, Pond, & Tamura, 2012). However, in Grummer et al. (2018) even 580 loci were not enough to provide significant support for interspecific relationships in the *L. fitzingerii* species group. In *R. auricomus*, 500 to 600 loci were able to resolve phylogenetic relationships among the main lineages of the complex. Incongruence was found mostly within the Illyrian group and within the clade of *R. cassubicifolius* s.l.. There, relationships among formerly described species were inconsistent across analyses and clades did not receive significant support, confirming the broader species delimitation provided by Karbstein et al. (2019). Diversification processes within these two species (i.e., *R. notabilis* s.l. and *R. cassubicifolius* s.l.) took place recently and in a very short time frame (i.e., the last 176 ka and 136 ka, respectively). We believe that the lack of phylogenetic divergence might be the cause of the badly resolved intraspecific relationships in these two species. As already observed in comparable complexes (Grummer et al. (2018); Stephens et al., (2015) for organellar genomes), there might have been too little time for the genomes to accumulate enough informative polymorphisms.

When looking at the backbone of the phylogenetic trees, the main source of incongruence is the tetraploid *R. marsicus*, which exhibits inconsistent positions in different concatML analyses, and in the ASTRAL and *BEAST species trees. However, all these sister relationships are not supported and the branches describing them are usually very short. Possible causes of these contrasting patterns can be either ILS or hybridisation/introgression. An allotetraploid origin of *R. marsicus* with the parental contribution of *R. notabilis* and of an ancestral relative of *R. envalirensis* or *R. flabellifolius* might be a realistic scenario. However, the high QD values of the quartet sampling (for the concatML; Figure 2), and the comparable frequencies of the two alternative topologies in the quartet support of the ASTRAL trees, sustain the hypothesis of ILS as the cause of incongruence rather than hybridization (Figure 3; Figures S2-7).

Concerning differences among tree inference methods (concatenated vs. coalescence-based) and performance of dataset from different filtering settings, we note that concatenation usually resulted in higher bootstrap values (compared to equivalent coalescent-based analyses) even though topologies were more discordant among trees. Similar patterns were already observed in empirical and simulation studies (Kubatko & Degnan, 2007; Weisrock et al., 2012; Herrando-Moraira et al., 2018). Remarkably, trees obtained by applying coalescent-based analyses to the phased dataset were topologically identical, and minor differences concerned only branch lengths and support values (Figures S5-S7). This is a further evidence that using allelic information together with coalescent-based species tree approaches can considerably increase the trustworthiness of phylogenetic analyses (Andermann et al., 2019). Concerning the performance of different phasing and filtering settings, the paralog filtering implemented in HybPhyloMaker significantly worsened branch supports. Applying this option on diploids possibly resulted in the elimination of many non-paralog (but variable) alignments. No significant amelioration in support values of the bootstrap ASTRAL trees was produced when applying phasing (but see above considerations on tree topologies) or position filtering (Figure S1).

### 4.2 Time frame of diversification processes

The crown age of the *R. auricomus* complex was estimated to be about 730 ka (Table 3). Hörandl (2004) estimated the divergence time between three sexual species using isozyme allelic frequencies and genetic distances after Nei, (1975). Ages of these divergence events were estimated to 914,000 and 535,000 years, respectively, even though they describe the same split in the phylogeny of the complex (the crown age of the whole group). Therefore, using a broad prior for the crown age of the complex and a relatively informative prior for the clock rate (according to the mutation rates estimated for the complex by Pellino et al., 2013), we were able to estimate the age of this node more precisely. Moreover, we obtained the time frame of all speciation events among sexual representatives of *R. auricomus*. Interestingly, almost all these events have taken place in a very short time between 830 and 580 ka (Figure 4). The only exception is the tetraploid species *R. marsicus*, which seems to have diverged much more recently (but see considerations about uncertainty above). The divergence estimates for all these speciation events coincide nicely with the MPT (also known as known as Mid-Pleistocene Revolution; Lisiecki & Raymo, 2005; Clark et al., 2006), one of the most important climatic changes occurring in the Pleistocene. During this interval (1.2 Ma - 650 ka) the relatively low-amplitude 41-ka climate cycles of the earlier Pleistocene were progressively replaced by high-amplitude 100-ka cycles (Lisiecki & Raymo, 2005). Glaciations became longer and more severe, and the average global ice volume increased consistently, with larger increases at high and middle latitudes of Eurasia and North America (Raymo, Lisiecki & Nisancioglu, 2006; Elderfield et al., 2012). This process became progressively stronger, and evidence from all northern continents indicates that the Marine Isotope Stage (MIS) 22 (∼870-880 ka) was the first of the major glaciation events characterising the later Pleistocene (Head & Gibbard, 2005). At the end of the Mid-Pleistocene climatic transition (MIS 16; ∼650 ka), the most substantial glaciation yet experienced in the northern hemisphere took place (Head & Gibbard, 2005). This tremendous climatic transition was probably the consequence of the increase of atmospheric moisture provided by warm subtropical water in the Western Mediterranean region that, combined with a decrease in boreal summer insolation, contributed to the feed of the ice-sheets in Central and Northern Europe (Sánchez-Goñi et al., 2016; Bahr et al., 2018). The onset of the global cooling phase and ice expansion caused a major restructuring of the global vegetation and faunal systems (Alroy, Koch, & Zachos, 2000; Gaboardi, Deng, & Wang, 2005; Zhao et al., 2017; Zhou et al., 2018).

Diversification processes within species took place more recently, most probably within the last two glaciation cycles. Diverging populations of *R. envalirensis* s.l., *R. notabilis* s.l. and *R. cassubicifolius* s.l. differentiated allopatrically (in the first case) or parapatrically (in the latter two cases) within the last 160 ka (Figure 4). As for the alpine taxa, probably also for montane plant species the last couple of glaciation cycles were a triggering force shaping intraspecific genetic patterns, but not sufficiently long to complete speciation processes. Overall, our age estimates are concordant with those found in previous studies (Hörandl, 2004; Emadzade & Hörandl, 2011; Pellino et al., 2013). In Hörandl (2004) and in Emadzade & Hörandl (2011) diversification processes within *R. cassubicifolius* were found to be slightly older, 317 ka and 640 ka, respectively. In our analyses, sexual tetraploids in *R. cassubicifolius* were found to be approximatively 95,000 years old. This result is in accordance with previous studies estimating the maximum age of hexaploid apomictic derivatives as 80,000 years (Pellino et al., 2013), and confirms the general assumption that the origin of polyploids is associated with the last glaciation cycle.

### 4.3 Biogeographical history

In our biogeographical analyses, the AIC preferred models supporting the existence of a widespread ancestor for the *R. auricomus* complex, stressing the importance of vicariance in triggering speciation events among sexual representatives of the complex (Table 4). The ancestor of the whole complex was reconstructed to be distributed in whole of Europe, although with some uncertainty (Figure 5). A first vicariance event separated populations with a northern distribution (giving rise to *R. cassubicifolius* in the Northern Prealps and the Northern Carpathians) from those with a southern distributional range. Further vicariance events then gave rise through allopatric speciation to the remaining sexual species of the complex (i.e., *R. flabellifolius*, *R. envalirensis*, *R. marsicus* and *R. notabilis*).

Almost all of these vicariance events occurred in a restricted time period, coinciding with the final stage of the MPT. At the beginning of this drastic climatic change, humid and mild temperate-forests with deciduous oaks and tertiary elements (such as *Carya* and *Zelkowa*) occupied a vast area of continental Europe (Stuchlik & Wójcik, 2001; Pastre et al., 2007; Magri & Palombo, 2013). Such environments must have been similar to the ecosystems inhabited nowadays by many of the sexual representatives of the complex. Climatic deterioration (the progressive cooling and drying of the climate) resulted in the fragmentation of such environments and in the establishment of grass-dominated open vegetation adapted to cold and arid conditions (Magri & Palombo, 2013). Towards the end of the MPT, tundra vegetation was well established in Central and Northern Europe (Stuchlik & Wójcik, 2001). A similar situation was registered in temperate Eastern Asia, where broadleaved mixed forest dominated by *Quercus* and *Pinus* shrank significantly during the MPT, transforming into grass-dominated vegetation definitively by around 0.7 Ma (Zhou et al., 2018).

Areas at the foothills of mountain systems might have acted as refugia for species of deciduous temperate forest ecosystems. Indeed, for the boreo-montane *P. verticillatum* putative glacial refugia were identified in the area surrounding the glaciated Alps and in the foothills of the Carpathian and Balkan mountain systems (Kramp et al., 2008). The ancestor of the *R. auricomus* complex might have therefore inhabited the deciduous temperate forests widespread in Europe during the first phases of the MPT. Climatic deterioration and the retreat of temperate forests to isolated refugial areas efficiently separated *R. auricomus* populations, which finally gave rise to the different sexual species of the complex by allopatric speciation. A similar pattern of allopatric speciation in the Mid Pleistocene was inferred for the temperate-montane Asian species *Dysosma versipellis*. (Qiu, Guan, Fu, & Comes, 2009) Surprisingly, the sexual species of the *R. auricomus* complex did not show long distance dispersal, as it has been observed in many other plant genera (Knapp et al. 2005), and also in other forest species from other clades of the genus *Ranunculus* (Emadzade et al., 2011). The achenes as diaspores are not much differentiated. The absence of long-distance dispersal in the sexual *R. auricomus* species may be due to more limited abilities to adapt to new environments or to novel niches. Outcrossing and self-sterility in the diploids may be other reasons for low colonization abilities (Hörandl. 2008).

A second wave of intraspecific vicariance occurred around 200-100 ka in the Alpine-Carpathian system (*R. cassubicifolius* s.l.) between the Pyrenees and the Massif Central (*R. envalirensis* s.l.) and, to a lesser extent, within the Illyrian clade (*R. notabilis* s.l.). Most taxa occur nowadays in very restricted areas with few populations, although appropriate habitats would be broadly available. The rapid range expansion of younger, polyploid apomictic derivatives of the *R. auricomus* complex could have blocked sexual progenitors (Hörandl, 2006). However, also other distantly related and widespread species of the genus occupying the same habitats (e.g., *R. polyanthemos* group, or *R. lanuginosus*) could have been competitors for sexual taxa of *R. auricomus*.

## Supporting information

see Supporting Information

## Acknowledgments

We thank the German Research Foundation for project funding (DFG, Ho4395/10-1) to E.H., within the priority program ‘Taxon-Omics: New Approaches for Discovering and Naming Biodiversity’ (SPP 1991), and the Botanical Museum, University of Oslo (O) for the material of *R. pygmaeus*. We also thank Mr. F.G. Dunkel for providing herbarium vouchers and living material for the recently discovered diploid populations from the Illyrian region and from France.

## Data Accessibility Statement

The raw reads were deposited in GenBank under the BioProject XXX. Assemblies, and Target alignments are archived in XXX.

## Author contributions

S.T. and E.H. conceived the ideas; S.T., K.K., L.H., and E.H. collected materials and data; S.T. and C.P. analysed the data; and S.T. wrote the paper with contributions from all co-authors.

