## Supplementary material for "Phylogenomics unravels speciation patterns in temperate-montane plant species: a case study on the recently radiating *Ranunculus auricomus* species complex": Figures S8

**Figure S8** Maximum clade credibility tree estimated in *BEAST using the 50 most informative loci from the consensus dataset and including outgroup samples. Blue bars indicate 95% highest posterior density (HPD) intervals of the age estimate. Numbers above branches indicate Bayesian posterior probabilities. Only values above 0.7 are shown. On the right, bars indicate the current nomenclature following Karbstein et al. (2019).

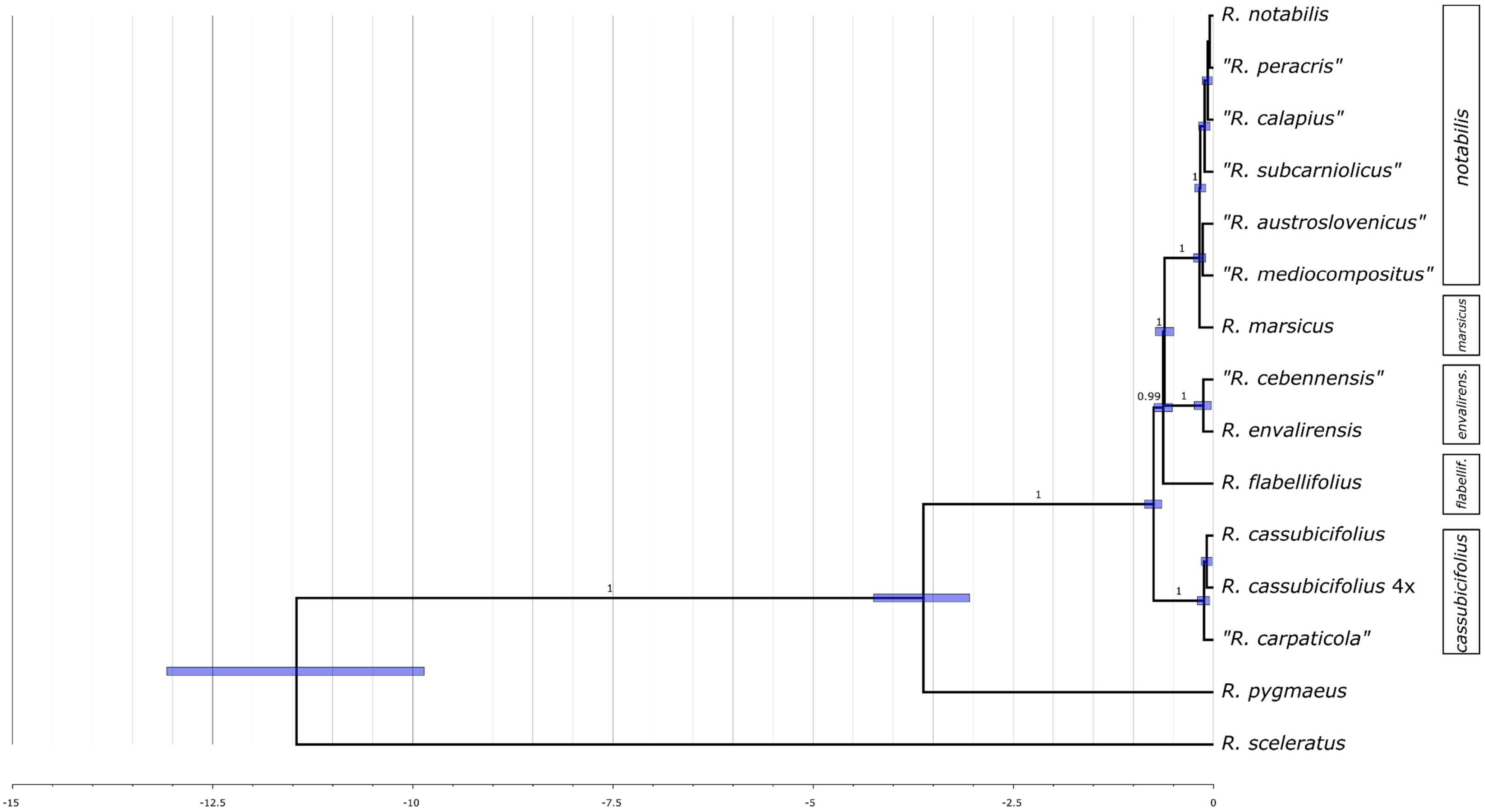
