## Supplementary material for "Phylogenomics unravels speciation patterns in temperate-montane plant species: a case study on the recently radiating *Ranunculus auricomus* species complex": see Supporting Information

### SUPPLEMENTARY METHODS

#### Library Preparation

Genomic DNA was extracted from ~1.5 cm² leaf material of silica-dried samples or herbarium collection using the Qiagen DNeasy Plant Mini Kit® (Qiagen, Hilden, Germany), following the manufacturer's instructions, except for the sample incubation time in the lysis buffer, which was increased to one hour. DNA quality and fragments length were checked by gel electrophoresis in a 1.5 Agarose gel and using the Roti®-Load DNAstain 3 (Carl Roth, Karlsruhe, Germany). Extracts concentration was estimated using the Qubit® fluorometer and the Qubit® dsDNA HS Assay Kit (ThermoFisher Scientific, Waltham, USA). Sequencing libraries were prepared using either the "NEBNext Ultra II DNA Library Prep Kit for Illumina®" (E7645) or the "NEBNext Ultra II FS DNA Library Prep Kit for Illumina®" (E7805) (New England BioLabs, Ipswich, USA). In the former case, we shared DNA with a Bioruptor® Pico (Diagenode, Seraing, Belgium) prior library preparation. Extracts were diluted to 10 ng/µL and sonicated for eight cycles of 15" sonication and 90" break, in order to obtain fragments of approximately 300-500 bp. In the latter case, enzymatic shearing is combined to the first steps of the library preparation. Fragmentation was carried out for 12' at 37°C in order to obtain DNA fragments of the same length of those of the sonicated samples. For the herbarium collections Hoe5615 and Du33351-15, sharing incubation was shorter, 3' and 10', respectively. In both cases we followed the manufacturer's instructions. At the end of the library preparation procedure, samples were PCR-amplified for 14 cycles, during which samples-specific dual indices ("NEBNext Multiplex Oligos for Illumina®", E7600; New England BioLabs, Ipswich, USA) were added to the fragments. Indexed samples were pooled in equal quantities (four samples per 500 ng pool), dehydrated in a Concentrator Plus (Eppendorf, Hamburg, Germany), and diluted in 7 µL of dd$H_2O$. Each pool was enriched using the custom baits kit and following the manufacturer's protocol. Hybridization took place for 21 h at 65 °C.

Enriched products were PCR-amplified for 14 cycles using the 2X KAPA HiFi HotStart Mix (KAPA Biosystems, Wilmington, USA) and the P7 and P5 adapters as primers. Amplified enriched libraries were purified with 50 µL of AMPure XP Beads (New England BioLabs, Ipswich, USA), following the manufacturer's protocol. Concentrations were measured with the Qubit® fluorometer and fragment length distributions were checked with a Bioanalyzer (Agilent, Santa Clara, USA). In the few cases in which fragment length was different from the desired one (and especially in case short fragments were present), pools were undergone to side selection with the BluePippin (Sage Science, Beverly, USA). All sequencing took place on an Illumina MiSeq System (Illumina Inc., San Diego, USA) at the Transcriptome and Genome Analysis Laboratory (Georg-August-Universität, Gottingen, Germany). Pools were mixed equimolarly and sequenced in two different paired-end runs (6 pools, 24 samples each) with a 2x 250 bp (500 cycles) v2 kit.

**Reads processing und alignment**

We checked the quality of raw reads with FastQC (available at: http://www.bioinformatics.bbsrc.ac.uk/projects/fastqc). Further processing of the raw reads was done using the pipeline HybPhyloMaker (Fér & Schmickl, 2018). Quality-trimmed individual raw reads were mapped to a reference sequence and then merged into contigs that are aligned for each gene separately. As pseudo-reference for read mapping, we used a sequence consisting of the concatenation of the target exonic sequences, separated by stretches of 800 Ns. Sequence adapters were removed, and reads were quality-trimmed using Trimmomatic v. 0.32 (Bolger, Lohse, & Usade, 2014), with the default settings used in HybPhyloMaker. Duplicated reads were removed with FastUniq v. 1.1 (Xu et al., 2012). Mapping to the pseudo-reference genome was done with BWA (Li & Durbin, 2010), and consensus sequences of the mapped reads were produced with ConsensusFixer (available at: https://github.com/cbgethz/ConsensusFixer), since this is the only of the approaches available in HybPhyloMaker able to call ambiguity DNA codes in case of multiple bases per site in the mapped reads. For ConsensusFixer we used the following settings: minimum relative abundance of the alternative base ("plurality" in the setting file of HybPhyloMaker) of 0.2 and a minimum read coverage for ambiguity calling ("mincov") of five.

Consensus sequences were matched to sequences of the target exons to produce PSLX files using BLAT (Kent, 2002). They were therefore combined to produce exon-wise matrices with 'assembled_exons_to_fastas.py' (Weitemier et al., 2014). Matrices were aligned with MAFFT v. 7.029 (Katoh, 2013), using the default program settings, and then gene-wise concatenated with AMAS (Borowiec, 2016). We filtered potential paralogs running the script "HybPhyloMaker4a2_selectNonHet.sh" and setting the maximum number of heterozygous sites per locus ("maxhet" in the HybPhyloMaker settings file) to 10. We continued the pipeline with both datasets, the filtered and unfiltered ones. HybPhyloMaker performs two consecutive steps of missing-data filtering. First, sequences with more than a certain percentage of Ns in an alignment ("missingpercent" in the settings file) are deleted. We set this option to 40. Secondly, alignments with less than a certain percent of sequences ("speciespresence" in the settings file) are filtered out. We set this value to 75, so that alignments with more than 25% of missing sequences (i.e., with less than 25 sequences) were excluded. Finally, we calculated alignments statistics for the selected genes with AMAS, trimAl v. 1.2 (Capella-Gutiérrez, Silla-Martínez, & Gabaldón, 2009), and MstatX (https://github.com/gcollet/MstatX/), as implemented in HybPhyloMaker.

**Phasing**

Since the importance of retrieving allele information for a correct phylogeny estimation (especially in recently diverged group) has been emphasized in recent studies (Eriksson et al.,

2018; Andermann et al., 2019), we tested the impact of allele phasing on our phylogenetic analyses. To avoid the loss of allelic information during the process of allele mapping and consensus sequence production, we took out of the HybPhyloMaker pipeline the .bam files produced after mapping the reads to the pseudo-reference sequence. These were phased with SAMtools v0.1.19 (Li et al., 2009), using the combination of the commands "samtools sort" and "samtools phase". The phased .bam and .bai files were then placed in a new HybPhyloMaker working directory for further processing within the pipeline workflow. A new "/10rawreads/" folder was also placed in the HybPhyloMaker working directory, containing only the file with the modified samples names. The pipeline was therefore resumed for the computing of the consensus sequences (this time allele-wise consensus sequences) by calling the script "HybPhyloMaker2_readmapping.sh" but specifying "mapping=no" in the setting file. The alignments obtained are hereafter called allele alignments. We run the pipeline further as above-explained, with the only differences that (i) matrices of exons belonging to the same gene were not concatenated; and (ii) exon trees (instead of gene trees) were inferred and used for further analyses.

**Position filtering**

We evaluated the effect of filtering alignments for positions that could add phylogenetic noise to the phylogenetic reconstructions, the so-called "phantom" spike positions (ambiguous positions, usually situated close to indel-rich regions of the alignment, with abnormally high substitution rates). We followed the procedure illustrated by Fragoso-Martínez et al. (2017), calculating first the Phylogenetic Informativeness (PI; Townsend, 2007) and net PI profiles using HyPhy (Pond Frost & Muse, 2008) in the web portal PhyDesign (López-Giráldez & Townsend, 2011). As tree inputs, we used trees (different trees for different datasets; e.g., consensus or allele alignments, paralog filtering or not, etc...) obtained from HybPhyloMaker by concatenating gene alignments and FastTree (Price, Dehal & Arkin 2010). The phylograms were then transformed to ultrametric trees using the Penalised Likelihood methods (Sanderson, 2002) as implemented in the R package "ape" (Paradis & Schliep, 2018), with correlated rates and smoothing parameter (λ) set to 1. Relative time scale was set to one at the root.

Secondly, we used the estimated substitution rate per locus from HyPhy and the R script from Fragoso-Martínez et al. (2017) to identify the specific positions in the alignments with an unusually high substitution rate. As substitution rate threshold, we use 20 [the highest value tried in Fragoso-Martínez et al. (2017)] because we were just interested in deleting badly aligned positions and artefacts, without running the risk of excluding naturally highly variable regions in the alignments. Detected sites were checked and removed manually from the alignments. Lists of the removed sites for each scenario are given in the Supporting Information Table S7. The cleaned alignments were then reintroduced in the HybPhyloMaker pipeline, just before the missing-data filtering step.
