## Supplementary material for "Phylogenomics unravels speciation patterns in temperate-montane plant species: a case study on the recently radiating *Ranunculus auricomus* species complex": Table S1

**Table S1** List of targeted loci and information about their function in *Arabidopsis thaliana* (source: The Arabidopsis Information Resource (TAIR); https://www.arabidopsis.org/tools/bulk/genes/index.jsp).

| Locus Identifier | Gene Model Name | Gene Model Description | Gene Model Type | Primary Gene Symbol | All Gene Symbols |
|---|---|---|---|---|---|
| AT1G78800 | AT1G78800.1 | UDP-Glycosyltransferase superfamily protein;(source:Araport11) | protein_coding | | |
| AT5G06830 | AT5G06830.1 | hypothetical protein;(source:Araport11) | protein_coding | | |
| AT2G31740 | AT2G31740.1 | S-adenosyl-L-methionine-dependent methyltransferases superfamily protein;(source:Araport11) | protein_coding | | |
| AT5G11960 | AT5G11960.1 | magnesium transporter, putative (DUF803);(source:Araport11) | protein_coding | | |
| AT4G00560 | AT4G00560.4 | NAD(P)-binding Rossmann-fold superfamily protein;(source:Araport11) | protein_coding | | |
| AT1G80510 | AT1G80510.1 | Encodes a close relative of the amino acid transporter ANT1 (AT3G11900). | protein_coding | | |
| AT2G21250 | AT2G21250.1 | NAD(P)-linked oxidoreductase superfamily protein;(source:Araport11) | protein_coding | | |
| AT5G04420 | AT5G04420.1 | Galactose oxidase/kelch repeat superfamily protein;(source:Araport11) | protein_coding | | |
| AT4G34910 | AT4G34910.1 | P-loop containing nucleoside triphosphate hydrolases superfamily protein;(source:Araport11) | protein_coding | | |
| AT5G66120 | AT5G66120.2 | 3-dehydroquinate synthase;(source:Araport11) | protein_coding | | |
| AT1G45110 | AT1G45110.1 | Tetrapyrrole (Corrin/Porphyrin) Methylase;(source:Araport11) | protein_coding | | |
| AT1G67420 | AT1G67420.2 | Zn-dependent exopeptidases superfamily protein;(source:Araport11) | protein_coding | | |
| AT3G62370 | AT3G62370.1 | heme binding protein;(source:Araport11) | protein_coding | | |
| AT2G19940 | AT2G19940.1 | Putative N-acetyl-gamma-glutamyl-phosphate reductase;(source:Araport11) | protein_coding | | |
| AT1G47670 | AT1G47670.1 | Transmembrane amino acid transporter family protein;(source:Araport11) | protein_coding | | |
| AT5G10910 | AT5G10910.1 | mraW methylase family protein;(source:Araport11) | protein_coding | | |
| AT5G03430 | AT5G03430.1 | phosphoadenosine phosphosulfate (PAPS) reductase family protein;(source:Araport11) | protein_coding | | |
| AT1G67570 | AT1G67570.1 | zinc finger CONSTANS-like protein (DUF3537);(source:Araport11) | protein_coding | | |
| AT3G03890 | AT3G03890.1 | FMN binding protein;(source:Araport11) | protein_coding | | |
| AT4G20325 | AT4G20325.1 | ribonuclease H2 subunit B;(source:Araport11) | protein_coding | | |
| AT4G28020 | AT4G28020.1 | tRNA-thr(GGU) m(6)t(6)A37 methyltransferase;(source:Araport11) | protein_coding | | |
| AT2G23390 | AT2G23390.1 | acyl-CoA;(source:Araport11) | protein_coding | | |
| AT5G61530 | AT5G61530.1 | small G protein family protein / RhoGAP family protein;(source:Araport11) | protein_coding | | |
| AT5G61970 | AT5G61970.1 | signal recognition particle-related / SRP-like protein;(source:Araport11) | protein_coding | | |
| AT3G01060 | AT3G01060.1 | lysine-tRNA ligase;(source:Araport11) | protein_coding | | |
| AT3G04020 | AT3G04020.1 | hypothetical protein;(source:Araport11) | protein_coding | | |
| AT5G01010 | AT5G01010.2 | retinal-binding protein;(source:Araport11) | protein_coding | | |

| | | | |
|---|---|---|---|
| AT3G17430 | AT3G17430.1 | Nucleotide/sugar transporter family protein The mRNA is cell-to-cell mobile. | protein_coding |
| AT2G38000 | AT2G38000.1 | chaperone protein dnaJ-like protein;(source:Araport11) | protein_coding |
| AT4G35850 | AT4G35850.1 | Pentatricopeptide repeat (PPR) superfamily protein;(source:Araport11) | protein_coding |
| AT2G16405 | AT2G16405.1 | Transducin/WD40 repeat-like superfamily protein;(source:Araport11) | protein_coding |
| AT3G20015 | AT3G20015.1 | Eukaryotic aspartyl protease family protein;(source:Araport11) | protein_coding |
| AT3G07180 | AT3G07180.1 | GPI transamidase component PIG-S-like protein;(source:Araport11) | protein_coding |
| AT3G11830 | AT3G11830.1 | TCP-1/cpn60 chaperonin family protein;(source:Araport11) | protein_coding |
| AT1G74240 | AT1G74240.1 | Mitochondrial substrate carrier family protein;(source:Araport11) | protein_coding |
| AT5G64380 | AT5G64380.1 | Inositol monophosphatase family protein;(source:Araport11) | protein_coding |
| AT2G18940 | AT2G18940.1 | Tetratricopeptide repeat (TPR)-like superfamily protein;(source:Araport11) | protein_coding |
| AT3G10210 | AT3G10210.1 | SEC14 cytosolic factor family protein / phosphoglyceride transfer family protein;(source:Araport11) | protein_coding |
| AT3G01380 | AT3G01380.1 | sulfatase and phosphatidylinositolglycan class N domain-containing protein;(source:Araport11) | protein_coding |
| AT1G74580 | AT1G74580.1 | Pentatricopeptide repeat (PPR) superfamily protein;(source:Araport11) | protein_coding |
| AT3G02710 | AT3G02710.1 | Encodes a protein with a putative role in mRNA splicing. | protein_coding |
| AT3G54510 | AT3G54510.2 | Early-responsive to dehydration stress protein (ERD4);(source:Araport11) | protein_coding |
| AT4G34360 | AT4G34360.1 | S-adenosyl-L-methionine-dependent methyltransferases superfamily protein;(source:Araport11) | protein_coding |
| AT4G32050 | AT4G32050.1 | neurochondrin family protein;(source:Araport11) | protein_coding |
| AT5G53070 | AT5G53070.1 | Ribosomal protein L9/RNase H1;(source:Araport11) | protein_coding |
| AT1G77550 | AT1G77550.1 | tubulin-tyrosine ligase;(source:Araport11) | protein_coding |
| AT5G10730 | AT5G10730.1 | NAD(P)-binding Rossmann-fold superfamily protein;(source:Araport11) | protein_coding |
| AT1G04850 | AT1G04850.1 | ubiquitin-associated (UBA)/TS-N domain-containing protein;(source:Araport11) | protein_coding |
| AT4G31790 | AT4G31790.1 | Tetrapyrrole (Corrin/Porphyrin) Methylase;(source:Araport11) | protein_coding |
| AT3G25660 | AT3G25660.1 | Amidase family protein;(source:Araport11) | protein_coding |
| AT2G20360 | AT2G20360.1 | NAD(P)-binding Rossmann-fold superfamily protein;(source:Araport11) | protein_coding |
| AT4G24750 | AT4G24750.1 | Rhodanese/Cell cycle control phosphatase superfamily protein;(source:Araport11) | protein_coding |
| AT3G11210 | AT3G11210.1 | SGNH hydrolase-type esterase superfamily protein;(source:Araport11) | protein_coding |
| AT2G46915 | AT2G46915.1 | DUF3754 family protein, putative (DUF3754);(source:Araport11) | protein_coding |
| AT3G58530 | AT3G58530.1 | RNI-like superfamily protein;(source:Araport11) | protein_coding |
| AT2G01460 | AT2G01460.1 | P-loop containing nucleoside triphosphate hydrolases superfamily protein;(source:Araport11) | protein_coding |

| | | | |
|---|---|---|---|
| AT5G64150 | AT5G64150.1 | RNA methyltransferase family protein;(source:Araport11) | protein_coding |
| AT5G63280 | AT5G63280.1 | C2H2-like zinc finger protein;(source:Araport11) | protein_coding |
| AT1G14300 | AT1G14300.2 | ARM repeat superfamily protein;(source:Araport11) | protein_coding |
| AT4G10130 | AT4G10130.1 | DNAJ heat shock N-terminal domain-containing protein;(source:Araport11) | protein_coding |
| AT5G55960 | AT5G55960.1 | transmembrane protein C9orf5 protein;(source:Araport11) | protein_coding |
| AT1G16650 | AT1G16650.1 | S-adenosyl-L-methionine-dependent methyltransferases superfamily protein;(source:Araport11) | protein_coding |
| AT2G43540 | AT2G43540.1 | transmembrane protein;(source:Araport11) | protein_coding |
| AT4G16330 | AT4G16330.2 | 2-oxoglutarate (2OG) and Fe(II)-dependent oxygenase superfamily protein;(source:Araport11) | protein_coding |
| AT3G04480 | AT3G04480.1 | endoribonuclease;(source:Araport11) | protein_coding |
| AT1G32220 | AT1G32220.1 | NAD(P)-binding Rossmann-fold superfamily protein;(source:Araport11) | protein_coding |
| AT4G15450 | AT4G15450.1 | Senescence/dehydration-associated protein-like protein;(source:Araport11) | protein_coding |
| AT4G16490 | AT4G16490.1 | ARM repeat superfamily protein;(source:Araport11) | protein_coding |
| AT5G27920 | AT5G27920.1 | F-box family protein;(source:Araport11) | protein_coding |
| AT5G62030 | AT5G62030.1 | diphthamide synthesis DPH2 family protein;(source:Araport11) | protein_coding |
| AT3G15520 | AT3G15520.1 | Cyclophilin-like peptidyl-prolyl cis-trans isomerase family protein;(source:Araport11) | protein_coding |
| AT1G74470 | AT1G74470.1 | Encodes for a multifunctional protein with geranylgeranyl reductase activity shown to catalyze the reduction of prenylated geranylgeranyl-chlorophyll a to phytyl-chlorophyll a (chlorophyll a) and free geranylgeranyl pyrophosphate to phytyl pyrophosphate. The mRNA is cell-to-cell mobile. | protein_coding |
| AT2G24230 | AT2G24230.1 | Leucine-rich repeat protein kinase family protein;(source:Araport11) | protein_coding |
| AT5G50310 | AT5G50310.1 | Galactose oxidase/kelch repeat superfamily protein;(source:Araport11) | protein_coding |
| AT3G52210 | AT3G52210.3 | S-adenosyl-L-methionine-dependent methyltransferases superfamily protein;(source:Araport11) | protein_coding |
| AT3G28720 | AT3G28720.1 | transmembrane protein;(source:Araport11) | protein_coding |
| AT5G19150 | AT5G19150.1 | AT5G19150 is a dehydratase that converts (S)-NAD(P)HX to NAD(P)H. | protein_coding |
| AT3G07670 | AT3G07670.1 | Rubisco methyltransferase family protein;(source:Araport11) | protein_coding |
| AT3G21540 | AT3G21540.1 | transducin family protein / WD-40 repeat family protein;(source:Araport11) | protein_coding |
| AT3G03790 | AT3G03790.3 | ankyrin repeat family protein / regulator of chromosome condensation (RCC1) family protein;(source:Araport11) | protein_coding |
| AT3G02300 | AT3G02300.1 | Regulator of chromosome condensation (RCC1) family protein;(source:Araport11) | protein_coding |
| AT4G02820 | AT4G02820.1 | Pentatricopeptide repeat (PPR) superfamily protein;(source:Araport11) | protein_coding |
| AT4G14000 | AT4G14000.1 | Putative methyltransferase family protein;(source:Araport11) | protein_coding |
| AT1G55000 | AT1G55000.1 | peptidoglycan-binding LysM domain-containing protein;(source:Araport11) | protein_coding |

| | | | |
|---|---|---|---|
| AT5G07910 | AT5G07910.1 | Leucine-rich repeat (LRR) family protein;(source:Araport11) | protein_coding |
| AT2G35360 | AT2G35360.1 | ubiquitin family protein;(source:Araport11) | protein_coding |
| AT1G52310 | AT1G52310.1 | protein kinase family protein / C-type lectin domain-containing protein;(source:Araport11) | protein_coding |
| AT1G25570 | AT1G25570.1 | Di-glucose binding protein with Leucine-rich repeat domain-containing protein;(source:Araport11) | protein_coding |
| AT5G04520 | AT5G04520.1 | 3-oxoacyl-acyl-carrier synthase-like protein (Protein of unknown function DUF455);(source:Araport11) | protein_coding |
| AT3G09850 | AT3G09850.1 | D111/G-patch domain-containing protein;(source:Araport11) | protein_coding |
| AT3G17940 | AT3G17940.1 | Galactose mutarotase-like superfamily protein;(source:Araport11) | protein_coding |
| AT4G26980 | AT4G26980.1 | RNI-like superfamily protein;(source:Araport11) | protein_coding |
| AT1G73470 | AT1G73470.3 | hypothetical protein;(source:Araport11) | protein_coding |
| AT5G54910 | AT5G54910.1 | DEA(D/H)-box RNA helicase family protein;(source:Araport11) | protein_coding |
| AT5G59900 | AT5G59900.1 | Pentatricopeptide repeat (PPR) superfamily protein;(source:Araport11) | protein_coding |
| AT5G66530 | AT5G66530.1 | Galactose mutarotase-like superfamily protein;(source:Araport11) | protein_coding |
| AT2G43770 | AT2G43770.1 | Transducin/WD40 repeat-like superfamily protein;(source:Araport11) | protein_coding |
| AT3G07720 | AT3G07720.1 | Galactose oxidase/kelch repeat superfamily protein;(source:Araport11) | protein_coding |
| AT5G10620 | AT5G10620.1 | methyltransferase;(source:Araport11) | protein_coding |
| AT4G01935 | AT4G01935.1 | insulin-induced protein;(source:Araport11) | protein_coding |
| AT2G45990 | AT2G45990.1 | ribosomal RNA small subunit methyltransferase G;(source:Araport11) | protein_coding |
| AT5G61540 | AT5G61540.1 | N-terminal nucleophile aminohydrolases (Ntn hydrolases) superfamily protein;(source:Araport11) | protein_coding |
| AT1G21370 | AT1G21370.1 | transmembrane protein;(source:Araport11) | protein_coding |
| AT3G22290 | AT3G22290.1 | Endoplasmic reticulum vesicle transporter protein;(source:Araport11) | protein_coding |
| AT1G76920 | AT1G76920.1 | F-box family protein;(source:Araport11) | protein_coding |
| AT2G35790 | AT2G35790.1 | transmembrane protein;(source:Araport11) | protein_coding |
| AT1G30300 | AT1G30300.1 | Metallo-hydrolase/oxidoreductase superfamily protein;(source:Araport11) | protein_coding |
| AT5G24970 | AT5G24970.2 | Protein kinase superfamily protein;(source:Araport11) | protein_coding |
| AT3G20230 | AT3G20230.1 | Ribosomal L18p/L5e family protein;(source:Araport11) | protein_coding |
| AT3G44190 | AT3G44190.1 | FAD/NAD(P)-binding oxidoreductase family protein;(source:Araport11) | protein_coding |
| AT1G16180 | AT1G16180.1 | Serinc-domain containing serine and sphingolipid biosynthesis protein;(source:Araport11) | protein_coding |
| AT5G58480 | AT5G58480.1 | O-Glycosyl hydrolases family 17 protein;(source:Araport11) | protein_coding |
| AT5G14140 | AT5G14140.1 | nucleic acid binding / zinc ion binding protein;(source:Araport11) | protein_coding |
| AT3G24030 | AT3G24030.1 | hydroxyethylthiazole kinase family protein;(source:Araport11) | protein_coding |

| | | | |
|---|---|---|---|
| AT1G19600 | AT1G19600.1 | pfkB-like carbohydrate kinase family protein;(source:Araport11) | protein_coding |
| AT4G30310 | AT4G30310.2 | FGGY family of carbohydrate kinase;(source:Araport11) | protein_coding |
| AT4G29590 | AT4G29590.1 | S-adenosyl-L-methionine-dependent methyltransferases superfamily protein;(source:Araport11) | protein_coding |
| AT4G38370 | AT4G38370.1 | Phosphoglycerate mutase family protein;(source:Araport11) | protein_coding |
| AT1G08610 | AT1G08610.1 | Pentatricopeptide repeat (PPR) superfamily protein;(source:Araport11) | protein_coding |
| AT5G48440 | AT5G48440.1 | FAD-dependent oxidoreductase family protein;(source:Araport11) | protein_coding |
| AT5G13520 | AT5G13520.1 | peptidase M1 family protein;(source:Araport11) | protein_coding |
| AT3G24190 | AT3G24190.1 | Protein kinase superfamily protein;(source:Araport11) | protein_coding |
| AT5G19350 | AT5G19350.1 | RNA-binding (RRM/RBD/RNP motifs) family protein;(source:Araport11) | protein_coding |
| AT1G69800 | AT1G69800.2 | Cystathionine beta-synthase (CBS) protein;(source:Araport11) | protein_coding |
| AT5G27950 | AT5G27950.1 | P-loop containing nucleoside triphosphate hydrolases superfamily protein;(source:Araport11) | protein_coding |
| AT4G33945 | AT4G33945.1 | ARM repeat superfamily protein;(source:Araport11) | protein_coding |
| AT5G23590 | AT5G23590.1 | DNAJ heat shock N-terminal domain-containing protein;(source:Araport11) | protein_coding |
| AT1G65030 | AT1G65030.1 | This gene is predicted to encode a protein with a DWD motif. It can bind to DDB1a in Y2H assays, and DDB1b in co-IP assays, and may be involved in the formation of a CUL4-based E3 ubiquitin ligase | protein_coding |
| AT5G15680 | AT5G15680.1 | ARM repeat superfamily protein;(source:Araport11) | protein_coding |
| AT1G63110 | AT1G63110.1 | GPI transamidase subunit PIG-U;(source:Araport11) | protein_coding |
| AT5G65490 | AT5G65490.1 | suppressor-like protein;(source:Araport11) | protein_coding |
| AT5G12040 | AT5G12040.1 | Nitrilase/cyanide hydratase and apolipoprotein N-acyltransferase family protein;(source:Araport11) | protein_coding |
| AT4G16180 | AT4G16180.2 | transmembrane protein;(source:Araport11) | protein_coding |
| AT2G04850 | AT2G04850.1 | Auxin-responsive family protein;(source:Araport11) | protein_coding |
| AT1G03560 | AT1G03560.1 | Pentatricopeptide repeat (PPR-like) superfamily protein;(source:Araport11) | protein_coding |
| AT4G10950 | AT4G10950.1 | SGNH hydrolase-type esterase superfamily protein;(source:Araport11) | protein_coding |
| AT4G08960 | AT4G08960.1 | phosphotyrosyl phosphatase activator (PTPA) family protein;(source:Araport11) | protein_coding |
| AT4G36440 | AT4G36440.1 | G-protein coupled receptor;(source:Araport11) | protein_coding |
| AT1G03150 | AT1G03150.1 | Acyl-CoA N-acyltransferases (NAT) superfamily protein;(source:Araport11) | protein_coding |
| AT5G01470 | AT5G01470.2 | S-adenosyl-L-methionine-dependent methyltransferases superfamily protein;(source:Araport11) | protein_coding |
| AT5G51570 | AT5G51570.1 | SPFH/Band 7/PHB domain-containing membrane-associated protein family;(source:Araport11) | protein_coding |
| AT1G06240 | AT1G06240.1 | diiron containing four-helix bundle family ferritin protein, putative (Protein of unknown function DUF455);(source:Araport11) | protein_coding |

| | | | |
|---|---|---|---|
| AT1G52155 | AT1G52155.1 | transmembrane protein;(source:Araport11) | protein_coding |
| AT1G73350 | AT1G73350.2 | ankyrin repeat protein;(source:Araport11) | protein_coding |
| AT3G02060 | AT3G02060.1 | DEAD/DEAH box helicase;(source:Araport11) | protein_coding |
| AT5G51220 | AT5G51220.1 | ubiquinol-cytochrome C chaperone family protein;(source:Araport11) | protein_coding |
| AT2G21720 | AT2G21720.1 | ArgH (DUF639);(source:Araport11) | protein_coding |
| AT1G05350 | AT1G05350.1 | NAD(P)-binding Rossmann-fold superfamily protein;(source:Araport11) | protein_coding |
| AT1G77030 | AT1G77030.1 | putative DEAD-box ATP-dependent RNA helicase 29;(source:Araport11) | protein_coding |
| AT5G47090 | AT5G47090.1 | coiled-coil protein;(source:Araport11) | protein_coding |
| AT3G19630 | AT3G19630.1 | Radical SAM superfamily protein;(source:Araport11) | protein_coding |
| AT5G15880 | AT5G15880.1 | golgin family A protein;(source:Araport11) | protein_coding |
| AT4G36530 | AT4G36530.2 | alpha/beta-Hydrolases superfamily protein;(source:Araport11) | protein_coding |
| AT5G20660 | AT5G20660.1 | Zn-dependent exopeptidases superfamily protein;(source:Araport11) | protein_coding |
| AT3G54970 | AT3G54970.1 | D-aminoacid aminotransferase-like PLP-dependent enzymes superfamily protein;(source:Araport11) | protein_coding |
| AT5G41330 | AT5G41330.1 | BTB/POZ domain with WD40/YVTN repeat-like protein;(source:Araport11) | protein_coding |
| AT2G31240 | AT2G31240.1 | Tetratricopeptide repeat (TPR)-like superfamily protein;(source:Araport11) | protein_coding |
| AT1G21480 | AT1G21480.1 | Exostosin family protein;(source:Araport11) | protein_coding |
| AT2G25830 | AT2G25830.1 | YebC-like protein;(source:Araport11) | protein_coding |
| AT1G28680 | AT1G28680.1 | HXXXD-type acyl-transferase family protein;(source:Araport11) | protein_coding |
| AT3G05625 | AT3G05625.1 | Tetratricopeptide repeat (TPR)-like superfamily protein;(source:Araport11) | protein_coding |
| AT4G28830 | AT4G28830.1 | S-adenosyl-L-methionine-dependent methyltransferases superfamily protein;(source:Araport11) | protein_coding |
| AT3G06920 | AT3G06920.1 | Tetratricopeptide repeat (TPR)-like superfamily protein;(source:Araport11) | protein_coding |
| AT5G39410 | AT5G39410.1 | Saccharopine dehydrogenase;(source:Araport11) | protein_coding |
| AT1G51730 | AT1G51730.1 | Ubiquitin-conjugating enzyme family protein;(source:Araport11) | protein_coding |
| AT5G65860 | AT5G65860.1 | ankyrin repeat family protein;(source:Araport11) | protein_coding |
| AT3G18860 | AT3G18860.1 | transducin family protein / WD-40 repeat family protein;(source:Araport11) | protein_coding |
| AT5G66005 | AT5G66005.3 | Expressed protein;(source:Araport11) | protein_coding |
| AT2G45500 | AT2G45500.1 | AAA-type ATPase family protein;(source:Araport11) | protein_coding |
| AT1G57770 | AT1G57770.1 | FAD/NAD(P)-binding oxidoreductase family protein;(source:Araport11) | protein_coding |
| AT4G15420 | AT4G15420.1 | Ubiquitin fusion degradation UFD1 family protein;(source:Araport11) | protein_coding |
| AT4G38890 | AT4G38890.1 | FMN-linked oxidoreductases superfamily protein;(source:Araport11) | protein_coding |

| | | | |
|---|---|---|---|
| AT5G12260 | AT5G12260.1 | transferring glycosyl group transferase;(source:Araport11) | protein_coding |
| AT1G74680 | AT1G74680.1 | Exostosin family protein;(source:Araport11) | protein_coding |
| AT1G52630 | AT1G52630.1 | O-fucosyltransferase family protein;(source:Araport11) | protein_coding |
| AT3G04970 | AT3G04970.1 | DHHC-type zinc finger family protein;(source:Araport11) | protein_coding |
| AT5G13400 | AT5G13400.1 | Major facilitator superfamily protein;(source:Araport11) | protein_coding |
| AT2G36630 | AT2G36630.1 | Sulfite exporter TauE/SafE family protein;(source:Araport11) | protein_coding |
| AT3G20790 | AT3G20790.1 | NAD(P)-binding Rossmann-fold superfamily protein;(source:Araport11) | protein_coding |
| AT4G33440 | AT4G33440.1 | Pectin lyase-like superfamily protein;(source:Araport11) | protein_coding |
| AT4G01040 | AT4G01040.1 | Glycosyl hydrolase superfamily protein;(source:Araport11) | protein_coding |
| AT5G19570 | AT5G19570.1 | transmembrane protein;(source:Araport11) | protein_coding |
| AT2G36360 | AT2G36360.5 | Galactose oxidase/kelch repeat superfamily protein;(source:Araport11) | protein_coding |
| AT3G12940 | AT3G12940.1 | 2-oxoglutarate (2OG) and Fe(II)-dependent oxygenase superfamily protein;(source:Araport11) | protein_coding |
| AT2G39670 | AT2G39670.2 | Radical SAM superfamily protein;(source:Araport11) | protein_coding |
| AT5G67140 | AT5G67140.1 | F-box/RNI-like superfamily protein;(source:Araport11) | protein_coding |
| AT3G05410 | AT3G05410.2 | Photosystem II reaction center PsbP family protein;(source:Araport11) | protein_coding |
| AT1G09280 | AT1G09280.1 | rhodanese-like domain protein;(source:Araport11) | protein_coding |
| AT1G55880 | AT1G55880.1 | Pyridoxal-5-phosphate-dependent enzyme family protein;(source:Araport11) | protein_coding |
| AT2G21960 | AT2G21960.1 | transmembrane protein;(source:Araport11) | protein_coding |
| AT2G03430 | AT2G03430.1 | Ankyrin repeat family protein;(source:Araport11) | protein_coding |
| AT1G74640 | AT1G74640.1 | alpha/beta-Hydrolases superfamily protein;(source:Araport11) | protein_coding |
| AT1G01770 | AT1G01770.1 | propionyl-CoA carboxylase;(source:Araport11) | protein_coding |
| AT5G10920 | AT5G10920.1 | L-Aspartase-like family protein;(source:Araport11) | protein_coding |
| AT1G08125 | AT1G08125.2 | S-adenosyl-L-methionine-dependent methyltransferases superfamily protein;(source:Araport11) | protein_coding |
| AT5G47860 | AT5G47860.1 | Gut esterase (DUF1350);(source:Araport11) | protein_coding |
| AT3G03440 | AT3G03440.1 | ARM repeat superfamily protein;(source:Araport11) | protein_coding |
| AT3G47610 | AT3G47610.1 | transcription regulator/ zinc ion binding protein;(source:Araport11) | protein_coding |
| AT4G29310 | AT4G29310.1 | DUF1005 family protein (DUF1005);(source:Araport11) | protein_coding |
| AT5G10460 | AT5G10460.1 | Haloacid dehalogenase-like hydrolase (HAD) superfamily protein;(source:Araport11) | protein_coding |
| AT1G04900 | AT1G04900.1 | NADH dehydrogenase ubiquinone complex I, assembly factor-like protein (DUF185);(source:Araport11) | protein_coding |
| AT4G19010 | AT4G19010.1 | Encodes for a 4-coumarate-CoA ligase involved in the biosynthesis of the benzenoid ring of ubiquinone from phenylalanine. | protein_coding |
| AT1G75210 | AT1G75210.1 | HAD-superfamily hydrolase, subfamily IG, 5-nucleotidase;(source:Araport11) | protein_coding |

| | | | |
|---|---|---|---|
| AT4G06676 | AT4G06676.1 | etoposide-induced protein;(source:Araport11) | protein_coding |
| AT5G23340 | AT5G23340.1 | RNI-like superfamily protein;(source:Araport11) | protein_coding |
| AT3G16840 | AT3G16840.1 | P-loop containing nucleoside triphosphate hydrolases superfamily protein;(source:Araport11) | protein_coding |
| AT5G56900 | AT5G56900.2 | CwfJ-like family protein / zinc finger (CCCH-type) family protein;(source:Araport11) | protein_coding |
| AT3G43540 | AT3G43540.1 | initiation factor 4F subunit (DUF1350);(source:Araport11) | protein_coding |
| AT2G20790 | AT2G20790.1 | clathrin adaptor complexes medium subunit family protein;(source:Araport11) | protein_coding |
| AT4G02485 | AT4G02485.1 | 2-oxoglutarate (2OG) and Fe(II)-dependent oxygenase superfamily protein;(source:Araport11) | protein_coding |
| AT1G72090 | AT1G72090.1 | Methylthiotransferase;(source:Araport11) | protein_coding |
| AT4G01130 | AT4G01130.1 | GDSL-motif esterase/acyltransferase/lipase. Enzyme group with broad substrate specificity that may catalyze acyltransfer or hydrolase reactions with lipid and non-lipid substrates. | protein_coding |
| AT2G44970 | AT2G44970.1 | alpha/beta-Hydrolases superfamily protein;(source:Araport11) | protein_coding |
| AT2G42450 | AT2G42450.1 | alpha/beta-Hydrolases superfamily protein;(source:Araport11) | protein_coding |
| AT3G58470 | AT3G58470.1 | nucleic acid binding / methyltransferase;(source:Araport11) | protein_coding |
| AT5G11280 | AT5G11280.1 | tail fiber;(source:Araport11) | protein_coding |
| AT1G32160 | AT1G32160.1 | beta-casein (DUF760);(source:Araport11) | protein_coding |
| AT2G17670 | AT2G17670.1 | Tetratricopeptide repeat (TPR)-like superfamily protein;(source:Araport11) | protein_coding |
| AT4G23440 | AT4G23440.1 | Disease resistance protein (TIR-NBS class);(source:Araport11) | protein_coding |
| AT3G06950 | AT3G06950.1 | Pseudouridine synthase family protein;(source:Araport11) | protein_coding |
| AT5G35560 | AT5G35560.1 | DENN (AEX-3) domain-containing protein;(source:Araport11) | protein_coding |
| AT4G36390 | AT4G36390.1 | Methylthiotransferase;(source:Araport11) | protein_coding |
| AT5G19850 | AT5G19850.1 | alpha/beta-Hydrolases superfamily protein;(source:Araport11) | protein_coding |
| AT4G21520 | AT4G21520.1 | Transducin/WD40 repeat-like superfamily protein;(source:Araport11) | protein_coding |
| AT5G17410 | AT5G17410.2 | Spc97 / Spc98 family of spindle pole body (SBP) component;(source:Araport11) | protein_coding |
| AT2G30700 | AT2G30700.1 | GPI-anchored protein;(source:Araport11) | protein_coding |
| AT5G19680 | AT5G19680.1 | Leucine-rich repeat (LRR) family protein;(source:Araport11) | protein_coding |
| AT3G55070 | AT3G55070.1 | LisH/CRA/RING-U-box domains-containing protein;(source:Araport11) | protein_coding |
| AT1G54310 | AT1G54310.2 | S-adenosyl-L-methionine-dependent methyltransferases superfamily protein;(source:Araport11) | protein_coding |
| AT1G03250 | AT1G03250.2 | R3H domain protein;(source:Araport11) | protein_coding |
| AT1G53760 | AT1G53760.1 | K+-H+ exchange-like protein;(source:Araport11) | protein_coding |
| AT4G17100 | AT4G17100.2 | poly(U)-specific endoribonuclease-B protein;(source:Araport11) | protein_coding |

| | | | |
|---|---|---|---|
| AT5G24760 | AT5G24760.1 | GroES-like zinc-binding dehydrogenase family protein;(source:Araport11) | protein_coding |
| AT1G03687 | AT1G03687.1 | DTW domain-containing protein;(source:Araport11) | protein_coding |
| AT3G24730 | AT3G24730.1 | mRNA splicing factor, thioredoxin-like U5 snRNP;(source:Araport11) | protein_coding |
| AT5G17530 | AT5G17530.3 | phosphoglucosamine mutase family protein;(source:Araport11) | protein_coding |
| AT5G45760 | AT5G45760.1 | Transducin/WD40 repeat-like superfamily protein;(source:Araport11) | protein_coding |
| AT1G04420 | AT1G04420.1 | NAD(P)-linked oxidoreductase superfamily protein;(source:Araport11) | protein_coding |
| AT4G24880 | AT4G24880.1 | snurportin-1 protein;(source:Araport11) | protein_coding |
| AT3G52390 | AT3G52390.2 | TatD related DNase;(source:Araport11) | protein_coding |
| AT4G01995 | AT4G01995.1 | beta-carotene isomerase D27;(source:Araport11) | protein_coding |
| AT3G15410 | AT3G15410.2 | Leucine-rich repeat (LRR) family protein;(source:Araport11) | protein_coding |
| AT3G60810 | AT3G60810.1 | DUF1499 family protein;(source:Araport11) | protein_coding |
| AT5G13890 | AT5G13890.1 | plant viral-response family protein (DUF716);(source:Araport11) | protein_coding |
| AT3G48380 | AT3G48380.3 | Peptidase C78, ubiquitin fold modifier-specific peptidase 1/ 2;(source:Araport11) | protein_coding |
| AT2G47330 | AT2G47330.1 | P-loop containing nucleoside triphosphate hydrolases superfamily protein;(source:Araport11) | protein_coding |
| AT1G78280 | AT1G78280.1 | transferases, transferring glycosyl groups;(source:Araport11) | protein_coding |
| AT1G06050 | AT1G06050.1 | ENHANCED DISEASE RESISTANCE-like protein (DUF1336);(source:Araport11) | protein_coding |
| AT3G14910 | AT3G14910.1 | Rab3 GTPase-activating protein non-catalytic subunit;(source:Araport11) | protein_coding |
| AT2G40570 | AT2G40570.1 | initiator tRNA phosphoribosyl transferase family protein;(source:Araport11) | protein_coding |
| AT4G26240 | AT4G26240.1 | histone-lysine N-methyltransferase;(source:Araport11) | protein_coding |
| AT3G57790 | AT3G57790.1 | Pectin lyase-like superfamily protein;(source:Araport11) | protein_coding |
| AT3G16565 | AT3G16565.2 | threonyl and alanyl tRNA synthetase second additional domain-containing protein;(source:Araport11) | protein_coding |
| AT1G73180 | AT1G73180.1 | Eukaryotic translation initiation factor eIF2A family protein;(source:Araport11) | protein_coding |
| AT1G43860 | AT1G43860.1 | sequence-specific DNA binding transcription factor;(source:Araport11) | protein_coding |
| AT4G19860 | AT4G19860.1 | Encodes a cytosolic calcium-independent phospholipase A. | protein_coding |
| AT3G01920 | AT3G01920.1 | DHBP synthase RibB-like alpha/beta domain-containing protein;(source:Araport11) | protein_coding |
| AT2G28790 | AT2G28790.1 | Pathogenesis-related thaumatin superfamily protein;(source:Araport11) | protein_coding |
| AT3G12150 | AT3G12150.1 | alpha/beta hydrolase family protein;(source:Araport11) | protein_coding |
| AT2G23890 | AT2G23890.1 | HAD-superfamily hydrolase, subfamily IG, 5-nucleotidase;(source:Araport11) | protein_coding |
| AT3G10970 | AT3G10970.1 | Haloacid dehalogenase-like hydrolase (HAD) superfamily protein;(source:Araport11) | protein_coding |
| AT3G14075 | AT3G14075.1 | Mono-/di-acylglycerol lipase, N-terminal;(source:Araport11) | protein_coding |

| | | | |
|---|---|---|---|
| AT4G04940 | AT4G04940.1 | transducin family protein / WD-40 repeat family protein;(source:Araport11) | protein_coding |
| AT5G21070 | AT5G21070.1 | Fe(3+) dicitrate transport system permease;(source:Araport11) | protein_coding |
| AT2G37500 | AT2G37500.1 | arginine biosynthesis protein ArgJ family;(source:Araport11) | protein_coding |
| AT2G23540 | AT2G23540.1 | GDSL-motif esterase/acyltransferase/lipase. Enzyme group with broad substrate specificity that may catalyze acyltransfer or hydrolase reactions with lipid and non-lipid substrates. | protein_coding |
| AT3G04560 | AT3G04560.1 | nucleolar/coiled-body phosphoprotein;(source:Araport11) | protein_coding |
| AT2G25280 | AT2G25280.1 | AmmeMemoRadiSam system protein B;(source:Araport11) | protein_coding |
| AT1G07040 | AT1G07040.1 | plant/protein;(source:Araport11) | protein_coding |
| AT3G05510 | AT3G05510.1 | Phospholipid/glycerol acyltransferase family protein;(source:Araport11) | protein_coding |
| AT3G28700 | AT3G28700.1 | NADH dehydrogenase ubiquinone complex I, assembly factor-like protein (DUF185);(source:Araport11) | protein_coding |
| AT5G14240 | AT5G14240.1 | Thioredoxin superfamily protein;(source:Araport11) | protein_coding |
| AT1G21780 | AT1G21780.1 | BTB/POZ domain-containing protein. Contains similarity to gb:AJ000644 SPOP (speckle-type POZ protein) from Homo sapiens and contains a PF:00651 BTB/POZ domain. ESTs gb:T75841, gb:R89974, gb:R30221, gb:N96386, gb:T76457, gb:AI100013 and gb:T76456 come from this gene;supported by full-length. Interacts with CUL3A and CUL3B. | protein_coding |
| AT5G14550 | AT5G14550.1 | Core-2/I-branching beta-1,6-N-acetylglucosaminyltransferase family protein;(source:Araport11) | protein_coding |
| AT4G04320 | AT4G04320.1 | malonyl-CoA decarboxylase family protein;(source:Araport11) | protein_coding |
| AT5G45170 | AT5G45170.1 | Haloacid dehalogenase-like hydrolase (HAD) superfamily protein;(source:Araport11) | protein_coding |
| AT1G03030 | AT1G03030.1 | P-loop containing nucleoside triphosphate hydrolases superfamily protein;(source:Araport11) | protein_coding |
| AT4G17760 | AT4G17760.1 | PCNA domain-containing protein;(source:Araport11) | protein_coding |
| AT5G66290 | AT5G66290.1 | hypothetical protein;(source:Araport11) | protein_coding |
| AT2G15860 | AT2G15860.2 | BAT2 domain protein;(source:Araport11) | protein_coding |
| AT1G50510 | AT1G50510.1 | indigoidine synthase A family protein;(source:Araport11) | protein_coding |
| AT3G50685 | AT3G50685.1 | anti-muellerian hormone type-2 receptor;(source:Araport11) | protein_coding |
| AT5G05200 | AT5G05200.1 | Protein kinase superfamily protein;(source:Araport11) | protein_coding |
| AT5G02230 | AT5G02230.1 | Haloacid dehalogenase-like hydrolase (HAD) superfamily protein;(source:Araport11) | protein_coding |
| AT2G33630 | AT2G33630.1 | NAD(P)-binding Rossmann-fold superfamily protein;(source:Araport11) | protein_coding |
| AT3G52570 | AT3G52570.1 | alpha/beta-Hydrolases superfamily protein;(source:Araport11) | protein_coding |
| AT5G19630 | AT5G19630.1 | alpha/beta-Hydrolases superfamily protein;(source:Araport11) | protein_coding |
| AT1G65270 | AT1G65270.2 | ER membrane protein complex subunit-like protein;(source:Araport11) | protein_coding |

| | | | | | |
|---|---|---|---|---|---|
| AT2G01070 | AT2G01070.1 | Lung seven transmembrane receptor family protein;(source:Araport11) | protein_coding | | |
| AT5G04910 | AT5G04910.1 | DNA repair REX1-B protein;(source:Araport11) | protein_coding | | |
| AT2G27680 | AT2G27680.1 | NAD(P)-linked oxidoreductase superfamily protein;(source:Araport11) | protein_coding | | |
| AT4G34500 | AT4G34500.1 | Protein kinase superfamily protein;(source:Araport11) | protein_coding | | |
| AT5G39900 | AT5G39900.1 | Small GTP-binding protein;(source:Araport11) | protein_coding | | |
| AT3G08010 | AT3G08010.1 | Encodes a chloroplast-localized protein ATAB2. ATAB2 is involved in the biogenesis of Photosystem I and II. ATAB2 has A/U-rich RNA-binding activity and presumably functions as an activator of translation with targets at PS I and PS II. | protein_coding | (ATAB2) | (ATAB2) |
| AT2G39190 | AT2G39190.2 | member of ATH subfamily | protein_coding | (ATATH8) | (ATATH8) |
| AT3G08970 | AT3G08970.1 | J domain protein localized in ER lumen. Can compensate for the growth defect in jem1 scj1 mutant yeast. Also shows similarity to HSP40 proteins and is induced by heat stress. At high temperatures, mutant alleles are not transmitted through the pollen due to defects in pollen tube growth. | protein_coding | (ATERDJ3A) | (ATERDJ3A);THERMOSENSITIVE MALE STERILE 1 (TMS1) |
| AT4G29170 | AT4G29170.1 | A homolog of yeast, mouse and human mnd1delta protein. Null mutants exhibit normal vegetative and flower development; however, during prophase I, chromosomes become fragmented resulting in random distribution of the fragments between polyads. Both male and female meiosis are defective and strong accumulation of AtRAD51 was observed in the inflorescence nuclei of mutant plants. Similarly to its yeast and animal homologues, AtMnd1 might play a role in DSB repair during meiosis. | protein_coding | (ATMND1) | (ATMND1) |
| AT3G01720 | AT3G01720.1 | peptidyl serine alpha-galactosyltransferase;(source:Araport11) | protein_coding | (ATSERGT1) | (ATSERGT1) |
| AT4G14790 | AT4G14790.1 | encodes a nuclear-encoded DExH box RNA helicase, which is localized to mitochondria and whose in vitro ATPase activity is stimulated with mitochondrial RNA. | protein_coding | (ATSUV3) | EMBRYO SAC DEVELOPMENT ARREST 15 (EDA15);SUPPRESSOR OF VAR 3 (SUV3); (ATSUV3) |
| AT2G16860 | AT2G16860.1 | GCIP-interacting family protein;(source:Araport11) | protein_coding | (ATSYF2) | (ATSYF2) |
| AT3G54860 | AT3G54860.2 | Homologous to yeast VPS33. Forms a complex with VCL1 and AtVPS11. Involved in vacuolar biogenesis. | protein_coding | (ATVPS33) | VACUOLAR PROTEIN SORTING 33 (VPS33); (ATVPS33) |
| AT5G20990 | AT5G20990.1 | Involved in molybdenum cofactor (Moco) biosynthesis, inserting Mo into Molybdopterin. sir loss-of-function mutants are resistant to sirtinol, a modulator of auxin signaling. | protein_coding | (B73) | (B73);SIRTINOL 4 (SIR4);CO-FACTOR FOR NITRATE REDUCTASE AND XANTHINE DEHYDROGENASE 1 (CNX1);CHLORATE RESISTANT 6 (CHL6);CO-FACTOR FOR NITRATE REDUCTASE AND XANTHINE DEHYDROGENASE (CNX) |
| AT3G12300 | AT3G12300.1 | Similar to Bug22p in Paramecium, a conserved centrosomal/ciliary protein. This protein is widespread in eukaryotes harboring centrioles/cilia at some stage of their life cycles. Among eukaryotes devoid of centrioles/cilia, plants possess BUG22 genes whereas some fungi (at least ascomycetes) do not. | protein_coding | (BUG22) | (BUG22); (ATBUG22) |
| AT1G23400 | AT1G23400.1 | Promotes the splicing of chloroplast group II introns. | protein_coding | (CAF2) | (CAF2);ARABIDOPSIS THALIANA HOMOLOG OF MAIZE CAF2 (ATCAF2) |
| AT5G57300 | AT5G57300.1 | S-adenosyl-L-methionine-dependent methyltransferases superfamily protein;(source:Araport11) | protein_coding | (COQ5) | (COQ5) |
| AT1G17760 | AT1G17760.1 | Encodes a homolog of the mammalian protein CstF77, a polyadenylation factor subunit. RNA 3′-end?processing factor of antisense FLC transcript. Mediates | protein_coding | (CSTF77) | ARABIDOPSIS THALIANA CLEAVAGE STIMULATION FACTOR 77 (ATCSTF77); (CSTF77) |

| | | | | | |
|---|---|---|---|---|---|
| | | silencing of the floral repressor gene FLC. Member of CstF complex. | | | |
| AT5G54290 | AT5G54290.2 | Encodes CcdA, a thylakoid membrane protein required for the transfer of reducing equivalents from stroma to thylakoid lumen. | protein_coding | (CcdA) | (CcdA) |
| AT4G39520 | AT4G39520.1 | Encodes a member of the DRG (developmentally regulated G-protein) family. Has GTPase activity. | protein_coding | (DRG1-1) | (DRG1-1) |
| AT5G11480 | AT5G11480.1 | P-loop containing nucleoside triphosphate hydrolases superfamily protein;(source:Araport11) | protein_coding | (ENGB-2) | (ENGB-2) |
| AT3G51820 | AT3G51820.1 | Encodes a protein with chlorophyll synthase activity. This enzyme has been shown to perform the esterification of chlorophyllide (a and b), the last step of chlorophyll biosynthesis. Although it can use either geranylgeranyl pyrophosphate (GGPP) or phytyl pyrophosphate (PhyPP) as substrates, the esterification reaction was faster with GGPP than with PhyPP. | protein_coding | (G4) | (ATG4);PIGMENT DEFECTIVE 325 (PDE325); (CHLG); (G4) |
| AT3G10850 | AT3G10850.1 | glyoxalase II cytoplasmic isozyme (Glx2-2) mRNA, complete | protein_coding | (GLY2) | GLYOXALASE 2-2 (GLX2-2); (GLY2) |
| AT5G08720 | AT5G08720.1 | Encodes PIN2 PROMOTER BINDING PROTEIN 1 (PPP1), an evolutionary conserved plant-specific DNA binding protein that acts on transcription of PIN genes. Also named as HCF145. Mutations in HCF145 have reduced level of the tricistronic psaA-psaB-rps (small-subunit ribosomal protein)14 mRNA which encodes for the major subunits of the photosystem I (PSI). HCF145 binds to the 5'UTR of PSAA via a novel TMR domain. It functions to stabilize the PSAA transcript. | protein_coding | (HCF145) | (HCF145);PIN2 PROMOTER BINDING PROTEIN 1 (PPP1);HIGH CHLOROPHYLL FLUORESCENCE 145 (HCF145) |
| AT1G69740 | AT1G69740.1 | Encodes a putative 5-aminolevulinate dehydratase involved in chlorophyll biosynthesis. | protein_coding | (HEMB1) | 5-AMINOLEVULINIC ACID DEHYDRATASE 1 (ALAD1); (HEMB1) |
| AT5G59250 | AT5G59250.1 | Encodes a chloroplast localized H+/glucose antiporter. | protein_coding | (HP59) | PLASTIDIC SUGAR TRANSPORTER (PSUT); (HP59) |
| AT1G48050 | AT1G48050.1 | Ku80 and ku70 form the heterodimer complex Ku, required for proper maintenance of the telomeric C strand. Ku regulates the extension of the telomeric G strand. Interacts with WEX, and this interaction stimulates the WEX exonuclease activity. Binds double stranded DNA breaks as a heterodimer with Ku70, involved in non-homologous end joining repair. Mutants are defective in T-DNA integration. Over expression confers increased resistance to DNA damage agents and increased susceptibility to T-DNA transformation. | protein_coding | (KU80) | ARABIDOPSIS THALIANA KU80 HOMOLOG (ATKU80); (KU80) |
| AT3G52200 | AT3G52200.2 | Encodes a dihydrolipoamide S-acetyltransferase, a subunit of the mitochondrial pyruvate dehydrogenase complex. | protein_coding | (LTA3) | MITOCHONDRIAL PYRUVATE DEHYDROGENASE SUBUNIT 2-1 (MTE2-1); (LTA3) |
| AT1G32080 | AT1G32080.1 | Encodes a plant LrgAB/CidAB protein localized to the chloroplast envelope that is involved in chloroplast development, carbon partitioning, ABA/drought response, and leaf senescence. The gene may have evolved from gene fusion of bacterial lrgA and lrgB. | protein_coding | (LrgB) | (LrgB); (PLGG1); (PLGG); (AtLrgB) |
| AT1G18680 | AT1G18680.1 | HNH endonuclease domain-containing protein;(source:Araport11) | protein_coding | (M20) | (M20) |
| AT1G11090 | AT1G11090.1 | alpha/beta-Hydrolases superfamily protein;(source:Araport11) | protein_coding | (MAGL1) | (MAGL1) |
| AT5G11650 | AT5G11650.1 | alpha/beta-Hydrolases superfamily protein;(source:Araport11) | protein_coding | (MAGL13) | (MAGL13) |
| AT5G16120 | AT5G16120.2 | alpha/beta-Hydrolases superfamily protein;(source:Araport11) | protein_coding | (MAGL15) | (MAGL15) |

| | | | | | |
|---|---|---|---|---|---|
| AT1G03090 | AT1G03090.2 | MCCA is the biotinylated subunit of the dimer MCCase, which is involved in leucine degradation. Both subunits are nuclear coded and the active enzyme is located in the mitochondrion. | protein_coding | (MCCA) | (MCCA) |
| AT5G02130 | AT5G02130.1 | SSR1 encodes a tetratricopeptide repeat- containing protein localized in mitochondria. It is involved in root development, possibly by through effects on auxin transport. In ssr1 mutants, the expression PIN genes and trafficking of PIN2 is altered which in turn affects distribution of auxin in the roots. | protein_coding | (NDP1) | SHORT AND SWOLLEN ROOT 1 (SSR1); (NDP1) |
| AT4G08790 | AT4G08790.1 | NIT1 amidase involved in the breakdown of deaminated glutathione (dGSH). It is active towards dGSH and dOA with a preference for dGSH. | protein_coding | (NIT1) | (NIT1) |
| AT1G06560 | AT1G06560.1 | NOL1/NOP2/sun family protein;(source:Araport11) | protein_coding | (NOP2C) | (NOP2C); (ATTRM4F);TRNA METHYLTRANSFERASE 4F (TRM4F) |
| AT2G15430 | AT2G15430.1 | Non-catalytic subunit of nuclear DNA-dependent RNA polymerases II, IV and V; homologous to budding yeast RPB3 and the E. coli RNA polymerase alpha subunit. A closely related paralog, encoded by At2g15400, can substitute for At2g15430 in the context of Pol V. | protein_coding | (NRPB3) | (NRPD3); (NRPE3A); (NRPB3); (RPB35.5A); (RBP36A) |
| AT4G17300 | AT4G17300.1 | Asparaginyl-tRNA synthetase protein involved in amino acid activation/protein synthesis. | protein_coding | (NS1) | (NS1); (ATNS1);OVULE ABORTION 8 (OVA8) |
| AT1G07615 | AT1G07615.1 | GTP-binding protein Obg/CgtA;(source:Araport11) | protein_coding | (OBG A-1) | (OBG A-1) |
| AT3G44160 | AT3G44160.1 | Encodes a chloroplast-localized Omp85 family member. Members of this family chaperone the membrane insertion of beta-barrel-shaped outer membrane proteins in bacteria, mitochondria and probably chloroplasts and facilitate the transfer of nuclear-encoded cytosolically synthesized preproteins across the outer envelope of chloroplasts. | protein_coding | (P39) | (P39) |
| AT2G39550 | AT2G39550.1 | encodes the beta subunit of geranylgeranyl transferase (GGT-IB), involved in both ABA-mediated and auxin signaling pathways. | protein_coding | (PGGT-I) | GERANYLGERANYLTRANSFERASE-I BETA SUBUNIT (ATGGT-IB); (PGGT-I); (GGB) |
| AT5G41880 | AT5G41880.1 | DNA primase POLA3;(source:Araport11) | protein_coding | (POLA3) | (POLA4); (POLA3) |
| AT4G01690 | AT4G01690.1 | Encodes protoporphyrinogen oxidase (PPOX). | protein_coding | (PPOX) | (PPOX); (PPO1); (HEMG1) |
| AT3G12530 | AT3G12530.1 | PSF2;(source:Araport11) | protein_coding | (PSF2) | (PSF2) |
| AT3G61620 | AT3G61620.1 | exonuclease RRP41 (RRP41) | protein_coding | (RRP41) | (RRP41) |
| AT5G51340 | AT5G51340.1 | SCC4 is a tetratricopeptide repeat containing protein and a likely component of a plant cohesion loading complex along with its partner SSC2 It is expressed primarily in dividing cells. Loss of function mutants are embryo lethal, arresting by globular stage. | protein_coding | (SCC4) | (SCC4) |
| AT1G64350 | AT1G64350.1 | seh1-like protein | protein_coding | (SEH1H) | (SEH1H) |
| AT3G54690 | AT3G54690.1 | Sugar isomerase (SIS) family protein;(source:Araport11) | protein_coding | (SETH3) | (SETH3) |
| AT2G21280 | AT2G21280.1 | A nuclear-encoded, plastid-targeted protein (AtSulA) whose overexpression causes severe yet stochastic plastid (shown in chloroplasts and leucoplasts) division defects. The protein does not appear to interact with either AtFtsZ proteins when studied in a yeast two-hybrid system. | protein_coding | (SULA) | GIANT CHLOROPLAST 1 (GC1); (ATSULA); (SULA) |
| AT5G09860 | AT5G09860.1 | Encodes a component of the putative Arabidopsis THO/TREX complex: THO1 or HPR1 (At5g09860), THO2 (At1g24706), THO3 or TEX1 (At5g56130), THO5 (At5g42920, At1g45233), THO6 (At2g19430), and THO7 | protein_coding | (THO1) | (AtTHO1); (HPR1); (AtHPR1); (THO1) |

| | | | | | |
|---|---|---|---|---|---|
| | | (At5g16790, At3g02950). THO/TREX complexes in animals have been implicated in the transport of mRNA precursors. Mutants of THO3/TEX1, THO1, THO6 accumulate reduced amount of small interfering (si)RNA, suggesting a role of the putative Arabidopsis THO/TREX in siRNA biosynthesis. One of the pathways affected by THO1 is the miRNA399-PHO2 pathway that regulates root APase activity. | | | |
| AT5G55220 | AT5G55220.1 | trigger factor type chaperone family protein;(source:Araport11) | protein_coding | (TIG1) | (TIG1) |
| AT4G39820 | AT4G39820.1 | Part of multi-protein complex, acting as guanine nucleotide exchange factors (GEFs) and possibly as tethers, regulating intracellular trafficking. | protein_coding | (TRAPPC12) | (TRAPPC12) |
| AT1G78010 | AT1G78010.1 | tRNA modification GTPase;(source:Araport11) | protein_coding | (TRME) | (TRME) |
| AT3G56740 | AT3G56740.1 | ATI3A interacting protein containing a large N-terminal rhomboid-like transmembrane domain and a UBA domain at their C terminus, localized in the ER with an important role in plant heat tolerance. UBAC2 proteins may act as both cargo receptors and inducers of an ATI3-mediated selective autophagy pathway, where ATI3 and UBAC2 proteins are delivered to the vacuole under ER stress in an autophagy-dependent manner. | protein_coding | (UBAC2A) | (UBAC2A) |
| AT1G43620 | AT1G43620.1 | Encodes a UDP-glucose:sterol-glucosyltransferase. Mutants produce pale greenish-brown seeds whose dormancy was slightly reduced | protein_coding | (UGT80B1) | TRANSPARENT TESTA 15 (TT15);TRANSPARENT TESTA GLABROUS 15 (TTG15); (UGT80B1) |
| AT5G44560 | AT5G44560.1 | SNF7 family protein;(source:Araport11) | protein_coding | (VPS2.2) | (VPS2.2) |
| AT4G19490 | AT4G19490.1 | Putative homolog of yeast Vps54. Thought to associate with POK and ATVPS53 in a plant GARP-like complex involved in the membrane trafficking system. | protein_coding | (VPS54) | (VPS54);ARABIDOPSIS THALIANA VPS54 HOMOLOG (ATVPS54) |
| AT5G14220 | AT5G14220.1 | Encodes PPO2, a putative protoporphyrinogen oxidase based on sequence homology. Also known as MEE61 (maternal effect embryo arrest 61). mee61 mutant shows arrested endosperm development. | protein_coding | (hemg2) | (PPO2);MATERNAL EFFECT EMBRYO ARREST 61 (MEE61); (hemg2) |
| AT5G55500 | AT5G55500.1 | Encodes a beta-1,2-xylosyltransferase that is glycosylated at two positions. The mRNA is cell-to-cell mobile. | protein_coding | "BETA-1,2-XYLOSYLTRANSFERASE" (XYLT) | ARABIDOPSIS THALIANA BETA-1,2-XYLOSYLTRANSFERASE (ATXYLT);"BETA-1,2-XYLOSYLTRANSFERASE" (XYLT) |
| AT1G01280 | AT1G01280.1 | member of CYP703A CYP703A2 is expressed specifically in anthers of land plants, catalyzing the in-chain hydroxylation at the C-7 position of medium-chain saturated fatty acids (lauric acid in-chain hydroxylase) which is involved in pollen development (sporopollenin synthesis). | protein_coding | "CYTOCHROME P450, FAMILY 703, SUBFAMILY A, POLYPEPTIDE 2" (CYP703A2) | (CYP703);"CYTOCHROME P450, FAMILY 703, SUBFAMILY A, POLYPEPTIDE 2" (CYP703A2) |
| AT1G69500 | AT1G69500.1 | Encodes a cytochrome P450, designated CYP704B1. Expressed in the developing anthers. Essential for pollen exine development. Mutations in CYP704B1 result in impaired pollen walls that lack a normal exine layer and exhibit a characteristic striped surface, termed zebra phenotype. Heterologous expression of CYP704B1 in yeast cells demonstrated that it catalyzes omega-hydroxylation of long-chain fatty acids, implicating these molecules in sporopollenin synthesis. | protein_coding | "CYTOCHROME P450, FAMILY 704, SUBFAMILY B, POLYPEPTIDE 1" (CYP704B1) | "CYTOCHROME P450, FAMILY 704, SUBFAMILY B, POLYPEPTIDE 1" (CYP704B1) |
| AT1G31800 | AT1G31800.1 | Encodes a protein with β-ring carotenoid hydroxylase activity. The mRNA is cell-to-cell mobile. | protein_coding | "CYTOCHROME P450, FAMILY 97, SUBFAMILY A, POLYPEPTIDE 3" (CYP97A3) | LUTEIN DEFICIENT 5 (LUT5);"CYTOCHROME P450, FAMILY 97, SUBFAMILY A, POLYPEPTIDE 3" (CYP97A3) |

| | | | | | |
|---|---|---|---|---|---|
| AT4G15110 | AT4G15110.1 | member of CYP97B | protein_coding | "CYTOCHROME P450, FAMILY 97, SUBFAMILY B, POLYPEPTIDE 3" (CYP97B3) | "CYTOCHROME P450, FAMILY 97, SUBFAMILY B, POLYPEPTIDE 3" (CYP97B3) |
| AT5G43280 | AT5G43280.1 | Encodes the peroxisomal delta 3,5-delta2,4-dienoyl-CoA isomerase, a enzyme involved in degradation of unsaturated fatty acids. Gene expression is induced upon seed germination. | protein_coding | "DELTA(3,5),DELTA(2,4)-DIENOYL-COA ISOMERASE 1" (DCI1) | "DELTA(3,5),DELTA(2,4)-DIENOYL-COA ISOMERASE 1" (ATDCI1);DELTA(3,5),DELTA(2,4)-DIENOYL-COA ISOMERASE 1 (DCI1);"DELTA(3,5),DELTA(2,4)-DIENOYL-COA ISOMERASE 1" (DCI1) |
| AT1G10830 | AT1G10830.1 | Encodes a functional 15-cis-zeta-carotene isomerase (Z-ISO). | protein_coding | 15-CIS-ZETA-CAROTENE ISOMERASE (Z-ISO) | 15-CIS-ZETA-CAROTENE ISOMERASE (Z-ISO); (Z-ISO1.1) |
| AT5G47760 | AT5G47760.1 | serine/threonine protein kinase | protein_coding | 2-PHOSPHOGLYCOLATE PHOSPHATASE 2 (PGLP2) | (PGLP2);2-PHOSPHOGLYCOLATE PHOSPHATASE 2 (PGLP2); (ATPK5);2-PHOSPHOGLYCOLATE PHOSPHATASE 2 (ATPGLP2) |
| AT4G34030 | AT4G34030.1 | MCC-B is involved in leucine degradation in mitochondria. The active protein is a dimer of MCC-A and MCC-B. MCC-A is biotinylated whereas MCC-B is not. The mRNA is cell-to-cell mobile. | protein_coding | 3-METHYLCROTONYL-COA CARBOXYLASE (MCCB) | 3-METHYLCROTONYL-COA CARBOXYLASE (MCCB) |
| AT5G57850 | AT5G57850.1 | ADCL encodes a protein that acts as a 4-amino-4-deoxychorismate lyase. It catalyzes the production 4-aminobenzoate (pABA) production which is required for folate biosynthesis. The enzyme localizes to chloroplasts based on an import assay and GFP localization experiments. | protein_coding | 4-AMINO-4-DEOXYCHORISMATE LYASE (ADCL) | 4-AMINO-4-DEOXYCHORISMATE LYASE (ADCL) |
| AT2G05830 | AT2G05830.1 | Encodes a 5-methylthioribose-1-phosphate isomerase. | protein_coding | 5-METHYLTHIORIBOSE KINASE 1 (MTI1) | 5-METHYLTHIORIBOSE KINASE 1 (MTI1);5-METHYLTHIORIBOSE-1-PHOSPHATE-ISOMERASE1 (MTI1) |
| AT5G08530 | AT5G08530.1 | 51 kDa subunit of complex I;(source:Araport11) | protein_coding | 51 KDA SUBUNIT OF COMPLEX I (CI51) | 51 KDA SUBUNIT OF COMPLEX I (CI51); (NDUFV1) |
| AT1G04620 | AT1G04620.1 | Encodes a 7-hydroxymethyl chlorophyll a reductase, an enzyme of the chlorophyll cycle that converts 7-hydroxymethyl chlorophyll a to chlorophyll a. | protein_coding | 7-HYDROXYMETHYL CHLOROPHYLL A (HMCHL) REDUCTASE (HCAR) | 7-HYDROXYMETHYL CHLOROPHYLL A (HMCHL) REDUCTASE (HCAR) |
| AT5G46800 | AT5G46800.1 | Seedling lethal mutation; Mitochondrial Carnitine Acyl Carrier-Like Protein | protein_coding | A BOUT DE SOUFFLE (BOU) | A BOUT DE SOUFFLE (BOU) |
| AT5G67030 | AT5G67030.1 | Encodes a single copy zeaxanthin epoxidase gene that functions in first step of the biosynthesis of the abiotic stress hormone abscisic acid (ABA). Mutants in this gene are unable to express female sterility in response to beta-aminobutyric acid, as wild type plants do. | protein_coding | ABA DEFICIENT 1 (ABA1) | ABA DEFICIENT 1 (ABA1);LOW EXPRESSION OF OSMOTIC STRESS-RESPONSIVE GENES 6 (LOS6);NON-PHOTOCHEMICAL QUENCHING 2 (NPQ2);ARABIDOPSIS THALIANA ABA DEFICIENT 1 (ATABA1);ZEAXANTHIN EPOXIDASE (ZEP);IMPAIRED IN BABA-INDUCED STERILITY 3 (IBS3);ARABIDOPSIS THALIANA ZEAXANTHIN EPOXIDASE (ATZEP) |
| AT2G13540 | AT2G13540.1 | Encodes a nuclear cap-binding protein that forms a heterodimeric complex with CBP20 and is involved in ABA signaling and flowering. Mutants are early flowering and exhibit hypersensitive response to ABA in germination inhibition.Loss of ABH1 function results in abnormal processing of mRNAs for several important floral regulators (FLC, CO, FLM). Analysis of loss of function mutations suggests a role in pri-miRNA processing and mRNA splicing. Note that two different mutant alleles were given the same name abh1-7 (Kuhn et al 2007; Kim et al 2008). To avoid confusion, abh1-7 described in Kim et al (2008) has been renamed abh1-107 (other names: ensalada-1, ens-1). | protein_coding | ABA HYPERSENSITIVE 1 (ABH1) | ENSALADA (ENS);ABA HYPERSENSITIVE 1 (ABH1);CAP-BINDING PROTEIN 80 (CBP80); (ATCBP80) |
| AT5G13680 | AT5G13680.1 | A subunit of Elongator, a histone acetyl transferase complex, consisting of six subunits (ELP1?ELP6), that copurifies with the elongating RNAPII in yeast and humans. Three Arabidopsis thaliana genes, encoding homologs of the yeast Elongator subunits ELP1, ELP3 (histone acetyl transferase), | protein_coding | ABA-OVERLY SENSITIVE 1 (ABO1) | ELONGATA 2 (ELO2); (AtELP1);ABA-OVERLY SENSITIVE 1 (ABO1) |

| | | | | | |
|---|---|---|---|---|---|
| | | and ELP4 are responsible for the narrow leaf phenotype in elongata mutants and for reduced root growth that results from a decreased cell division rate. Mutants have no ncm5U (5-carbamoylmethyluridine). | | | |
| AT1G79600 | AT1G79600.1 | Encodes a chloroplast ABC1-like kinase that regulates vitamin E metabolism. | protein_coding | ABC1-LIKE KINASE 3 (ABC1K3) | ABC1-LIKE KINASE 3 (ABC1K3) |
| AT4G31390 | AT4G31390.1 | Protein kinase superfamily protein;(source:Araport11) | protein_coding | ABC1-LIKE KINASE RELATED TO CHLOROPHYLL DEGRADATION AND OXIDATIVE STRESS 1 (ACDO1) | ABC1-LIKE KINASE RELATED TO CHLOROPHYLL DEGRADATION AND OXIDATIVE STRESS 1 (ACDO1);PROTON GRADIENT REGULATION 6 (PGR6);ABC1-LIKE KINASE 1 (ABC1K1); (ATACDO1) |
| AT5G64940 | AT5G64940.1 | ABC1K8 is a member of an atypical protein kinase family that is induced by heavy metals. Loss of function mutations affect the metabolic profile of chloroplast lipids. It appears to function along with ABC1K7 in mediating lipid membrane changes in response to stress. The mRNA is cell-to-cell mobile. | protein_coding | ABC2 HOMOLOG 13 (ATH13) | (ABC1K8);ARABIDOPSIS THALIANA ABC2 HOMOLOG 13 (ATATH13);A. THALIANA OXIDATIVE STRESS-RELATED ABC1-LIKE PROTEIN 1 (ATOSA1);OXIDATIVE STRESS-RELATED ABC1-LIKE PROTEIN 1 (OSA1);ABC2 HOMOLOG 13 (ATH13) |
| AT3G44880 | AT3G44880.1 | Encodes a pheide a oxygenase (PAO). Accelerated cell death (acd1) mutants show rapid, spreading necrotic responses to both virulent and avirulent Pseudomonas syringae pv. maculicola or pv. tomato pathogens and to ethylene. | protein_coding | ACCELERATED CELL DEATH 1 (ACD1) | ACCELERATED CELL DEATH 1 (ACD1);PHEOPHORBIDE A OXYGENASE (PAO);LETHAL LEAF-SPOT 1 HOMOLOG (LLS1) |
| AT3G19720 | AT3G19720.1 | Encodes a novel chloroplast division protein. Mutants of exhibit defects in chloroplast constriction, have enlarged, dumbbell-shaped chloroplasts. The ARC5 gene product shares similarity with the dynamin family of GTPases, which mediate endocytosis, mitochondrial division, and other organellar fission and fusion events in eukaryotes. Phylogenetic analysis showed that ARC5 is related to a group of dynamin-like proteins unique to plants. A GFP-ARC5 fusion protein localizes to a ring at the chloroplast division site. Chloroplast import and protease protection assays indicate that the ARC5 ring is positioned on the outer surface of the chloroplast. Facilitates separation of the two daughter chloroplasts. | protein_coding | ACCUMULATION AND REPLICATION OF CHLOROPLAST 5 (ARC5) | DYNAMIN RELATED PROTEIN 5B (DRP5B);ACCUMULATION AND REPLICATION OF CHLOROPLAST 5 (ARC5) |
| AT5G42480 | AT5G42480.1 | Shows homology to the cyanobacterial cell division protein Ftn2, mutant only has two mesophyll cell chloroplasts. Protein was localized to a ring at the center of the chloroplasts. Probably involved in functions in the assembly and/or stabilization of the plastid-dividing FtsZ ring, inhibiting FtsZ filament formation in the chloroplast. | protein_coding | ACCUMULATION AND REPLICATION OF CHLOROPLASTS 6 (ARC6) | ACCUMULATION AND REPLICATION OF CHLOROPLASTS 6 (ARC6) |
| AT1G64810 | AT1G64810.2 | Encodes a chloroplast localized RNA binding protein that is involved in group II intron splicing. Splicing defects can account for the loss of photosynthetic complexes in apo1 mutants. | protein_coding | ACCUMULATION OF PHOTOSYSTEM ONE 1 (APO1) | ACCUMULATION OF PHOTOSYSTEM ONE 1 (APO1) |
| AT3G27000 | AT3G27000.1 | encodes a protein whose sequence is similar to actin-related proteins (ARPs) in other organisms. its transcript level is down regulated by light and is expressed in very low levels in all organs examined. | protein_coding | ACTIN RELATED PROTEIN 2 (ARP2) | WURM (WRM);ACTIN RELATED PROTEIN 2 (ARP2);ACTIN RELATED PROTEIN 2 (ATARP2) |
| AT3G60830 | AT3G60830.1 | Encodes an actin-related protein required for normal embryogenesis, plant architecture and floral organ abscission. | protein_coding | ACTIN-RELATED PROTEIN 7 (ARP7) | (ARP7);ACTIN-RELATED PROTEIN 7 (ARP7);ACTIN-RELATED PROTEIN 7 (ATARP7) |
| AT1G31730 | AT1G31730.1 | Encodes a component of the AP4 complex and is involved in vacuolar sorting of storage proteins. | protein_coding | ADAPTOR PROTEIN COMPLEX 4E (AP4E) | ADAPTOR PROTEIN COMPLEX 4E (AP4E) |
| AT1G08520 | AT1G08520.1 | Encodes the CHLD subunit of the Mg-chelatase enzyme involved in chlorophyll biosynthesis. Lines carrying recessive mutations of this locus are white and seedling lethal. | protein_coding | ALBINA 1 (ALB1) | ALBINA 1 (ALB1); (ALB-1V); (CHLD);PIGMENT DEFECTIVE EMBRYO 166 (PDE166); (V157) |

| | | | | | |
|---|---|---|---|---|---|
| AT3G62910 | AT3G62910.1 | Encodes a plastid-localized ribosome release factor 1 that is essential in chloroplast development. Pale green, albino mutant seedlings arrest early in seedling development. | protein_coding | ALBINO AND PALE GREEN (APG3) | ALBINO AND PALE GREEN (APG3);CHLOROPLAST RIBOSOME RELEASE FACTOR 1 (CPRF1);ARABIDOPSIS THALIANA CHLOROPLAST RF1 (ATCPRF1) |
| AT5G62530 | AT5G62530.1 | Encodes mitochondrial Delta-pyrroline-5- carboxylate dehydrogenase. Involved in the catabolism of proline to glutamate. Involved in protection from proline toxicity. Induced at pathogen infection sites. P5CDH and SRO5 (an overlapping gene in the sense orientation) generate 24-nt and 21-nt siRNAs, which together are components of a regulatory loop controlling reactive oxygen species (ROS) production and stress response. | protein_coding | ALDEHYDE DEHYDROGENASE 12A1 (ALDH12A1) | ALDEHYDE DEHYDROGENASE 12A1 (ALDH12A1);DELTA1-PYRROLINE-5-CARBOXYLATE DEHYDROGENASE (P5CDH);ARABIDOPSIS THALIANA DELTA1-PYRROLINE-5-CARBOXYLATE DEHYDROGENASE (ATP5CDH) |
| AT3G66658 | AT3G66658.2 | Encodes a putative aldehyde dehydrogenase. The gene is not responsive to osmotic stress and is expressed constitutively at a low level in plantlets and root cultures. | protein_coding | ALDEHYDE DEHYDROGENASE 22A1 (ALDH22a1) | ALDEHYDE DEHYDROGENASE 22A1 (ALDH22a1) |
| AT4G04955 | AT4G04955.1 | Encodes an allantoinase which is involved in allantoin degradation and assimilation. Gene expression was induced when allantoin was added to the medium. The insertion mutant, ataln m2-1, did not grow well on the MS medium where allantoin, instead of ammonium nitrate, was supplied. | protein_coding | ALLANTOINASE (ALN) | ALLANTOINASE (ATALN);ALLANTOINASE (ALN) |
| AT1G76130 | AT1G76130.1 | alpha-amylase, putative / 1,4-alpha-D-glucan glucanohydrolase, putative, strong similarity to alpha-amylase GI:7532799 from (Malus x domestica);contains Pfam profile PF00128: Alpha amylase, catalytic domain. Predicted to be secreted based on SignalP analysis. | protein_coding | ALPHA-AMYLASE-LIKE 2 (AMY2) | ALPHA-AMYLASE-LIKE 2 (AMY2);ARABIDOPSIS THALIANA ALPHA-AMYLASE-LIKE 2 (ATAMY2) |
| AT1G69830 | AT1G69830.1 | Encodes a plastid-localized α-amylase. Expression is reduced in the SEX4 mutant. Loss of function mutations show normal diurnal pattern of starch accumulation/degradation. Expression follows circadian rhythms. | protein_coding | ALPHA-AMYLASE-LIKE 3 (AMY3) | ALPHA-AMYLASE-LIKE 3 (AMY3);ALPHA-AMYLASE-LIKE 3 (ATAMY3) |
| AT5G08370 | AT5G08370.1 | Member of Glycoside Hydrolase Family 27 (GH27)that functions as an α-galactosidase. | protein_coding | ALPHA-GALACTOSIDASE 2 (AGAL2) | ALPHA-GALACTOSIDASE 2 (AGAL2);ALPHA-GALACTOSIDASE 2 (AtAGAL2) |
| AT4G10030 | AT4G10030.1 | Alpha/beta hydrolase domain containing protein involved in lipid biosynthesis. | protein_coding | ALPHA/BETA HYDROLASE DOMAIN 11 (ABHD11) | ALPHA/BETA HYDROLASE DOMAIN 11 (ABHD11) |
| AT2G04660 | AT2G04660.1 | a highly conserved ubiquitin-protein ligase involved in cell cycle regulation | protein_coding | ANAPHASE-PROMOTING COMPLEX/CYCLOSOME 2 (APC2) | ANAPHASE-PROMOTING COMPLEX/CYCLOSOME 2 (APC2) |
| AT2G21120 | AT2G21120.1 | Encodes a putative magnesium transporter that was identified through a forward genetic screen, directly isolating antiviral RNAi-defective (avi) mutant using a Cucumber Mosaic Virus (CMV) mutant. Compared to Wildtype Col-0, avi2 mutant showed severe disease symptom after viral infection and viral accumulation was significantly increased while viral siRNAs and virus-activated endogenous siRNAs (vasiRNAs) were reduced in avi2 mutant. Detailed genetic study indicated that AVI2 modulated RNAi-mediated antiviral immunity by regulating the biogenesis of secondary viral siRNAs and vasiRNAs in Arabidopsis. | protein_coding | ANTIVIRAL RNAI-DEFECTIVE 2 (AVI2) | ENHANCER OF RDR6 3 (ENOR3);ANTIVIRAL RNAI-DEFECTIVE 2 (AVI2) |
| AT2G31440 | AT2G31440.1 | Encodes a gamma-secretase subunit. Associates with other subunits in intracellular membrane compartments. | protein_coding | APH-1 (APH-1) | APH-1 (APH-1) |
| AT3G23620 | AT3G23620.1 | BRIX domain containing protein, similar to RNA biogenesis factors in yeast. Binds rRNA and likely also functions in RNA biogenesis in Arabidopsis. Essential gene, mutants are embryo lethal and does not transmit well through the gametophyte. | protein_coding | ARABIDOPSIS HOMOLOG OF YEAST RPF2 (ARPF2) | ARABIDOPSIS HOMOLOG OF YEAST RPF2 (ARPF2) |
| AT1G13330 | AT1G13330.1 | Encodes the Arabidopsis Hop2 homologue. In other species, Hop2 is proposed to be involved in inter-homolog bias in double strand break repair. | protein_coding | ARABIDOPSIS HOP2 HOMOLOG (AHP2) | HOMOLOGOUS-PAIRING PROTEIN 2 (HOP2);ARABIDOPSIS HOP2 HOMOLOG (AHP2) |

| | | | | | |
|---|---|---|---|---|---|
| AT5G15550 | AT5G15550.1 | Transducin/WD40 repeat-like superfamily protein;(source:Araport11) | protein_coding | ARABIDOPSIS THALIANA PESCADILLO ORTHOLOG (ATPEP2) | (ATPEIP2);ARABIDOPSIS THALIANA PESCADILLO ORTHOLOG (ATPEP2) |
| AT5G08100 | AT5G08100.1 | Encodes an asparaginase that catalyzes the degradation of L-asparagine to L-aspartic acid and ammonia. | protein_coding | ASPARAGINASE A1 (ASPGA1) | ASPARAGINASE A1 (ASPGA1) |
| AT2G47760 | AT2G47760.1 | Encodes an α-1,3-mannosyltransferase. Plants with mutations in the ALG3 protein have abnormal gylcoslation profiles. They also exhibit abnormal responses to MAMPs possibly because the glycan properties of FL22 are affected. | protein_coding | ASPARAGINE-LINKED GLYCOSYLATION 3 (ALG3) | ARABIDOPSIS THALIANA ASPARAGINE-LINKED GLYCOSYLATION 3 (AtALG3);ASPARAGINE-LINKED GLYCOSYLATION 3 (ALG3) |
| AT4G31990 | AT4G31990.3 | Encodes a plastid-localized aspartate aminotransferase. Does not display any PAT (glutamate/aspartate-prephenate aminotransferase) activity even in the presence of a high concentration of prephenate. | protein_coding | ASPARTATE AMINOTRANSFERASE 5 (ASP5) | ASPARTATE AMINOTRANSFERASE 5 (ASP5); (ATAAT1);ASPARTATE AMINOTRANSFERASE DEFICIENT 3 (AAT3) |
| AT1G54350 | AT1G54350.1 | ABC transporter family protein;(source:Araport11) | protein_coding | ATP-BINDING CASSETTE D2 (ABCD2) | ATP-BINDING CASSETTE D2 (ABCD2) |
| AT1G64550 | AT1G64550.1 | Encodes a member of GCN subfamily. Predicted to be involved in stress-associated protein translation control. The mutant is affected in MAMP ((microbe-associated molecular patterns)-induced stomatal closure, but not other MAMP-induced responses in the leaves. | protein_coding | ATP-BINDING CASSETTE F3 (ABCF3) | GENERAL CONTROL NON-REPRESSIBLE 3 (ATGCN3);GENERAL CONTROL NON-REPRESSIBLE 3 (GCN3);ATP-BINDING CASSETTE F3 (ABCF3); (ATGCN20);SUSCEPTIBLE TO CORONATINE-DEFICIENT PST DC3000 5 (SCORD5);GENERAL CONTROL NON-REPRESSIBLE 20 (GCN20); (ATABCF3) |
| AT2G28070 | AT2G28070.1 | ABC-2 type transporter family protein;(source:Araport11) | protein_coding | ATP-BINDING CASSETTE G3 (ABCG3) | ATP-BINDING CASSETTE G3 (ABCG3) |
| AT2G01320 | AT2G01320.3 | ABC-2 type transporter family protein;(source:Araport11) | protein_coding | ATP-BINDING CASSETTE G7 (ABCG7) | ATP-BINDING CASSETTE G7 (ABCG7) |
| AT1G63270 | AT1G63270.1 | Encodes a member of a heterogenous group of non-intrinsic ATP-binding cassette (ABC) proteins. Members of this group bear no close resemblance to each other nor to representatives of specific ABC protein subfamilies from other organisms. This grouping is arbitrary and will likely change upon acquisition of further data. | protein_coding | ATP-BINDING CASSETTE I1 (ABCI1) | NON-INTRINSIC ABC PROTEIN 10 (ATNAP10);CYTOCHROME C MATURATION A (ATCCMA);ATP-BINDING CASSETTE I1 (ABCI1);NON-INTRINSIC ABC PROTEIN 10 (NAP10) |
| AT1G19800 | AT1G19800.2 | Encodes a permease-like protein involved in lipid transfer from the ER to the chloroplast, more specifically, transfer of phosphatidate across the chloroplast inner membrane. Mutant leaves accumulate trigalactosyldiacylglycerol, triacylglycerol and phosphatidate. Chloroplast lipids are altered in their fatty acid composition and as a consequence the development of chloroplasts in the mutants are impacted. The mutant seeds has a higher abortion rate. Mutations in this gene suppress the low temperature-induced phenotype of Arabidopsis tocopherol-deficient mutant vte2. | protein_coding | ATP-BINDING CASSETTE I14 (ABCI14) | TRIGALACTOSYLDIACYLGLYCEROL 1 (TGD1);ATP-BINDING CASSETTE I14 (ABCI14) |
| AT5G02270 | AT5G02270.2 | member of NAP subfamily | protein_coding | ATP-BINDING CASSETTE I20 (ABCI20) | ATP-BINDING CASSETTE I20 (ABCI20);NON-INTRINSIC ABC PROTEIN 9 (NAP9) |
| AT5G48520 | AT5G48520.1 | Encodes AUGMIN subunit3 (AUG3), a homolog of animal dim gamma-tubulin 3/human augmin-like complex, subunit 3. Plays a critical role in microtubule organization during plant cell division. | protein_coding | AUGMIN 3 (AUG3) | (AtAUG3);AUGMIN 3 (AUG3) |
| AT5G50230 | AT5G50230.1 | autophagy-related (ATG) gene | protein_coding | AUTOPHAGY 16 (ATG16) | AUTOPHAGY 16 (ATG16) |
| AT5G45900 | AT5G45900.1 | Component of autophagy conjugation pathway. Required for proper senescence. Contributes to plant basal immunity towards fungal infection. | protein_coding | AUTOPHAGY 7 (APG7) | AUTOPHAGY 7 (APG7); (ATAPG7); (ATATG7);AUTOPHAGY-RELATED 7 (ATG7);PEROXISOME UNUSUAL POSITIONING 4 (PEUP4) |
| AT4G08540 | AT4G08540.1 | DNA-directed RNA polymerase II protein;(source:Araport11) | protein_coding | BECLIN 1-ASSOCIATED AUTOPHAGY-RELATED KEY REGULATOR 14B (ATG14B) | BECLIN 1-ASSOCIATED AUTOPHAGY-RELATED KEY REGULATOR 14B (ATG14B) |
| AT4G02990 | AT4G02990.1 | Encodes BELAYA SMERT (BSM), a plastid-localized protein homologous to mitochondrial transcription termination factors (mTERF) found in animal. Mutant bsm | protein_coding | BELAYA SMERT (BSM) | RUGOSA 2 (RUG2);BELAYA SMERT (BSM) |

| | | | | | |
|---|---|---|---|---|---|
| | | cells are albino, are compromised in growth, and suffer defects in global plastidic gene expression. The mRNA is cell-to-cell mobile. | | | |
| AT3G61320 | AT3G61320.1 | Encodes a bestrophin-like protein (Best1). Located in the stroma thylakoid membrane. Functions as a chloride ion channel. Proposed to modulate proton motive force partitioning by mediating chloride ion influx in the thylakoid lumen. Major isoform (based on transcript analysis), redundant function with AtBest2. | protein_coding | BESTROPHIN-LIKE PROTEIN (BEST) | (ATVCCN1);BESTROPHIN-LIKE PROTEIN (BEST) |
| AT3G55260 | AT3G55260.1 | Encodes a protein with β-hexosaminidase activity (the enzyme is active with p-nitrophenyl-β-N-acetylglucosaminide as substrate but displayed only a minor activity toward p-nitrophenyl-β-N-acetylgalactosaminide). The enzyme displays no distinct preference for a specific terminal GlcNAc residue and indeed cleaved the asialoagalactodabsylglycopeptide GnGn to a mixture of products. | protein_coding | BETA-HEXOSAMINIDASE 1 (HEXO1) | BETA-HEXOSAMINIDASE 1 (HEXO1); (ATHEX2) |
| AT5G64370 | AT5G64370.1 | PYD3 encodes a beta-ureidopropionase which, when expressed in E. coli, has been shown to convert beta-ureidopropionate into beta-alanine. It localizes to the cytosol and plays an important role in uracil degradation. | protein_coding | BETA-UREIDOPROPIONASE (BETA-UP) | PYRIMIDINE 3 (PYD3);BETA-UREIDOPROPIONASE (BETA-UP) |
| AT1G11190 | AT1G11190.1 | Encodes a bifunctional nuclease that acts on both RNA and DNA involved in nucleic acid degradation to facilitate nucleotide and phosphate recovery during senescence. It has mismatch-specific endonuclease activity with wide recognition of single base mismatches as well as the ability to cleave indel types of mismatches (heteroduplexes with loops). | protein_coding | BIFUNCTIONAL NUCLEASE I (BFN1) | ENDONUCLEASE 1 (ENDO1);BIFUNCTIONAL NUCLEASE I (BFN1) |
| AT2G26900 | AT2G26900.1 | Sodium Bile acid symporter family;(source:Araport11) | protein_coding | BILE ACID:SODIUM SYMPORTER FAMILY PROTEIN 2 (BASS2) | BILE ACID:SODIUM SYMPORTER FAMILY PROTEIN 2 (BASS2) |
| AT4G30825 | AT4G30825.1 | P-class pentatricopeptide repeat (PPR) protein essential for accumulation of the dicistronic atpH/F transcript in chloroplasts. Acts as barrier to prevent the atpH/F transcript degradation by exoribonucleases by binding to the consensus sequence of the atpF-atpA intergenic region. | protein_coding | BIOGENESIS FACTOR REQUIRED FOR ATP SYNTHASE 2 (BFA2) | BIOGENESIS FACTOR REQUIRED FOR ATP SYNTHASE 2 (BFA2) |
| AT5G57590 | AT5G57590.1 | Encodes a bifunctional enzyme with both dethiobiotin synthetase and diaminopelargonic acid aminotransferase activities that is involved in biotin synthesis. | protein_coding | BIOTIN AUXOTROPH 1 (BIO1) | BIOTIN AUXOTROPH 1 (BIO1) |
| AT2G33560 | AT2G33560.2 | Encodes BUBR1. May have the spindle assembly checkpoint protein functions conserved from yeast to humans. | protein_coding | BUB1-RELATED (BUB1: BUDDING UNINHIBITED BY BENZYMIDAZOL 1) (BUBR1) | BUB1-RELATED (BUB1: BUDDING UNINHIBITED BY BENZYMIDAZOL 1) (BUBR1) |
| AT3G17470 | AT3G17470.1 | Ca2+-activated RelA/spot-like protein;(source:Araport11) | protein_coding | CA2+-ACTIVATED RELA/SPOT HOMOLOG (CRSH) | CA2+-ACTIVATED RELA/SPOT HOMOLOG (CRSH); (ATCRSH) |
| AT1G03910 | AT1G03910.2 | Encodes the Arabidopsis homolog of a conserved eukaryotic protein without known functional domains. The protein that localizes to nuclear speckles and colocalizes with known splicing proteins. | protein_coding | CACTIN (CTN) | CACTIN (CTN) |
| AT5G15730 | AT5G15730.2 | Protein kinase superfamily protein;(source:Araport11) | protein_coding | CALCIUM/CALMODULIN-REGULATED RECEPTOR-LIKE KINASE 2 (CRLK2) | (AtCRLK2);CALCIUM/CALMODULIN-REGULATED RECEPTOR-LIKE KINASE 2 (CRLK2) |
| AT1G71790 | AT1G71790.1 | Encodes a heterodimeric actin binding protein composed of an alpha and a beta sumunit. Stabilizes actin filament cytoskeleton by capping. | protein_coding | CAPPING PROTEIN B (CPB) | (ATCPB);CAPPING PROTEIN B (CPB) |
| AT1G08960 | AT1G08960.1 | Encodes a member of the Potassium-dependent sodium-calcium exchanger like-family that localizes to the plasma membrane and nuclear periphery, and has a role in mediating high-afﬁnity K+ uptake and Na+ transport in yeast. | protein_coding | CATION EXCHANGER 11 (CAX11) | CATION EXCHANGER 11 (CAX11);ARABIDOPSIS THALIANA CATION CALCIUM EXCHANGER 5 (AtCXX5);CATION CALCIUM EXCHANGER 5 (CCX5);CATION EXCHANGER 11 (ATCAX11) |

| | | | | | |
|---|---|---|---|---|---|
| AT1G05940 | AT1G05940.1 | Encodes a member of the cationic amino acid transporter (CAT) subfamily of amino acid polyamine choline transporters. | protein_coding | CATIONIC AMINO ACID TRANSPORTER 9 (CAT9) | CATIONIC AMINO ACID TRANSPORTER 9 (CAT9) |
| AT1G65320 | AT1G65320.1 | Cystathionine beta-synthase (CBS) family protein;(source:Araport11) | protein_coding | CBS DOMAIN CONTAINING PROTEIN 6 (CBSX6) | CBS DOMAIN CONTAINING PROTEIN 6 (CBSX6) |
| AT4G28980 | AT4G28980.2 | Encodes a CDK-activating kinase that regulates root initial cell differentiation. Phosphorylates CDKD2 and CDKD3, but not CDKD1. Controls CDK activities and basal transcription. | protein_coding | CDK-ACTIVATING KINASE 1AT (CAK1AT) | CDK-ACTIVATING KINASE 1AT (CAK1AT);CYCLIN-DEPENDENT KINASE F;1 (CDKF;1) |
| AT4G13590 | AT4G13590.1 | Chloroplast manganese transporter required for chloroplast manganese homeostasis and photosynthetic function. | protein_coding | CHLOROPLAST MANGANESE TRANSPORTER1 (CMT1) | CHLOROPLAST MANGANESE TRANSPORTER1 (CMT1) |
| AT1G76080 | AT1G76080.1 | Encodes a thioredoxin localized in chloroplast stroma. Known as CDSP32 (CHLOROPLASTIC DROUGHT-INDUCED STRESS PROTEIN OF 32 KD). | protein_coding | CHLOROPLASTIC DROUGHT-INDUCED STRESS PROTEIN OF 32 KD (CDSP32) | CHLOROPLASTIC DROUGHT-INDUCED STRESS PROTEIN OF 32 KD (CDSP32);ARABIDOPSIS THALIANA CHLOROPLASTIC DROUGHT-INDUCED STRESS PROTEIN OF 32 KD (ATCDSP32) |
| AT1G08490 | AT1G08490.1 | Chloroplastic NifS-like protein that can catalyze the conversion of cysteine into alanine and elemental sulfur (S(0)) and of selenocysteine into alanine and elemental Se (Se(0)). Overexpression enhances selenium tolerance and accumulation. | protein_coding | CHLOROPLASTIC NIFS-LIKE CYSTEINE DESULFURASE (CPNIFS) | CHLOROPLASTIC NIFS-LIKE CYSTEINE DESULFURASE (CPNIFS); (ATSUFS); (ATNFS2); (ATCPNIFS); (SUFS) |
| AT5G55740 | AT5G55740.1 | Encodes a member of the E+ subgroup of the PPR protein family, containing the E and E+ motifs following a tandem array of PPR motifs. It also contains an unknown motif consisting of 15 aa, which is highly conserved in some PPR proteins, including CRR4. CRR21 is involved in RNA editing of the site 2 of ndhD (ndhD-2),which encodes a subunit of the NDH complex. The RNA editing changes aa 128 from Ser to Leu. Mutants have impaired NDH complex activity. | protein_coding | CHLORORESPIRATORY REDUCTION 21 (CRR21) | CHLORORESPIRATORY REDUCTION 21 (CRR21) |
| AT1G11290 | AT1G11290.1 | Pentatricopeptide Repeat Protein containing the DYW motif. Required for editing of multiple plastid transcripts. Endonuclease activity. | protein_coding | CHLORORESPIRATORY REDUCTION22 (CRR22) | CHLORORESPIRATORY REDUCTION22 (CRR22) |
| AT1G65380 | AT1G65380.1 | Receptor-like protein containing leucine-rich repeats. Regulates both meristem and organ development in Arabidopsis. | protein_coding | CLAVATA 2 (CLV2) | RECEPTOR LIKE PROTEIN 10 (AtRLP10);CLAVATA 2 (CLV2) |
| AT1G12410 | AT1G12410.1 | Encodes a ClpP-related sequence. Though similar to ClpP proteins, this does not contains the highly conserved catalytic triad of Ser-type proteases (Ser-His-Asp). The name reflects nomenclature described in Adam et. al (2001). | protein_coding | CLP PROTEASE PROTEOLYTIC SUBUNIT 2 (CLP2) | (CLPR2);EMBRYO DEFECTIVE 3146 (EMB3146);CLP PROTEASE PROTEOLYTIC SUBUNIT 2 (CLP2);NUCLEAR-ENCODED CLP PROTEASE P2 (NCLPP2) |
| AT1G66670 | AT1G66670.1 | One of several nuclear-encoded ClpPs (caseinolytic protease). Contains a highly conserved catalytic triad of Ser-type proteases (Ser-His-Asp). The name reflects nomenclature described in Adam et. al (2001). | protein_coding | CLP PROTEASE PROTEOLYTIC SUBUNIT 3 (CLPP3) | CLP PROTEASE PROTEOLYTIC SUBUNIT 3 (CLPP3); (NCLPP3) |
| AT4G26180 | AT4G26180.1 | Encodes a mitochondrial CoA transporter. | protein_coding | COA CARRIER 2 (COAC2) | COA CARRIER 2 (COAC2) |
| AT4G38240 | AT4G38240.1 | Encodes N-acetyl glucosaminyl transferase I, the first enzyme in the pathway of complex glycan biosynthesis. | protein_coding | COMPLEX GLYCAN LESS 1 (CGL1) | COMPLEX GLYCAN LESS 1 (CGL1);N-ACETYLGLUCOSAMINYLTRANSFERASE I (GNTI);COMPLEX GLYCAN LESS (CGL) |
| AT2G34560 | AT2G34560.2 | P-loop containing nucleoside triphosphate hydrolases superfamily protein;(source:Araport11) | protein_coding | CONSERVED IN CILIATED SPECIES AND IN THE LAND PLANTS 1 (CCP1) | (ATCCP1);CONSERVED IN CILIATED SPECIES AND IN THE LAND PLANTS 1 (CCP1) |
| AT1G73430 | AT1G73430.1 | COG3 is a component of a putative conserved oligomeric Golgi (COG) complex that is thought to be involved in tethering of retrograde intra Golgi vesicles. In mutant pollen,golgi appear abnormal. It is required for proper deposition of cell wall materials in pollen tube growth. When homozygotes can be produced (by complementing the | protein_coding | CONSERVED OLIGOMERIC GOLGI COMPLEX 3 (COG3) | CONSERVED OLIGOMERIC GOLGI COMPLEX 3 (COG3) |

| | | | | | |
|---|---|---|---|---|---|
| | | defect in pollen), the plants are embryo lethal suggesting an essential function. COG3 interacts with several other putative COG components. | | | |
| AT1G31780 | AT1G31780.1 | oligomeric golgi complex subunit;(source:Araport11) | protein_coding | CONSERVED OLIGOMERIC GOLGI COMPLEX 7 (COG7) | CONSERVED OLIGOMERIC GOLGI COMPLEX 7 (COG7) |
| AT5G14250 | AT5G14250.1 | Encodes subunit 3 of the COP9 signalosome. | protein_coding | CONSTITUTIVE PHOTOMORPHOGENIC 13 (COP13) | FUSCA 11 (FUS11);COP9 SIGNALOSOME SUBUNIT 3 (CSN3);CONSTITUTIVE PHOTOMORPHOGENIC 13 (COP13) |
| AT3G56940 | AT3G56940.1 | Encodes a putative ZIP protein with varying mRNA accumulation in leaves, stems and roots. Has a consensus carboxylate-bridged di-iron binding site. The mRNA is cell-to-cell mobile. | protein_coding | COPPER RESPONSE DEFECT 1 (CRD1) | COPPER RESPONSE DEFECT 1 (CRD1); (ACSF); (CHL27) |
| AT5G24850 | AT5G24850.1 | Binds flavin adenine dinucleotide and DNA. It does not have photolyase activity, and it is likely to act as photoreceptor. Closely related to Synechocystis cryptochrome. | protein_coding | CRYPTOCHROME 3 (CRY3) | CRYPTOCHROME 3 (CRY3) |
| AT3G01480 | AT3G01480.1 | Encodes a chloroplast cyclophilin functioning in the assembly and maintenance of photosystem II (PSII) supercomplexes. The mRNA is cell-to-cell mobile. | protein_coding | CYCLOPHILIN 38 (CYP38) | ARABIDOPSIS CYCLOPHILIN 38 (ATCYP38);CYCLOPHILIN 38 (CYP38) |
| AT4G33060 | AT4G33060.1 | Cyclophilin-like peptidyl-prolyl cis-trans isomerase family protein;(source:Araport11) | protein_coding | CYCLOPHILIN 57 (CYP57) | (ATCYP57);CYCLOPHILIN 57 (CYP57) |
| AT2G31170 | AT2G31170.1 | Encodes the cysteinyl t-RNA synthetase SYCO ARATH (SYCO), which is expressed and required in the central cell but not in the antipodals. SYCO, localized to the mitochondria, is necessary for mitochondrial cristae integrity. Mutation of this gene affects the lifespan of adjacent accessory cells. | protein_coding | CYSTEINYL T-RNA SYNTHETASE (SYCO ARATH) | FIONA (FIONA);CYSTEINYL T-RNA SYNTHETASE (SYCO ARATH) |
| AT5G38630 | AT5G38630.1 | Encodes for cytochrome b561. | protein_coding | CYTOCHROME B561-1 (CYB-1) | CYTOCHROME B561-1 (CYB-1);CYTOCHROME B561-1 (ACYB-1) |
| AT2G35030 | AT2G35030.1 | Pentatricopeptide repeat (PPR) superfamily protein;(source:Araport11) | protein_coding | CYTOCHROME C OXIDASE DEFICIENT 1 (COD1) | CYTOCHROME C OXIDASE DEFICIENT 1 (COD1) |
| AT4G35910 | AT4G35910.1 | Encodes a cytoplasmic thiouridylase that is essential for tRNA thiolation. Its activity appears to be important in root development. | protein_coding | CYTOPLASMIC THIOURIDYLASE 2 (CTU2) | CYTOPLASMIC THIOURIDYLASE 2 (CTU2) |
| AT2G29560 | AT2G29560.1 | Encodes a putative phosphoenolpyruvate enolase that is localized both to the nucleus and the cytoplasm. The mRNA is cell-to-cell mobile. | protein_coding | CYTOSOLIC ENOLASE (ENOC) | ENOLASE 3 (ENO3);CYTOSOLIC ENOLASE (ENOC) |
| AT4G36400 | AT4G36400.1 | Encodes a (D)-2-hydroxyglutarate dehydrogenase. | protein_coding | D-2-HYDROXYGLUTARATE DEHYDROGENASE (D2HGDH) | D-2-HYDROXYGLUTARATE DEHYDROGENASE (D2HGDH) |
| AT5G06580 | AT5G06580.1 | Encodes a protein with glycolate dehydrogenase activity, which was shown to complement various subunits of the E. coli glycolate oxidase complex. It has not been ruled out that the enzyme might be involved in other catalytic activities in vivo. | protein_coding | D-LACTATE DEHYDROGENASE (D-LDH) | D-LACTATE DEHYDROGENASE (D-LDH) |
| AT5G58760 | AT5G58760.1 | Encodes a DDB1a interacting protein DDB2 required for UV-B tolerance and genomic integrity. | protein_coding | DAMAGED DNA BINDING 2 (DDB2) | DAMAGED DNA BINDING 2 (DDB2) |
| AT1G14620 | AT1G14620.1 | decoy;(source:Araport11) | protein_coding | DECOY (DECOY) | DECOY (DECOY) |
| AT2G47940 | AT2G47940.1 | Encodes DegP2 protease (DEGP2); nuclear gene for chloroplast product. | protein_coding | DEGRADATION OF PERIPLASMIC PROTEINS 2 (DEG2) | EMBRYO DEFECTIVE 3117 (EMB3117);DEGP PROTEASE 2 (DEGP2);DEGRADATION OF PERIPLASMIC PROTEINS 2 (DEG2) |
| AT5G39830 | AT5G39830.1 | Encodes DEG8. Forms a hexamer with DEG5 in the thylakoid lumen. Involved in the cleavage of photodamaged D2 protein of photosystem II (PSII). Recombinant DEG8 is proteolytically active toward both a model substrate (beta-casein) and photodamaged D1 protein of photosystem II. | protein_coding | DEGRADATION OF PERIPLASMIC PROTEINS 8 (DEG8) | DEG PROTEASE 8 (DEGP8);DEGRADATION OF PERIPLASMIC PROTEINS 8 (DEG8) |

| | | | | | |
|---|---|---|---|---|---|
| AT5G67570 | AT5G67570.1 | Encodes a pentratricopeptide repeat containing protein that is targeted to the chloroplast. Mutants have pale young leave and reduced accumulation of plastid encoded transcripts suggesting a role for DG1 in regulation of plastid gene expression. | protein_coding | DELAYED GREENING 1 (DG1) | EMBRYO DEFECTIVE 1408 (EMB1408);EMBRYO DEFECTIVE 246 (EMB246);DELAYED GREENING 1 (DG1) |
| AT5G05920 | AT5G05920.1 | Encodes a deoxyhypusine synthase. | protein_coding | DEOXYHYPUSINE SYNTHASE (DHS) | EMBRYO SAC DEVELOPMENT ARREST 22 (EDA22);DEOXYHYPUSINE SYNTHASE (DHS) |
| AT1G63900 | AT1G63900.2 | Encodes a RING-type ubiquitin E3 ligase of the chloroplast outer membrane that associates with TOC complexes and mediates ubiquitination of TOC components, promoting their degradation. It not only regulates chloroplast protein import but also targets components of the peroxisome protein import apparatus, PEX13 in particular. Several studies have been done to examine the peroxisomal localization of this protein, with varying interpretations. | protein_coding | DIAP1-LIKE PROTEIN 1 (DAL1) | SUPPRESSOR OF PPI1 LOCUS 1 (SP1);DIAP1-LIKE PROTEIN 1 (DAL1) |
| AT1G27980 | AT1G27980.1 | dihydrosphingosine phosphate lyase;(source:Araport11) | protein_coding | DIHYDROSPHINGOSINE PHOSPHATE LYASE (DPL1) | (ATDPL1);DIHYDROSPHINGOSINE PHOSPHATE LYASE (DPL1) |
| AT3G23940 | AT3G23940.1 | Encodes a member of the dihydroxyacid dehydrates family of proteins that encode enzymes involved in branched chain amino acid biosynthesis. Loss of function mutations have significantly reduced transmission and fertility due to defects in male and female gametophyte development and embryo lethality. Mutants have increased sensitivity to abiotic stressors which may be partially compensated by addition of amino acids to the growth medium. | protein_coding | DIHYDROXYACID DEHYDRATASE (DHAD) | DIHYDROXYACID DEHYDRATASE (DHAD) |
| AT5G64860 | AT5G64860.1 | Encodes a maltotriose-metabolizing enzyme with chloroplastic α-1,4-glucanotransferase activity. Mutant has altered starch degradation. | protein_coding | DISPROPORTIONATING ENZYME (DPE1) | (ATDPE1);DISPROPORTIONATING ENZYME (DPE1) |
| AT3G22880 | AT3G22880.1 | Expression of the AtDMC1 is restricted to pollen mother cells in anthers and to megaspore mother cells in ovules. Similar to meiosis-specific yeast DMC gene. | protein_coding | DISRUPTION OF MEIOTIC CONTROL 1 (DMC1) | DISRUPTION OF MEIOTIC CONTROL 1 (DMC1);ARABIDOPSIS THALIANA DISRUPTION OF MEIOTIC CONTROL 1 (ATDMC1);ARABIDOPSIS HOMOLOG OF LILY MESSAGES INDUCED AT MEIOSIS 15 (ARLIM15) |
| AT1G53280 | AT1G53280.1 | Encodes a homolog of animal DJ-1 superfamily protein. In the A. thaliana genome, three genes encoding close homologs of human DJ-1 were identified AT3G14990 (DJ1A), AT1G53280 (DJ1B) and AT4G34020 (DJ1C). Among the three homologs, DJ1C is essential for chloroplast development and viability. It exhibits glyoxalase activity towards glyoxal and methylglyoxal. | protein_coding | DJ-1 HOMOLOG B (DJ1B) | DJ-1 HOMOLOG B (DJ1B);DJ-1 HOMOLOG B (AtDJ1B); (DJ-1B) |
| AT1G80030 | AT1G80030.1 | Molecular chaperone Hsp40/DnaJ family protein;(source:Araport11) | protein_coding | DNA J PROTEIN A7 (DJA7) | DNA J PROTEIN A7 (DJA7) |
| AT5G25480 | AT5G25480.1 | Encodes a DNA methyltransferase homolog. Human Dnmt2 methylates tRNA-Asp and can methylate Arabidopsis tRNA-Asp in vitro. | protein_coding | DNA METHYLTRANSFERASE-2 (DNMT2) | DNA METHYLTRANSFERASE 2 (AtDNMT2); (TRDMT1);DNA METHYLTRANSFERASE-2 (DNMT2) |
| AT1G10520 | AT1G10520.1 | Encodes a homolog of the mammalian DNA polymerase lambda that is involved in the repair of UV-B induced DNA damage. | protein_coding | DNA POLYMERASE {LAMBDA} (Pol{lambda}) | (AtPol{lambda});DNA POLYMERASE {LAMBDA} (Pol{lambda}) |
| AT1G20575 | AT1G20575.1 | Encodes the catalytic core of the dolichol phosphate mannase synthase (DPMS) complex. It is not active on its own but requires the presence of DPMS2 and DPMS3 for full activity. It is localized in the ER and mediates isoprenyl-linked glycan biogenesis, inﬂuences development, stress response, and ammonium hypersensitivity. | protein_coding | DOLICHOL PHOSPHATE MANNOSE SYNTHASE 1 (DPMS1) | DOLICHOL PHOSPHATE MANNOSE SYNTHASE 1 (DPMS1) |

| | | | | | |
|---|---|---|---|---|---|
| AT5G19540 | AT5G19540.1 | DY1 is a novel nuclear encoded protein that is imported into the chloroplast stroma. Mutants have reduced pigmentation and somewhat abnormal thylakoid membranes. | protein_coding | DWARF AND YELLOW 1 (DY1) | DWARF AND YELLOW 1 (DY1) |
| AT2G40550 | AT2G40550.1 | Encodes a nuclear localized target of E2Fa-DPa, transcription factors controlling cell cycle progression. Required for sister chromatid cohesion and DNA repair. | protein_coding | E2F TARGET GENE 1 (ETG1) | E2F TARGET GENE 1 (ETG1) |
| AT1G30360 | AT1G30360.1 | Early-responsive to dehydration stress protein (ERD4);(source:Araport11) | protein_coding | EARLY-RESPONSIVE TO DEHYDRATION 4 (ERD4) | (OSCA3.1);EARLY-RESPONSIVE TO DEHYDRATION 4 (ERD4) |
| AT1G50940 | AT1G50940.1 | Encodes the electron transfer flavoprotein ETF alpha, a putative subunit of the mitochondrial electron transfer flavoprotein complex (ETF beta is At5g43430.1) in Arabidopsis. Mutations of the ETF beta gene results in accelerated senescence and early death compared to wild-type during extended darkness. | protein_coding | ELECTRON TRANSFER FLAVOPROTEIN ALPHA (ETFALPHA) | ELECTRON TRANSFER FLAVOPROTEIN ALPHA (ETFALPHA) |
| AT5G43430 | AT5G43430.1 | Encodes the electron transfer flavoprotein ETF beta, a putative subunit of the mitochondrial electron transfer flavoprotein complex (ETF alpha is At1g50940) in Arabidopsis. Mutations of the ETF beta gene result in accelerated senescence and early death compared to wild-type during extended darkness. Also involved in the catabolism of leucine and chlorophyll degradation pathway activated during darkness-induced carbohydrate deprivation. | protein_coding | ELECTRON TRANSFER FLAVOPROTEIN BETA (ETFBETA) | ELECTRON TRANSFER FLAVOPROTEIN BETA (ETFBETA) |
| AT5G11260 | AT5G11260.1 | Basic leucine zipper (bZIP) transcription factor. Nuclear localization. Involved in light-regulated transcriptional activation of G-box-containing promoters. Negatively regulated by Cop1. Although cytokinins do not appear to affect the gene's promoter activity, they appear to stabilize the protein. HY5 plays a role in anthocyanin accumulation in far-red light and blue light, but not in red light or in the dark. Mutant studies showed that the gene product is involved in the positive regulation of the PHYA-mediated inhibition of hypocotyl elongation. Binds to G- and Z-boxes, and other ACEs, but not to E-box. Loss of function mutation shows ABA resistant seedling phenotypes suggesting involvement for HY5 in mediating ABA responses. Binds to the promoter of ABI5 and regulates its expression. | protein_coding | ELONGATED HYPOCOTYL 5 (HY5) | ELONGATED HYPOCOTYL 5 (HY5);REVERSAL OF THE DET PHENOTYPE 5 (TED 5) |
| AT2G26830 | AT2G26830.1 | Encodes a member of a small family of choline/ethanolamine kinases that is localized to the plasma membrane. Homozygous loss of function alleles are embryo lethal. Overexpression results in altered phospholipid levels suggesting a critical role in phospholipid biosynthesis. | protein_coding | EMBRYO DEFECTIVE 1187 (emb1187) | EMBRYO DEFECTIVE 1187 (emb1187);CHOLINE/ETHANOLAMINE KINASE 4 (CEK4) |
| AT1G04635 | AT1G04635.1 | ribonuclease P family protein / Rpp14 family protein;(source:Araport11) | protein_coding | EMBRYO DEFECTIVE 1687 (EMB1687) | EMBRYO DEFECTIVE 1687 (EMB1687);A. THALIANA HOMOLOG OF YEAST POP5 (POP5); (ATPOP5) |
| AT5G08170 | AT5G08170.1 | porphyromonas-type peptidyl-arginine deiminase family protein;(source:Araport11) | protein_coding | EMBRYO DEFECTIVE 1873 (EMB1873) | AGMATINE IMINOHYDROLASE (ATAIH);EMBRYO DEFECTIVE 1873 (EMB1873) |
| AT2G41720 | AT2G41720.1 | Encodes a pentatricopeptide repeat protein that is essential for trans-splicing of a chloroplast small ribosomal subunit transcript. | protein_coding | EMBRYO DEFECTIVE 2654 (EMB2654) | EMBRYO DEFECTIVE 2654 (EMB2654) |
| AT5G02250 | AT5G02250.1 | Encodes a exoribonuclease involved in rRNA processing in mitochondria and chloroplasts.Loss of function mutations are pale green and require supplementation with sucrose for germination and early development. Plants are pale green due to defects in chloroplast biogenesis. | protein_coding | EMBRYO DEFECTIVE 2730 (EMB2730) | EMBRYO DEFECTIVE 2730 (EMB2730);ARABIDOPSIS THALIANA MITOCHONDRIAL RNASE II (ATMTRNASEII);RIBONUCLEOTIDE REDUCTASE 1 (RNR1) |
| AT2G17510 | AT2G17510.2 | ribonuclease II family protein;(source:Araport11) | protein_coding | EMBRYO DEFECTIVE 2763 (EMB2763) | EMBRYO DEFECTIVE 2763 (EMB2763);RRP44 HOMOLOG A (RRP44A); (ATRRP44A) |

| | | | | | |
|---|---|---|---|---|---|
| AT3G02660 | AT3G02660.1 | Tyrosyl-tRNA synthetase, class Ib, bacterial/mitochondrial;(source:Araport11) | protein_coding | EMBRYO DEFECTIVE 2768 (emb2768) | EMBRYONIC FACTOR 31 (FAC31);EMBRYO DEFECTIVE 2768 (emb2768) |
| AT1G67320 | AT1G67320.3 | DNA primase, large subunit family;(source:Araport11) | protein_coding | EMBRYO DEFECTIVE 2813 (EMB2813) | EMBRYO DEFECTIVE 2813 (EMB2813) |
| AT4G00620 | AT4G00620.1 | Amino acid dehydrogenase family protein;(source:Araport11) | protein_coding | EMBRYO DEFECTIVE 3127 (EMB3127) | EMBRYO DEFECTIVE 3127 (EMB3127) |
| AT4G20740 | AT4G20740.1 | Pentatricopeptide repeat (PPR-like) superfamily protein;(source:Araport11) | protein_coding | EMBRYO DEFECTIVE 3131 (EMB3131) | EMBRYO DEFECTIVE 3131 (EMB3131) |
| AT2G01860 | AT2G01860.1 | Tetratricopeptide repeat (TPR)-like superfamily protein;(source:Araport11) | protein_coding | EMBRYO DEFECTIVE 975 (EMB975) | EMBRYO DEFECTIVE 975 (EMB975) |
| AT1G16900 | AT1G16900.1 | Encodes the Arabidopsis ortholog of the yeast/human ALG9 catalyzing the luminal addition of two alpha-1,2 Man residues in assembling Glc3Man9GlcNAc2. | protein_coding | EMS-MUTAGENIZED BRI1 SUPPRESSOR 3 (EBS3) | EMS-MUTAGENIZED BRI1 SUPPRESSOR 3 (EBS3) |
| AT3G09030 | AT3G09030.1 | EAP3 is a cytolsolic BTB/POZ-domain protein involved in trafficking of PEN3. | protein_coding | ENDOPLASMIC RETICULUM-ARRESTED PEN3 (EAP3) | ENDOPLASMIC RETICULUM-ARRESTED PEN3 (EAP3) |
| AT2G19560 | AT2G19560.1 | encodes a protein with a PAM domain involved in ethylene signaling. eer5 mutants show ethylene hypersensitivity in relation to hypocotyl elongation. EER5 interacts with EIN2 and with COP9 in Y2H assays. EIN3 protein levels are the same in WT and eer5-1 mutants. EER5 may be involved in promoting a dampening of the ethylene response. | protein_coding | ENHANCED ETHYLENE RESPONSE 5 (EER5) | ENHANCED ETHYLENE RESPONSE 5 (EER5); (THP1); (AtTHP1);ECTOPIC EXPRESSION OF SEED STORAGE PROTEINS 1 (ESSP1) |
| AT1G74030 | AT1G74030.1 | Encodes the plastid-localized phosphoenolpyruvate enolase. Mutant plants have abnormal trichomes. | protein_coding | ENOLASE 1 (ENO1) | ENOLASE 1 (ENO1) |
| AT4G16210 | AT4G16210.1 | enoyl-CoA hydratase/isomerase A;(source:Araport11) | protein_coding | ENOYL-COA HYDRATASE/ISOMERASE A (ECHIA) | ENOYL-COA HYDRATASE/ISOMERASE A (ECHIA);ENOYL-COA HYDRATASE 2 (E-COAH-2) |
| AT1G70330 | AT1G70330.1 | encodes an adenosine transporter that catalyze a proton-dependent adenosine transport. The mRNA is cell-to-cell mobile. | protein_coding | EQUILIBRATIVE NUCLEOTIDE TRANSPORTER 1 (ENT1) | EQUILIBRATIVE NUCLEOTIDE TRANSPORTER 1 (ENT1);EQUILIBRATIVE NUCLEOTIDE TRANSPORTER 1 (ENT1,AT) |
| AT5G05740 | AT5G05740.1 | S2P-like putative metalloprotease, also contain transmembrane helices near their C-termini and many of them, five of seven, contain a conserved zinc-binding motif HEXXH. Homolog of EGY1. Each of the EGY1 and EGY-like proteins share two additional highly conserved motifs, the previously reported NPDG motif (aa 442?454 in EGY1, Rudner et al., 1999) and a newly defined GNLR motif (aa 171?179 in EGY1). The GNLR motif is a novel signature motif unique to EGY1 and EGY-like proteins as well as other EGY1 orthologs found in cyanobacteria. | protein_coding | ETHYLENE-DEPENDENT GRAVITROPISM-DEFICIENT AND YELLOW-GREEN-LIKE 2 (EGY2) | ETHYLENE-DEPENDENT GRAVITROPISM-DEFICIENT AND YELLOW-GREEN-LIKE 2 (EGY2); (ATEGY2) |
| AT1G17870 | AT1G17870.1 | S2P-like putative metalloprotease, also contain transmembrane helices near their C-termini and many of them, five of seven, contain a conserved zinc-binding motif HEXXH. Homolog of EGY1. Each of the EGY1 and EGY-like proteins share two additional highly conserved motifs, the previously reported NPDG motif (aa 442?454 in EGY1, Rudner et al., 1999) and a newly defined GNLR motif (aa 171?179 in EGY1). The GNLR motif is a novel signature motif unique to EGY1 and EGY-like proteins as well as other EGY1 orthologs found in cyanobacteria. | protein_coding | ETHYLENE-DEPENDENT GRAVITROPISM-DEFICIENT AND YELLOW-GREEN-LIKE 3 (EGY3) | ETHYLENE-DEPENDENT GRAVITROPISM-DEFICIENT AND YELLOW-GREEN-LIKE 3 (EGY3);ETHYLENE-DEPENDENT GRAVITROPISM-DEFICIENT AND YELLOW-GREEN-LIKE 3 (ATEGY3) |
| AT3G52940 | AT3G52940.1 | Encodes a sterol C-14 reductase required for cell division and expansion and is involved in proper organization of the embryo. | protein_coding | FACKEL (FK) | EXTRA-LONG-LIFESPAN 1 (ELL1); (HYD2);FACKEL (FK) |
| AT3G59380 | AT3G59380.1 | Encodes the alpha-subunit shared between protein farnesyltransferase and protein geranylgeranyltransferase-I. Involved in protein prenylation: covalent attachment of the C-15 isoprene farnesyl or the C-20 isoprene geranylgeranyl groups to the C-terminal end of some proteins. Involved in | protein_coding | FARNESYLTRANSFERASE A (FTA) | FARNESYLTRANSFERASE A (FTA); (PFT/PGGT-IALPHA);FARNESYLTRANSFERASE A (ATFTA);PLURIPETALA (PLP) |

| | | | | | |
|---|---|---|---|---|---|
| | | shoot and flower meristem homeostasis, and response to ABA and drought. Also regulates leaf cell shape. Mutant is epistatic to era1. | | | |
| AT5G64440 | AT5G64440.1 | AtFAAH (fatty acid amide hydrolase) modulates endogenous NAEs (N-Acylethanolamines) levels in plants by hydrolyzing NAEs to ethanolamine and their corresponding free fatty acids. NAE depletion likely participates in the regulation of plant growth. The mRNA is cell-to-cell mobile. | protein_coding | FATTY ACID AMIDE HYDROLASE (FAAH) | FATTY ACID AMIDE HYDROLASE (FAAH);FATTY ACID AMIDE HYDROLASE (AtFAAH) |
| AT4G30950 | AT4G30950.1 | Chloroplastic enzyme responsible for the synthesis of 16:2 and 18:2 fatty acids from galactolipids, sulpholipids and phosphatidylglycerol. Uses ferredoxin as electron donor. Gene mutation resulted in reduced level of unsaturated fatty acids leading to susceptibility to photoinhibition. | protein_coding | FATTY ACID DESATURASE 6 (FAD6) | FATTY ACID DESATURASE 6 (FAD6);FATTY ACID DESATURASE C (FADC);STEAROYL DESATURASE DEFICIENCY 4 (SFD4) |
| AT5G23310 | AT5G23310.1 | Fe superoxide dismutase | protein_coding | FE SUPEROXIDE DISMUTASE 3 (FSD3) | SUPEROXIDE DISMUTASE 3 (SOD3);FE SUPEROXIDE DISMUTASE 3 (FSD3) |
| AT3G20740 | AT3G20740.1 | Encodes a protein similar to the transcriptional regular of the animal Polycomb group and is involved in regulation of establishment of anterior-posterior polar axis in the endosperm and repression of flowering during vegetative phase. Mutation leads endosperm to develop in the absence of fertilization and flowers to form in seedlings and non-reproductive organs. Also exhibits maternal effect gametophytic lethal phenotype, which is suppressed by hypomethylation. Forms part of a large protein complex that can include VRN2 (VERNALIZATION 2), VIN3 (VERNALIZATION INSENSITIVE 3) and polycomb group proteins FERTILIZATION INDEPENDENT ENDOSPERM (FIE), CURLY LEAF (CLF) and SWINGER (SWN or EZA1). The complex has a role in establishing FLC (FLOWERING LOCUS C) repression during vernalization. In the ovule, the FIE transcript levels increase transiently just after fertilization. | protein_coding | FERTILIZATION-INDEPENDENT ENDOSPERM (FIE) | FERTILIZATION-INDEPENDENT ENDOSPERM 1 (FIE1); (FIS3);FERTILIZATION-INDEPENDENT ENDOSPERM (FIE) |
| AT3G54170 | AT3G54170.1 | Encodes protein that binds FKBP12. This interaction is disrupted by FK506 but not by cyclosporin A.FIP37 is a core component of the M6A methyltransferase complex. Homozygous loss of function mutations are embryo lethal. In the shoot meristem FIP37 methylation targets include WUS and STM. | protein_coding | FKBP12 INTERACTING PROTEIN 37 (FIP37) | FKBP12 INTERACTING PROTEIN 37 (FIP37);FKBP12 INTERACTING PROTEIN 37 (ATFIP37) |
| AT5G08640 | AT5G08640.1 | Encodes a flavonol synthase that catalyzes formation of flavonols from dihydroflavonols. Co-expressed with CHI and CHS (qRT-PCR). | protein_coding | FLAVONOL SYNTHASE 1 (FLS1) | (ATFLS1);FLAVONOL SYNTHASE (FLS);FLAVONOL SYNTHASE 1 (FLS1) |
| AT5G66380 | AT5G66380.1 | Encodes a folate transporter that is located in the chloroplast envelope and is able to mediate exogenous folate uptake when expressed in E. coli. However, this is not the sole folate transporter for chloroplasts as null mutants of this gene have no discernible phenotype when grown under folate-sufficient conditions and contained wild-type levels of folates in leaves. | protein_coding | FOLATE TRANSPORTER 1 (FOLT1) | FOLATE TRANSPORTER 1 (FOLT1);FOLATE TRANSPORTER 1 (ATFOLT1) |
| AT4G10260 | AT4G10260.1 | Encodes a member of the fructokinase gene family. Nomenclature according to Riggs 2017 has been adopted for the family by the community (personal communication, Boernke, Callis, Granot, Boernke, and Smeekens). | protein_coding | FRUCTOKINASE 4 (FRK4) | FRUCTOKINASE 5 (FRK5);FRUCTOKINASE 4 (FRK4) |
| AT3G54090 | AT3G54090.1 | Encodes a fructokinase-like protein (AT3G54090/FLN1, AT1G69200/FLN2), a member of the pfkB-carbohydrate kinase family. FLN1 and FLN2 are potential plastidial | protein_coding | FRUCTOKINASE-LIKE 1 (FLN1) | FRUCTOKINASE-LIKE 1 (FLN1) |

| | | | | | |
|---|---|---|---|---|---|
| | | thioredoxin z (TRX z) targets. Mutants display mutant chloroplast development, general plant growth and development defects and defects in PEP-dependent transcription. | | | |
| AT1G48270 | AT1G48270.1 | encodes a protein similar to G-coupled receptor with 7 transmembrane regions. Overexpression studies suggest this gene is involved in dormancy and flowering. Reduction of expression results in decreased sensitivity to cytokinin. | protein_coding | G-PROTEIN-COUPLED RECEPTOR 1 (GCR1) | (ATGCR1);G-PROTEIN-COUPLED RECEPTOR 1 (GCR1) |
| AT5G13630 | AT5G13630.1 | Encodes magnesium chelatase involved in plastid-to-nucleus signal transduction. | protein_coding | GENOMES UNCOUPLED 5 (GUN5) | GENOMES UNCOUPLED 5 (GUN5);CONDITIONAL CHLORINA (CCH);ABA-BINDING PROTEIN (ABAR);(CCH1);H SUBUNIT OF MG-CHELATASE (CHLH) |
| AT5G41480 | AT5G41480.1 | Encodes a dihydrofolate synthetase based on yeast complementation experiments. This protein is involved in folate biosynthesis. | protein_coding | GLOBULAR ARREST1 (GLA1) | EMBRYO DEFECTIVE 9 (EMB9);DHFS-FPGS HOMOLOG A (DFA);A. THALIANA DHFS-FPGS HOMOLOG A (ATDFA);GLOBULAR ARREST1 (GLA1) |
| AT1G09420 | AT1G09420.2 | Encodes a protein similar to glucose-6-phosphate dehydrogenase but, based on amino acid differences in the active site and lack of activity, does not encode a functional G6PDH. The amino acid sequence for the consensus sequence of the G6PDH active site (DHYLGKE) differs in three places in this protein. gc exon splice site at 20574 is based on protein alignment, and is not confirmed experimentally. | protein_coding | GLUCOSE-6-PHOSPHATE DEHYDROGENASE 4 (G6PD4) | GLUCOSE-6-PHOSPHATE DEHYDROGENASE 4 (G6PD4) |
| AT5G64050 | AT5G64050.1 | Glutamate-tRNA ligase. Targeted to mitochondria and chloroplast. Its inactivation causes developmental arrest of chloroplasts and mitochondria in Nicotiana benthamiana. | protein_coding | GLUTAMATE TRNA SYNTHETASE (ERS) | ETHYLENE RESPONSE SENSOR (ERS);GLUTAMATE TRNA SYNTHETASE (ERS);OVULE ABORTION 3 (OVA3);(ATERS) |
| AT1G42970 | AT1G42970.1 | Encodes chloroplast localized glyceraldehyde-3-phosphate dehydrogenase that can use both NADH and NADPH to reduce 1,3-diphosphate glycerate. It forms A2B2 heterotetramers with GapA forms of the GADPH enzyme. These complexes are active in the light under reducing conditions, but show reduced NADPH-dependent activity in response to oxidized thioredoxins and increased NAD(H)/NADP(H) ratios due to the formation of inactive A8B8 hexadecamers. The mRNA is cell-to-cell mobile. | protein_coding | GLYCERALDEHYDE-3-PHOSPHATE DEHYDROGENASE B SUBUNIT (GAPB) | GLYCERALDEHYDE-3-PHOSPHATE DEHYDROGENASE B SUBUNIT (GAPB) |
| AT2G13100 | AT2G13100.1 | Encodes a member of the phosphate starvation-induced glycerol-3-phosphate permease gene family: AT3G47420(G3Pp1), AT4G25220(G3Pp2), AT1G30560(G3Pp3), AT4G17550(G3Pp4) and AT2G13100(G3Pp5). The mRNA is cell-to-cell mobile. | protein_coding | GLYCEROL-3-PHOSPHATE PERMEASE 5 (G3Pp5) | GLYCEROL-3-PHOSPHATE PERMEASE 5 (AtG3Pp5);GLYCEROL-3-PHOSPHATE PERMEASE 5 (G3Pp5) |
| AT3G25530 | AT3G25530.1 | Encodes gamma-hydroxybutyrate dehydrogenase (AtGHBDH). Contains a NADP-binding domain. GHBDH is proposed to function in oxidative stress tolerance. | protein_coding | GLYOXYLATE REDUCTASE 1 (GLYR1) | (GHBDH); (ATGHBDH); (ATGLYR1);GLYOXYLATE REDUCTASE 1 (GR1);GLYOXYLATE REDUCTASE 1 (GLYR1) |
| AT3G27530 | AT3G27530.1 | This gene is predicted to encode a protein that functions as a Golgi apparatus structural component known as a golgin in mammals and yeast. A fluorescently-tagged version of GC6 co-localizes with Golgi markers, and this localization appears to be replicated using the C-terminal (225 aa) portion of the protein. | protein_coding | GOLGIN CANDIDATE 6 (GC6) | MAIGO 4 (MAG4);GOLGIN CANDIDATE 6 (GC6) |
| AT4G34460 | AT4G34460.1 | Encodes the heterotrimeric G-protein beta subunit and is involved in organ shape. A significant fraction of the protein is found in the ER. Mutants carrying null alleles express similar fruit phenotypes, as seen in er plants, but differ from er in that the stem is only slightly shorter than that in the wild type, the pedicel is slightly longer than that in the wild type, and the leaves are rounder than those in er mutants. Gene is expressed in all tissues examined, with highest | protein_coding | GTP BINDING PROTEIN BETA 1 (AGB1) | GTP BINDING PROTEIN BETA 1 (AGB1);(ATAGB1);ERECTA-LIKE 4 (ELK4) |

| | | | | | |
|---|---|---|---|---|---|
| | | expression level found in siliques. It is involved in resistance to Plectosphaerella cucumerina. The predicted protein has two DWD motifs. It can bind to DDB1a in Y2H assays and may be involved in the formation of a CUL4-based E3 ubiquitin ligase. It seems to be involved in the calcium-mediated response to extracellular ATP. | | | |
| AT1G01910 | AT1G01910.1 | P-loop containing nucleoside triphosphate hydrolases superfamily protein;(source:Araport11) | protein_coding | GUIDED ENTRY OF TAIL-ANCHORED PROTEIN 3A (GET3A) | GUIDED ENTRY OF TAIL-ANCHORED PROTEIN 3A (GET3A);GUIDED ENTRY OF TAIL-ANCHORED PROTEIN 3A (ATGET3A) |
| AT5G63220 | AT5G63220.1 | golgi-to-ER traffic-like protein;(source:Araport11) | protein_coding | GUIDED ENTRY OF TAIL-ANCHORED PROTEINS 4 (GET4) | GUIDED ENTRY OF TAIL-ANCHORED PROTEINS 4 (ATGET4);GUIDED ENTRY OF TAIL-ANCHORED PROTEINS 4 (GET4) |
| AT3G05040 | AT3G05040.1 | Encodes member of importin/exportin family. Involved in timing of shoot maturation. Involved in miRNA transport. Mutants flower early and have small, curled leaves and reduced abundance of certain miRNA species. | protein_coding | HASTY (HST) | HASTY (HST);HASTY 1 (HST1) |
| AT1G47840 | AT1G47840.1 | Encodes a putative hexokinase. | protein_coding | HEXOKINASE 3 (HXK3) | HEXOKINASE 3 (HXK3) |
| AT5G52110 | AT5G52110.1 | chaperone (DUF2930);(source:Araport11) | protein_coding | HIGH CHLOROPHYLL FLUORESCENCE 208 (HCF208) | COFACTOR ASSEMBLY OF COMPLEX C (CCB2);HIGH CHLOROPHYLL FLUORESCENCE 208 (HCF208) |
| AT5G36170 | AT5G36170.1 | Required for normal processing of polycistronic plastidial transcripts | protein_coding | HIGH CHLOROPHYLL FLUORESCENT 109 (HCF109) | (ATPRFB);HIGH CHLOROPHYLL FLUORESCENT 109 (HCF109) |
| AT1G66080 | AT1G66080.1 | Encodes a glucose-regulated protein that binds to the promoters of glucose-regulated heat shock responsive genes and promotes chromatin acetylation. HLP1 is required in maintaining histone H3K acetylation and H3K4 methylation marks at the promoters of heat shock protein genes in providing thermotolerance/thermomemory response. | protein_coding | HIKESHI-LIKE PROTEIN1 (HLP1) | HIKESHI-LIKE PROTEIN1 (HLP1);HIKESHI-LIKE (HKL) |
| AT5G63890 | AT5G63890.2 | Encodes histidinol dehydrogenase. Up-regulated in response to UV-B. | protein_coding | HISTIDINOL DEHYDROGENASE (HDH) | HISTIDINOL DEHYDROGENASE (ATHDH);HISTIDINOL DEHYDROGENASE (HDH);HISTIDINE BIOSYNTHESIS 8 (HISN8) |
| AT5G56740 | AT5G56740.1 | Encodes an enzyme with histone acetyltransferase activity. Histone H4 is the primary substrate for the enzyme. Prior acetylation of lysine 12 of histone H4 reduces radioactive acetylation by HAG2. HAG2 acetylates histone H4 lysine 12. | protein_coding | HISTONE ACETYLTRANSFERASE OF THE GNAT FAMILY 2 (HAG2) | HISTONE ACETYLTRANSFERASE OF THE GNAT FAMILY 2 (HAG2); (HAC07); (HAG02); (HAC7) |
| AT4G33470 | AT4G33470.1 | Encodes HDA14, a member of the histone deacetylase family proteins that can deacetylate a-tubulin, associates with a/b-tubulin and is retained on GTP/taxol-stabilized microtubules, at least in part, by direct association with the PP2A-A2 subunit. The association of a histone deacetylase with PP2A suggests a direct link between protein phosphorylation and acetylation. | protein_coding | HISTONE DEACETYLASE 14 (hda14) | HISTONE DEACETYLASE 14 (hda14); (ATHDA14);HISTONE DEACETYLASE 14 (HDAC14); (ATHDAC14) |
| AT1G08460 | AT1G08460.1 | histone deacetylase 8;(source:Araport11) | protein_coding | HISTONE DEACETYLASE 8 (HDA08) | HISTONE DEACETYLASE 8 (HDA08); (ATHDA8);HISTONE DEACETYLASE 8 (HDA8) |
| AT3G44680 | AT3G44680.1 | Encodes HDA9 (a RPD3-like histone deacetylase). Functions in promoting the onset of leaf senescence.The hda9 mutant shows enhanced H3K9 acetylation levels,based on immunodetection using H3K9ac antibodies. | protein_coding | HISTONE DEACETYLASE 9 (HDA9) | HISTONE DEACETYLASE 9 (HDAC9); (ATHDAC9);HISTONE DEACETYLASE 9 (HDA9); (HDA09); (ATHDA9) |
| AT4G20930 | AT4G20930.1 | Encodes a 3-hydroxyisobutyrate dehydrogenase. | protein_coding | HYDROXYISOBUTYRATE DEHYDROGENASE 1 (HDH1) | HYDROXYISOBUTYRATE DEHYDROGENASE 1 (HDH1) |
| AT5G25265 | AT5G25265.1 | Hyp O-arabinosyltransferase-like protein;(source:Araport11) | protein_coding | HYDROXYPROLINE O-ARABINOSYLATRANSFERASE 1 (HPAT1) | HYDROXYPROLINE O-ARABINOSYLATRANSFERASE 1 (HPAT1) |
| AT1G68010 | AT1G68010.2 | Encodes hydroxypyruvate reductase. | protein_coding | HYDROXYPYRUVATE REDUCTASE (HPR) | HYDROXYPYRUVATE REDUCTASE (HPR) |

| | | | | | |
|---|---|---|---|---|---|
| AT1G71750 | AT1G71750.2 | Encodes a protein with hypoxanthine-guanine-phosphoribosyltransferase activity. Unlike some related enzymes, it does not appear to act on xanthine in vitro. The enzyme catalyzes reactions occurring in both directions, but appears to prefer acting on guanine, followed by hypoxanthine, in vitro. The enzyme is likely to function in purine salvage pathways and appears to be important for seed germination. | protein_coding | HYPOXANTHINE-GUANINE PHOSPHORIBOSYLTRANSFERASE (HGPT) | HYPOXANTHINE-GUANINE PHOSPHORIBOSYLTRANSFERASE (HGPT) |
| AT1G68100 | AT1G68100.1 | member of IAA-alanine resistance protein 1 | protein_coding | IAA-ALANINE RESISTANT 1 (IAR1) | IAA-ALANINE RESISTANT 1 (IAR1) |
| AT1G44350 | AT1G44350.1 | encodes a protein similar to IAA amino acid conjugate hydrolase. | protein_coding | IAA-LEUCINE RESISTANT (ILR)-LIKE GENE 6 (ILL6) | IAA-LEUCINE RESISTANT (ILR)-LIKE GENE 6 (ILL6) |
| AT5G54140 | AT5G54140.1 | encodes a protein similar to IAA amino acid conjugate hydrolase | protein_coding | IAA-LEUCINE-RESISTANT (ILR1)-LIKE 3 (ILL3) | IAA-LEUCINE-RESISTANT (ILR1)-LIKE 3 (ILL3) |
| AT5G03070 | AT5G03070.1 | Putative importin alpha isoform. When overexpressed can rescue the impa-4 decreased transformation susceptibility phenotype. | protein_coding | IMPORTIN ALPHA ISOFORM 9 (IMPA-9) | IMPORTIN ALPHA ISOFORM 9 (IMPA-9) |
| AT5G26820 | AT5G26820.1 | Mutations in MAR1 confer resistance, while MAR1 overexpression causes hypersensitivity to multiple aminoglycoside antibiotics. Localizes to the chloroplast envelope. MAR1 may act as a plastid transporter involved in cellular iron homeostasis. The mRNA is cell-to-cell mobile. | protein_coding | IRON-REGULATED PROTEIN 3 (IREG3) | MULTIPLE ANTIBIOTIC RESISTANCE 1 (MAR1);IRON REGULATED 3 (IREG3); (RTS3) |
| AT2G39930 | AT2G39930.1 | Encodes an isoamylase-type debranching enzyme. Mutations in this gene cause the loss of detectable isoamylase activity and the disruption of normal starch structure. Mutants have reduced starch content and abnormally structured amylopectins and phytoglycogens. It has been postulated that AtISA1 interacts with AtISA2 to form the Iso1 complex. | protein_coding | ISOAMYLASE 1 (ISA1) | ARABIDOPSIS THALIANA ISOAMYLASE 1 (ATISA1);ISOAMYLASE 1 (ISA1) |
| AT4G09020 | AT4G09020.1 | Encodes an isoamylase-like protein. Mutant studies show that the gene is strongly involved in starch breakdown. A GUS-protein fusion product was shown to localize to the surface of chloroplastic structures reminiscent of starch granules. In the mutants, the chloroplastic α-amylase AMY3 is upregulated. The mRNA is cell-to-cell mobile. | protein_coding | ISOAMYLASE 3 (ISA3) | ISOAMYLASE 3 (ISA3); (ATISA3) |
| AT5G20040 | AT5G20040.3 | Encodes tRNA isopentenyltransferase AtIPT9. | protein_coding | ISOPENTENYLTRANSFERASE 9 (IPT9) | ISOPENTENYLTRANSFERASE 9 (IPT9) |
| AT3G45300 | AT3G45300.1 | Encodes isovaleryl-coenzyme a dehydrogenase. Mutants have increases in 12 seed free amino acids, accumulation of seed homomethionine and 3-isovaleroyloxypropyl-glucosinolate, with a concomitant decrease in seed 3-benzoyloxypropyl-glucosinolate. The mRNA is cell-to-cell mobile. | protein_coding | ISOVALERYL-COA-DEHYDROGENASE (IVD) | ISOVALERYL-COA-DEHYDROGENASE (IVD);ISOVALERYL-COA-DEHYDROGENASE (IVDH); (ATIVD) |
| AT5G03770 | AT5G03770.1 | Encodes a putative KDO (3-deoxy-D-manno-octulosonate) transferase | protein_coding | KDO TRANSFERASE A (KDTA) | (ATKDTA);KDO TRANSFERASE A (KDTA) |
| AT5G14760 | AT5G14760.1 | At5g14760 encodes for L-aspartate oxidase involved in the early steps of NAD biosynthesis. In contrary to the EC 1.4.3.16 (l-aspartate oxidase - deaminating) the enzyme catalyzes the reaction L-aspartate + O2 = iminoaspartate (alpha-iminosuccinate) + H2O2 | protein_coding | L-ASPARTATE OXIDASE (AO) | FLAGELLIN-INSENSITIVE 4 (FIN4);L-ASPARTATE OXIDASE (AO) |
| AT3G19260 | AT3G19260.1 | LAG1 homolog. Loss of function mutant is sensitive to AAL-toxin. LOH2 is presumed to function in sphingolipid metabolism. It encodes a ceramide synthase essential for production of LCFA-ceramides (mainly C16). | protein_coding | LAG1 HOMOLOGUE 2 (LOH2) | LAG1 HOMOLOGUE 2 (LOH2);LONGEVITY ASSURANCE GENE1 HOMOLOG 2 (LAG1 HOMOLOG 2) |
| AT2G40190 | AT2G40190.1 | Encodes a putative alpha-1,2-mannosyltransferase in N-linked glycoprotein (homologous to yeast ALG11). Plays | protein_coding | LEAF WILTING 3 (LEW3) | LEAF WILTING 3 (LEW3) |

| | | | | | |
|---|---|---|---|---|---|
| | | vital roles in cell-wall biosynthesis and abiotic stress response. Located in endoplasmic reticulum membrane. | | | |
| AT1G02050 | AT1G02050.1 | Chalcone and stilbene synthase family protein;(source:Araport11) | protein_coding | LESS ADHESIVE POLLEN 6 (LAP6) | POLYKETIDE SYNTHASE A (PKSA);LESS ADHESIVE POLLEN 6 (LAP6) |
| AT1G76570 | AT1G76570.1 | Chlorophyll A-B binding family protein;(source:Araport11) | protein_coding | LIGHT-HARVESTING COMPLEX B7 (LHCB7) | LIGHT-HARVESTING COMPLEX B7 (LHCB7); (ATLHCB7) |
| AT3G55760 | AT3G55760.1 | hypothetical protein;(source:Araport11) | protein_coding | LIKE EARLY STARVATION (LESV) | LIKE EARLY STARVATION (LESV) |
| AT5G04360 | AT5G04360.1 | Encodes an enzyme thought to be involved in the hydrolysis of the α-1,6 linkages during starch degradation in seed endosperm. However, a knockout mutant of Arabidopsis lacking limit dextrinase has normal rates of starch degradation in the leaf at night, indicating that more than one isoamylases might be involved in this process. | protein_coding | LIMIT DEXTRINASE (LDA) | PULLULANASE 1 (ATPU1);LIMIT DEXTRINASE (LDA);PULLULANASE 1 (PU1);LIMIT DEXTRINASE (ATLDA) |
| AT2G04560 | AT2G04560.1 | transferases, transferring glycosyl groups;(source:Araport11) | protein_coding | LIPID X B (LPXB) | LIPID X B (LPXB); (ATLPXB) |
| AT1G04640 | AT1G04640.1 | Lipoyltransferase, located in mitochondria but not found in chloroplasts | protein_coding | LIPOYLTRANSFERASE 2 (LIP2) | LIPOYLTRANSFERASE 2 (LIP2) |
| AT5G11950 | AT5G11950.1 | Encodes a protein of unknown function. It has been crystallized and shown to be structurally almost identical to the protein encoded by At2G37210. | protein_coding | LONELY GUY 8 (LOG8) | (MOBP2);LONELY GUY 8 (LOG8) |
| AT2G46090 | AT2G46090.1 | Encodes a putative sphingosine kinase (SphK) containing the five conserved domains (C1-C5) previously identified in SphKs. | protein_coding | LONG-CHAIN BASE (LCB) KINASE 2 (LCBK2) | LONG-CHAIN BASE (LCB) KINASE 2 (LCBK2) |
| AT1G02910 | AT1G02910.1 | Mutants defective in this gene were shown to have a reduced PSII content (overall reduction in the levels of several PSII subunits) and a disrupted grana stack structure. The N-terminal half of the protein contains two tetratricopeptide repeat (TPR) motifs that are arranged tandemly, each consisting of a 34-residue degenerate consensus sequence. The N-terminal sequence is rich in positive and hydroxylated amino acid residues. | protein_coding | LOW PSII ACCUMULATION1 (LPA1) | LOW PSII ACCUMULATION1 (LPA1) |
| AT5G47010 | AT5G47010.1 | Required for nonsense-mediated mRNA decay. Involved in RNA interference. lba1 mutants has reduced sugar-induced expression of Atb- amylase, is hypersensitive to glucose and abscisic acid and resistant to mannose, and shows early flowering, short day-sensitive growth, and seed germination phenotypes. The mRNA is cell-to-cell mobile. | protein_coding | LOW-LEVEL BETA-AMYLASE 1 (LBA1) | (UPF1); (ATUPF1);LOW-LEVEL BETA-AMYLASE 1 (LBA1) |
| AT3G53130 | AT3G53130.1 | Lutein-deficient 1 (LUT1) required for lutein biosynthesis, member of the xanthophyll class of carotenoids. Involved in epsilon ring hydroxylation. Maps at 67.3 cM on chromosome 3. | protein_coding | LUTEIN DEFICIENT 1 (LUT1) | CYTOCHROME P450 97C1 (CYP97C1);LUTEIN DEFICIENT 1 (LUT1) |
| AT3G10230 | AT3G10230.1 | Encodes a protein with lycopene β-cyclase activity. This enzyme uses the linear, symmetrical lycopene as substrate. However, unlike the ε-cyclase which adds only one ring, the β-cyclase introduces a ring at both ends of lycopene to form the bicyclic β-carotene. | protein_coding | LYCOPENE CYCLASE (LYC) | LYCOPENE CYCLASE (LYC);SUPPRESSOR OF ZEAXANTHIN-LESS 1 (SZL1); (AtLCY) |
| AT1G51940 | AT1G51940.1 | Encodes a LysM-containing receptor-like kinase. Induction of chitin-responsive genes by chitin treatment is not blocked in the mutant. Based on protein sequence alignment analysis, it has a typical RD signaling domain in its catalytic loop and possesses autophosphorylation activity.It is required for the suppression of defense responses in absence of pathogen infection or upon abscisic acid treatment. Loss-of-function mutants display enhanced resistance to Botrytis cinerea and Pectobacterium carotovorum. Its expression is repressed by | protein_coding | LYSM-CONTAINING RECEPTOR-LIKE KINASE 3 (LYK3) | (ATLYK3);LYSM-CONTAINING RECEPTOR-LIKE KINASE 3 (LYK3) |

| | | | | | |
|---|---|---|---|---|---|
| | | pathogen infection and biological elicitors and is induced abscisic acid.Expression is strongly repressed by elicitors and fungal infection, and is induced by the hormone abscisic acid (ABA). Insertional mutants show increased expression of PHYTOALEXIN-DEFICIENT 3 (PAD3), enhanced resistance to Botrytis cinerea and Pectobacterium carotovorum infection and reduced physiological responses to ABA, suggesting that LYK3 is important for the cross-talk between signaling pathways activated by ABA and pathogens (PMID:24639336). | | | |
| AT4G25080 | AT4G25080.3 | Encodes a protein with methyltransferase activity responsible for the methylation of magnesium protoporphyrin IX. Mutants defective in this gene are affected in chlorophyll biosynthesis and show a reduction in the accumulation of a number of major thylakoid-associated proteins including components of PSI (LHCI), PSII (LHCII, D1, CP43) and the cytochrome b6f complex (Cytf). By contrast, no significant changes were detected for the proteins of the stroma and the chloroplast envelope. | protein_coding | MAGNESIUM-PROTOPORPHYRIN IX METHYLTRANSFERASE (CHLM) | MAGNESIUM-PROTOPORPHYRIN IX METHYLTRANSFERASE (CHLM) |
| AT1G08660 | AT1G08660.1 | Encodes a sialyltransferase-like protein that is localized to the Golgi apparatus and is involved in pollen tube growth and pollen germination. | protein_coding | MALE GAMETOPHYTE DEFECTIVE 2 (MGP2) | SIALYLTRANSFERASE-LIKE 1 (SIA1);MALE GAMETOPHYTE DEFECTIVE 2 (MGP2) |
| AT1G27520 | AT1G27520.1 | Glycosyl hydrolase family 47 protein;(source:Araport11) | protein_coding | MANNOSIDASE 5 (MNS5) | MANNOSIDASE 5 (MNS5) |
| AT5G56580 | AT5G56580.1 | Encodes a member of the MAP Kinase Kinase family of proteins. It can phosphorylate MPK12 in vitro and it can be dephosphorylated by MKP2 in vitro. | protein_coding | MAP KINASE KINASE 6 (MKK6) | ARABIDOPSIS THALIANA MAP KINASE KINASE 6 (ATMKK6);SUPPRESSOR OF MKK1 MKK2 4 (SUMM4);MAP KINASE KINASE 6 (MKK6);ARABIDOPSIS NQK1 (ANQ1) |
| AT5G20170 | AT5G20170.1 | RNA polymerase II transcription mediator;(source:Araport11) | protein_coding | MEDIATOR 17 (MED17) | MEDIATOR 17 (MED17) |
| AT3G09180 | AT3G09180.1 | mediator of RNA polymerase II transcription subunit;(source:Araport11) | protein_coding | MEDIATOR 3 (MED3) | MEDIATOR 3 (MED3) |
| AT5G54260 | AT5G54260.1 | DNA repair and meiotic recombination protein, component of MRE11 complex with RAD50 and NBS1 | protein_coding | MEIOTIC RECOMBINATION 11 (MRE11) | MEIOTIC RECOMBINATION 11 (MRE11);ARABIDOPSIS MEIOTIC RECOMBINATION 11 (ATMRE11) |
| AT3G12100 | AT3G12100.1 | Cation efflux family protein;(source:Araport11) | protein_coding | METAL TRANSPORT PROTEIN 5 (MTP5) | (ATMTP5);METAL TRANSPORT PROTEIN 5 (MTP5) |
| AT4G37040 | AT4G37040.1 | encodes a methionine aminopeptidase | protein_coding | METHIONINE AMINOPEPTIDASE 1D (MAP1D) | METHIONINE AMINOPEPTIDASE 1D (MAP1D) |
| AT2G16440 | AT2G16440.1 | Regulates DNA replication via interaction with BICE1 and MCM7. | protein_coding | MINICHROMOSOME MAINTENANCE 4 (MCM4) | MINICHROMOSOME MAINTENANCE 4 (MCM4) |
| AT5G44635 | AT5G44635.1 | minichromosome maintenance (MCM2/3/5) family protein;(source:Araport11) | protein_coding | MINICHROMOSOME MAINTENANCE 6 (MCM6) | MINICHROMOSOME MAINTENANCE 6 (MCM6) |
| AT5G42130 | AT5G42130.1 | Encodes a protein belonging to the mitochondrial carrier family and similar to animal mitoferrin but likely NOT to be located in the mitochondria, but rather in chloroplasts. It is likely to be involved in transporting iron into the chloroplast. | protein_coding | MITOFERRINLIKE1 (Mfl1) | (AtMfl1);MITOFERRINLIKE1 (Mfl1) |
| AT3G25980 | AT3G25980.1 | Encodes MAD2 (MITOTIC ARREST-DEFICIENT 2). May have the spindle assembly checkpoint protein functions conserved from yeast to humans. | protein_coding | MITOTIC ARREST-DEFICIENT 2 (MAD2) | MITOTIC ARREST-DEFICIENT 2 (MAD2); (ATMAD2) |
| AT3G18165 | AT3G18165.1 | Encodes MOS4 (Modifier of snc1, 4), a nuclear protein homologous to human Breast Cancer-Amplified Sequence (BCAS2). MOS4 interacts with AtCDC5 and PRL1. All three proteins are essential for plant innate immunity. | protein_coding | MODIFIER OF SNC1,4 (MOS4) | MODIFIER OF SNC1,4 (MOS4) |
| AT2G28390 | AT2G28390.1 | SAND family protein;(source:Araport11) | protein_coding | MONENSIN SENSITIVITY1 (MON1) | MONENSIN SENSITIVITY1 (MON1) |

| | | | | | |
|---|---|---|---|---|---|
| AT4G10760 | AT4G10760.1 | Encodes a member of a core set of mRNA m6A writer proteins and is required for N6-adenosine methylation of mRNA. | protein_coding | MRNAADENOSINE METHYLASE (MTA) | MRNAADENOSINE METHYLASE (MTA);EMBRYO DEFECTIVE 1706 (EMB1706) |
| AT1G31190 | AT1G31190.1 | Encodes a myo-inositol monophosphatase IMPL1 (myo-Inositol monophosphatase like 1). | protein_coding | MYO-INOSITOL MONOPHOSPHATASE LIKE 1 (IMPL1) | MYO-INOSITOL MONOPHOSPHATASE LIKE 1 (IMPL1) |
| AT4G04880 | AT4G04880.1 | adenosine/AMP deaminase family protein;(source:Araport11) | protein_coding | N6-METHYL AMP DEAMINASE (MAPDA) | N6-METHYL AMP DEAMINASE (MAPDA);AMINOHYDROLASE (ADAL) |
| AT1G78590 | AT1G78590.1 | Encodes a NADH kinase which can synthesize NADPH from NADH; also utilizes NAD+ as substrate although NADH is the preferred substrate. | protein_coding | NAD(H) KINASE 3 (NADK3) | ARABIDOPSIS THALIANA NADH KINASE 3 (ATNADK-3);NAD(H) KINASE 3 (NADK3) |
| AT5G08740 | AT5G08740.1 | Encodes a dedicated Type II NADPH dehydrogenase that catalyzes the penultimate step in phylloquinone (vitamin K1). biosynthesis | protein_coding | NAD(P)H DEHYDROGENASE C1 (NDC1) | NAD(P)H DEHYDROGENASE C1 (NDC1) |
| AT4G15545 | AT4G15545.1 | NAI1 interacting protein, involved in ER body formation. | protein_coding | NAI2-INTERACTING PROTEIN 1 (NAIP1) | NAI2-INTERACTING PROTEIN 1 (NAIP1) |
| AT3G52640 | AT3G52640.2 | Encodes a gamma-secretase subunit. Associates with other subunits in intracellular membrane compartments. | protein_coding | NCT (NCT) | NCT (NCT) |
| AT3G51050 | AT3G51050.1 | NERD1 is a single copy locus encoding a protein of unknown function that is localized to the nucleus. Single mutants show defects in root hair growth, root meristem function, cell elongation. NERD1 appears to act synergistically with the exocyst in root development. | protein_coding | NEW ENHANCER OF ROOT DWARFISM1 (NERD1) | NEW ENHANCER OF ROOT DWARFISM1 (NERD1) |
| AT3G53140 | AT3G53140.1 | Nicotinate N-methyltransferase involved in N-methylnicotinate formation. | protein_coding | NICOTINATE N-METHYLTRANSFERASE (NANMT) | NICOTINATE N-METHYLTRANSFERASE (NANMT) |
| AT5G65720 | AT5G65720.1 | Encodes a cysteine desulfurase whose activity is dependent on AtSufE activation. It requires pyridoxal phosphate (PLP) for proper folding. Its catalytic efficiency is increase three-fold in the presence of AtFH (frataxin). | protein_coding | NITROGEN FIXATION S (NIFS)-LIKE 1 (NFS1) | NITROGEN FIXATION S HOMOLOG 1 (ATNIFS1);ARABIDOPSIS THALIANA NITROGEN FIXATION S (NIFS)-LIKE 1 (ATNFS1);NITROGEN FIXATION S (NIFS)-LIKE 1 (NFS1);NITROGEN FIXATION S HOMOLOG 1 (NIFS1) |
| AT3G47450 | AT3G47450.1 | Encodes a protein with similarity to the bacterial YqeH GTPase required for proper ribosome assembly. Mutant analyses show that this protein regulates growth and hormonal signaling and attenuates oxidative stress and reactive oxygen species (ROS). It also seems to be involved in regulating leaf senescence, cell death, nitric oxide biosynthesis in response to ABA but not exogenous H2O2. This protein also appears to be required for proper plastid biogenesis. Levels of several plastid-localized proteins, including RBCL, ClpP1, and the MEP biosynthesis enzymes DXS and DXR are altered in rif1-1 mutants. This protein was originally characterized as a mitochondrial-localized nitric oxide synthase, but, the synthase activity was later disproven. In addition, new studies with GFP fusion proteins and chloroplast import assays suggest that this protein is found in chloroplasts. Its localization to the chloroplast is enhanced by S-acylation. | protein_coding | NO ASSOCIATED 1 (NOA1) | SUPPRESSOR OF VARIEGATION 10 (SVR10);NO ASSOCIATED 1 (NOA1); (ATNOA1);NITRIC OXIDE SYNTHASE 1 (NOS1); (ATNOS1);RESISTANT TO INHIBITION WITH FOSMIDOMYCIN 1 (RIF1) |
| AT5G13390 | AT5G13390.1 | Required for normal pollen development and lipid accumulation within the tapetum | protein_coding | NO EXINE FORMATION 1 (NEF1) | NO EXINE FORMATION 1 (NEF1) |
| AT1G07230 | AT1G07230.1 | non-specific phospholipase C1;(source:Araport11) | protein_coding | NON-SPECIFIC PHOSPHOLIPASE C1 (NPC1) | NON-SPECIFIC PHOSPHOLIPASE C1 (NPC1) |
| AT4G13250 | AT4G13250.1 | Encodes a chlorophyll b reductase involved in the degradation of chlorophyll b and LHCII (light harvesting complex II). | protein_coding | NON-YELLOW COLORING 1 (NYC1) | NON-YELLOW COLORING 1 (NYC1) |
| AT5G18110 | AT5G18110.1 | Putative cap-binding protein;(source:Araport11) | protein_coding | NOVEL CAP-BINDING PROTEIN (NCBP) | NOVEL CAP-BINDING PROTEIN (NCBP) |

| | | | | | |
|---|---|---|---|---|---|
| AT5G11240 | AT5G11240.1 | GHS40 encodes a WD40 protein, that is localized in the nucleus and nucleolus. In the presence of high glucose it negatively regulates the expression of abscisic acid degradation and signaling genes. | protein_coding | NUCLEAR GLUCOSE-RESPONSIVE WD40 PROTEIN1 (NUGWD1) | NUCLEAR GLUCOSE-RESPONSIVE WD40 PROTEIN1 (NUGWD1);GLUCOSE HYPERSENSITIVE 40 (GHS40); (ATGHS40) |
| AT1G06790 | AT1G06790.1 | Encodes a subunit of RNA polymerase III involved in maintaining global RNA homeostasis, not just that of genes transcribed by RNA pol III. | protein_coding | NUCLEAR RNA POLYMERASE C, SUBUNIT 7 (NRPC7) | NUCLEAR RNA POLYMERASE C, SUBUNIT 7 (NRPC7) |
| AT5G20070 | AT5G20070.1 | nudix hydrolase homolog 19;(source:Araport11) | protein_coding | NUDIX HYDROLASE HOMOLOG 19 (NUDX19) | ARABIDOPSIS THALIANA NUDIX HYDROLASE HOMOLOG 19 (atnudt19);NUDIX HYDROLASE HOMOLOG 19 (NUDX19);NUDIX HYDROLASE HOMOLOG 19 (ATNUDX19) |
| AT5G47240 | AT5G47240.1 | nudix hydrolase homolog 8;(source:Araport11) | protein_coding | NUDIX HYDROLASE HOMOLOG 8 (NUDT8) | NUDIX HYDROLASE HOMOLOG 8 (atnudt8);NUDIX HYDROLASE HOMOLOG 8 (NUDT8);NUDIX HYDROLASE HOMOLOG 8 (NUDX8) |
| AT5G53450 | AT5G53450.1 | OBP3-responsive protein 1;(source:Araport11) | protein_coding | OBP3-RESPONSIVE GENE 1 (ORG1) | OBP3-RESPONSIVE GENE 1 (ORG1) |
| AT1G76400 | AT1G76400.1 | Ribophorin I;(source:Araport11) | protein_coding | OLIGOSACCHARYLTRANSFERASE 1B (OST1B) | OLIGOSACCHARYLTRANSFERASE 1B (OST1B) |
| AT3G57430 | AT3G57430.1 | Encodes a chloroplast RNA editing factor. | protein_coding | ORGANELLE TRANSCRIPT PROCESSING 84 (OTP84) | ORGANELLE TRANSCRIPT PROCESSING 84 (OTP84) |
| AT2G35720 | AT2G35720.1 | Encodes OWL1, a J-domain protein involved in perception of very low light fluences. | protein_coding | ORIENTATION UNDER VERY LOW FLUENCES OF LIGHT 1 (OWL1) | ORIENTATION UNDER VERY LOW FLUENCES OF LIGHT 1 (OWL1) |
| AT1G75330 | AT1G75330.1 | ornithine carbamoyltransferase;(source:Araport11) | protein_coding | ORNITHINE CARBAMOYLTRANSFERASE (OTC) | ORNITHINE CARBAMOYLTRANSFERASE (OTC) |
| AT5G46180 | AT5G46180.1 | Encodes an ornithine delta-aminotransferase that is transcriptionally up-regulated in young seedlings and in response to salt stress. It is unlikely to play a role in salt-stress-induced proline accumulation, however, it appears to participate in arginine and ornithine catabolism. | protein_coding | ORNITHINE-DELTA-AMINOTRANSFERASE (DELTA-OAT) | ORNITHINE-DELTA-AMINOTRANSFERASE (DELTA-OAT) |
| AT3G55400 | AT3G55400.1 | methionyl-tRNA synthetase / methionine-tRNA ligase / MetRS (cpMetRS);(source:Araport11) | protein_coding | OVULE ABORTION 1 (OVA1) | OVULE ABORTION 1 (OVA1) |
| AT2G25840 | AT2G25840.2 | Nucleotidylyl transferase superfamily protein;(source:Araport11) | protein_coding | OVULE ABORTION 4 (OVA4) | OVULE ABORTION 4 (OVA4) |
| AT5G52520 | AT5G52520.1 | Encodes a chloroplast and mitochondria localized prolyl-tRNA synthetase. | protein_coding | OVULE ABORTION 6 (OVA6) | OVULE ABORTION 6 (OVA6); (ATPRORS-ORG);PROLYL-TRNA SYNTHETASE ORGANELLAR (PRORS-ORG);PROLYL-TRNA SYNTHETASE 1 (PRORS1) |
| AT1G32520 | AT1G32520.1 | TLDc domain protein;(source:Araport11) | protein_coding | OXIDATION RESISTANCE 4 (OXR4) | OXIDATION RESISTANCE 4 (OXR4) |
| AT5G39590 | AT5G39590.1 | TLD-domain containing nucleolar protein;(source:Araport11) | protein_coding | OXIDATION RESISTANCE 5 (OXR5) | OXIDATION RESISTANCE 5 (OXR5) |
| AT2G06050 | AT2G06050.2 | Encodes a 12-oxophytodienoate reductase that is required for jasmonate biosynthesis. Mutants are male sterile and defective in pollen dehiscence. Shows activity towards 2,4,6-trinitrotoluene. CFA-Ile, CFA-Leu, CFA-Val, CFA-Met and CFA-Ala can restore the fertility of opr3 plants by inducing filament elongation and anther dehiscence. | protein_coding | OXOPHYTODIENOATE-REDUCTASE 3 (OPR3) | OXOPHYTODIENOATE-REDUCTASE 3 (OPR3); (AtOPR3) |
| AT2G02710 | AT2G02710.1 | Encodes a putative blue light receptor protein. | protein_coding | PAS/LOV PROTEIN B (PLPB) | PAS/LOV PROTEIN B (PLPB);PAS/LOV PROTEIN (PLP) |
| AT1G61870 | AT1G61870.1 | Generic translation factor involved in mitochondrial translation. | protein_coding | PENTATRICOPEPTIDE REPEAT 336 (PPR336) | RIBOSOMAL PPR PROTEIN 1 (RPPR1);PENTATRICOPEPTIDE REPEAT 336 (PPR336) |
| AT2G35130 | AT2G35130.2 | Tetratricopeptide repeat (TPR)-like superfamily protein;(source:Araport11) | protein_coding | PENTATRICOPEPTIDE REPEAT 66 (PPR_66) | PENTATRICOPEPTIDE REPEAT 66 (PPR_66) |
| AT3G59040 | AT3G59040.2 | Involved in chloroplast biogenesis and function. | protein_coding | PENTATRICOPEPTIDE REPEAT PROTEIN 287 (PPR287) | PENTATRICOPEPTIDE REPEAT PROTEIN 287 (PPR287) |
| AT5G48470 | AT5G48470.1 | hypothetical protein;(source:Araport11) | protein_coding | PEP-RELATED DEVELOPMENT ARRESTED 1 (PRDA1) | PEP-RELATED DEVELOPMENT ARRESTED 1 (PRDA1) |

| | | | | | |
|---|---|---|---|---|---|
| AT5G23940 | AT5G23940.1 | Encodes PERMEABLE LEAVES3 (PEL3), a putative acyl-transferase. Mutation in this locus results in altered trichome phenotype (trcichomes become tangled during leaf expansion). Additional phenotype includes altered cuticle layer. | protein_coding | PERMEABLE LEAVES3 (PEL3) | EMBRYO DEFECTIVE 3009 (EMB3009);DEFECTIVE IN CUTICULAR RIDGES (DCR);PERMEABLE LEAVES3 (PEL3) |
| AT4G33420 | AT4G33420.1 | Peroxidase superfamily protein;(source:Araport11) | protein_coding | PEROXIDASE 47 (PRX47) | PEROXIDASE 47 (PRX47) |
| AT2G26350 | AT2G26350.1 | Zinc-binding peroxisomal integral membrane protein (PEX10). Inserted directly from the cytosol into peroxisomes and is involved in importing proteins into the peroxisome. Required for embryogenesis. | protein_coding | PEROXIN 10 (PEX10) | (ATPEX10);PEROXIN 10 (PEX10) |
| AT3G04460 | AT3G04460.1 | RING finger protein involved in peroxisome biogenesis. Also involved in peroxisomal import of nitric oxide synthase. Has been demonstrated to have E3 ubiquitin ligase activity. | protein_coding | PEROXIN-12 (PEX12) | PEROXIN-12 (PEX12);ABERRANT PEROXISOME MORPHOLOGY 4 (APM4);PEROXIN-12 (ATPEX12) |
| AT3G06050 | AT3G06050.1 | Encodes a mitochondrial matrix localized peroxiredoxin involved in redox homeostasis. Knockout mutants have reduced root growth under certain oxidative stress conditions. | protein_coding | PEROXIREDOXIN IIF (PRXIIF) | PEROXIREDOXIN IIF (ATPRXIIF);PEROXIREDOXIN IIF (PRXIIF) |
| AT2G39970 | AT2G39970.1 | Encodes peroxisomal membrane protein 38 (PMP38). Mutation in this protein results in enlargement of peroxisomes. Delivers NAD+ for optimal fatty acid degradation during storage oil mobilization. | protein_coding | PEROXISOMAL NAD CARRIER (PXN) | PEROXISOMAL NAD CARRIER (PXN);ABERRANT PEROXISOME MORPHOLOGY 3 (APEM3);PEROXISOMAL MEMBRANE PROTEIN 38 (PMP38) |
| AT5G14520 | AT5G14520.1 | Encodes a nucleolar protein that plays an essential role in cell growth and survival through its regulation of ribosome biogenesis and mitotic progression. | protein_coding | PESCADILLO (PES) | PESCADILLO (PES) |
| AT5G61770 | AT5G61770.2 | A single-copy gene encoding a 346 aa protein with a single Brix domain. Similar to yeast ribosome biogenesis proteins Ssf1/2. | protein_coding | PETER PAN-LIKE PROTEIN (PPAN) | (SNAIL1);PETER PAN-LIKE PROTEIN (PPAN) |
| AT5G13800 | AT5G13800.2 | Encodes a pheophytinase that is involved in chlorophyll breakdown. Its transcript levels increase during senescence and pph-1 mutants have a stay-green phenotype. | protein_coding | PHEOPHYTINASE (PPH) | PHEOPHYTINASE (PPH);CO-REGULATED WITH NYE1 (CRN1) |
| AT2G38060 | AT2G38060.1 | Encodes an inorganic phosphate transporter (PHT4;2). | protein_coding | PHOSPHATE TRANSPORTER 4;2 (PHT4;2) | PHOSPHATE TRANSPORTER 4;2 (PHT4;2) |
| AT3G46980 | AT3G46980.3 | Encodes an inorganic phosphate transporter (PHT4;3). | protein_coding | PHOSPHATE TRANSPORTER 4;3 (PHT4;3) | PHOSPHATE TRANSPORTER 4;3 (PHT4;3) |
| AT5G20380 | AT5G20380.1 | Encodes an inorganic phosphate transporter (PHT4;5). | protein_coding | PHOSPHATE TRANSPORTER 4;5 (PHT4;5) | PHOSPHATE TRANSPORTER 4;5 (PHT4;5) |
| AT3G52190 | AT3G52190.1 | Encodes a plant specific protein structurally related to the SEC12 proteins of the early secretory pathway. Mutation of PHF1 impairs Pi transport. Expression was detected in all tissues, and was induced by Pi starvation. Localized in endoplasmic reticulum (ER), and mutation of PHF1 resulted in ER retention and reduced accumulation of the plasma membrane PHT1;1 transporter. Its expression is responsive to both phosphate (Pi) and phosphite (Phi) in shoots. | protein_coding | PHOSPHATE TRANSPORTER TRAFFIC FACILITATOR1 (PHF1) | (AtPHF1);PHOSPHATE TRANSPORTER TRAFFIC FACILITATOR1 (PHF1) |
| AT1G12370 | AT1G12370.2 | encodes an amino acid sequence with significant homology to the recently characterized type II photolyases. The uvr2-1 mutant is unable to remove CPDs in vivo, and plant extracts lack detectable photolyase activity , is sensitive to UV-B and is an allele | protein_coding | PHOTOLYASE 1 (PHR1) | UV RESISTANCE 2 (UVR2);PHOTOLYASE 1 (PHR1) |
| AT3G47390 | AT3G47390.1 | Encodes a protein that is believed to function as a pyrimidine reductase involved in riboflavin and FAD biosynthesis. phs1 was identified as a photosensitive mutant that shows reduced growth, chloroplast developmental abnormalities, reduced chlorophyll levels, increased oxidative stress, reduced NADPH/NADP+ ratios, reduced photosystem I electron | protein_coding | PHOTOSENSITIVE 1 (PHS1) | PYRIMIDINE REDUCTASE (PYRR);PHOTOSENSITIVE 1 (PHS1) |

| | | | | | |
|---|---|---|---|---|---|
| | | transport, and reduced photosynthetic protein levels under high light conditions. Many of these abnormal phenotypes likely arise from the reduction in the levels of FAD in the phs1 mutant. | | | |
| AT5G43750 | AT5G43750.1 | NAD(P)H dehydrogenase 18;(source:Araport11) | protein_coding | PHOTOSYNTHETIC NDH SUBCOMPLEX B 5 (PnsB5) | PHOTOSYNTHETIC NDH SUBCOMPLEX B 5 (PnsB5);NAD(P)H DEHYDROGENASE 18 (NDH18) |
| AT1G61520 | AT1G61520.1 | PSI type III chlorophyll a/b-binding protein (Lhca3*1) The mRNA is cell-to-cell mobile. | protein_coding | PHOTOSYSTEM I LIGHT HARVESTING COMPLEX GENE 3 (LHCA3) | PHOTOSYSTEM I LIGHT HARVESTING COMPLEX GENE 3 (LHCA3) |
| AT2G01490 | AT2G01490.1 | Encodes a phytanoyl-CoA 2-hydroxylase (PAHX). The mRNA is cell-to-cell mobile. | protein_coding | PHYTANOYL-COA 2-HYDROXYLASE (PAHX) | PHYTANOYL-COA 2-HYDROXYLASE (PAHX) |
| AT4G14210 | AT4G14210.1 | Encodes phytoene desaturase (phytoene dehydrogenase), an enzyme that catalyzes the desaturation of phytoene to zeta-carotene during carotenoid biosynthesis. Processed protein is localized to the plastid. | protein_coding | PHYTOENE DESATURASE 3 (PDS3) | PHYTOENE DESATURASE (PDS);PIGMENT DEFECTIVE 226 (PDE226);PHYTOENE DESATURASE 3 (PDS3) |
| AT3G48500 | AT3G48500.2 | Nucleic acid-binding, OB-fold-like protein;(source:Araport11) | protein_coding | PIGMENT DEFECTIVE 312 (PDE312) | PIGMENT DEFECTIVE 312 (PDE312); (TAC10);PLASTID TRANSCRIPTIONALLY ACTIVE 10 (PTAC10) |
| AT1G80770 | AT1G80770.1 | P-loop containing nucleoside triphosphate hydrolases superfamily protein;(source:Araport11) | protein_coding | PIGMENT DEFECTIVE 318 (PDE318) | PIGMENT DEFECTIVE 318 (PDE318) |
| AT3G55250 | AT3G55250.1 | Encodes a nucleus-encoded protein, Photosystem I Assembly 3 (PSA3), that is required for PSI accumulation. | protein_coding | PIGMENT DEFECTIVE 329 (PDE329) | PIGMENT DEFECTIVE 329 (PDE329);PHOTOSYSTEM I ASSEMBLY 3 (PSA3) |
| AT2G26510 | AT2G26510.1 | Encodes a plasma-membrane localized nucleobase transporter capable of transporting adenine, guanine, uracil and hypoxanthine. Likely to be a proton-nucleobase symporter. | protein_coding | PIGMENT DEFECTIVE EMBRYO 135 (PDE135) | NUCLEOBASE ASCORBATE TRANSPORTER 3 (NAT3);PIGMENT DEFECTIVE EMBRYO 135 (PDE135) |
| AT2G26700 | AT2G26700.1 | Member of AGC VIIIa Kinase gene family. Encodes PID2, a homolog of PID. Simultaneous disruption of PID(AT2G34650) and its 3 closest homologs (PID2/AT2G26700, WAG1/AT1G53700, and WAG2/AT3G14370) abolishes the formation of cotyledons. | protein_coding | PINOID2 (PID2) | AGC VIIIA KINASE 1-10 (AGC1-10);PINOID2 (PID2) |
| AT3G22590 | AT3G22590.1 | Encodes PLANT HOMOLOGOUS TO PARAFIBROMIN (PHP), a homolog of human Paf1 Complex (Paf1C) subunit Parafibromin. Human Parafibromin assists in mediating output from the Wnt signaling pathway, and dysfunction of the encoding gene HRPT2 conditions specific cancer-related disease phenotypes. PHP resides in a ~670-kDa protein complex in nuclear extracts, and physically interacts with other known Paf1C-related proteins in vivo. Loss of PHP specifically conditioned accelerated phase transition from vegetative growth to flowering and resulted in misregulation of a very limited subset of genes that included the flowering repressor FLOWERING LOCUS C (FLC). Member of PAF-C complex. | protein_coding | PLANT HOMOLOGOUS TO PARAFIBROMIN (PHP) | (CDC73);PLANT HOMOLOGOUS TO PARAFIBROMIN (PHP) |
| AT5G67530 | AT5G67530.1 | plant U-box 49;(source:Araport11) | protein_coding | PLANT U-BOX 49 (PUB49) | PLANT U-BOX 49 (PUB49);PLANT U-BOX 49 (ATPUB49) |
| AT2G34640 | AT2G34640.1 | Present in transcriptionally active plastid chromosomes. Involved in plastid gene expression. | protein_coding | PLASTID TRANSCRIPTIONALLY ACTIVE 12 (PTAC12) | PLASTID TRANSCRIPTIONALLY ACTIVE 12 (PTAC12); (TAC12);HEMERA (HMR) |
| AT1G74850 | AT1G74850.1 | Present in transcriptionally active plastid chromosomes. Involved in plastid gene expression. | protein_coding | PLASTID TRANSCRIPTIONALLY ACTIVE 2 (PTAC2) | PIGMENT DEFECTIVE 343 (PDE343);PLASTID TRANSCRIPTIONALLY ACTIVE 2 (PTAC2) |
| AT2G31320 | AT2G31320.1 | Encodes a poly(ADP-ribose) polymerase. | protein_coding | POLY(ADP-RIBOSE) POLYMERASE 1 (PARP1) | POLY(ADP-RIBOSE) POLYMERASE 2 (PARP2);POLY(ADP-RIBOSE) POLYMERASE 1 (PARP1); (ATPARP2) |
| AT1G04690 | AT1G04690.1 | potassium channel beta subunit 1;(source:Araport11) | protein_coding | POTASSIUM CHANNEL BETA SUBUNIT 1 (KAB1) | POTASSIUM CHANNEL BETA SUBUNIT 1 (KAB1); (KV-BETA1) |
| AT5G51700 | AT5G51700.1 | Encodes a resistance signalling protein with two zinc binding (CHORD) domains that are highly conserved across eukaryotic phyla. Mutant has reduced RPS5 and RPM1 | protein_coding | PPHB SUSCEPTIBLE 2 (PBS2) | PPHB SUSCEPTIBLE 2 (PBS2);REQUIRED FOR MLA12 RESISTANCE 1 (RAR1); (RPR2); (ATRAR1) |

| | | | | | |
|---|---|---|---|---|---|
| | | mediated resistance. Potentially involved in transduction of R gene mediated disease resistance. Required for R protein accumulation. | | | |
| AT5G60960 | AT5G60960.1 | Encodes PNM1 (for PPR protein localized to the nucleus and mitochondria 1), a PPR protein that is dual localized to mitochondria and nuclei. Loss of PNM1 function in mitochondria, but not in nuclei, is lethal for the embryo. In mitochondria, it is associated with polysomes and may play a role in translation. | protein_coding | PPR PROTEIN LOCALIZED TO THE NUCLEUS AND MITOCHONDRIA 1 (PNM1) | RIBOSOMAL PENTATRICOPEPTIDE REPEAT PROTEIN 9 (RPPR9);PPR PROTEIN LOCALIZED TO THE NUCLEUS AND MITOCHONDRIA 1 (PNM1) |
| AT4G02060 | AT4G02060.1 | Member of the minichromosome maintenance complex, involved in DNA replication initiation. Abundant in proliferating and endocycling tissues. Localized in the nucleus during G1, S and G2 phases of the cell cycle, and are released into the cytoplasmic compartment during mitosis. Binds chromatin. | protein_coding | PROLIFERA (PRL) | PROLIFERA (PRL); (MCM7) |
| AT2G42810 | AT2G42810.2 | Encodes a phytochrome-specific type 5 serine/threonine protein phosphatase. It dephosphorylates active Pfr-phytochromes. Controls light signal flux by enhancing phytochrome stability and affinity for a signal transducer. The gene is alternately spliced. This variant is an integral membrane protein localized to the ER and nuclear envelope. Belongs to one of the 36 carboxylate clamp (CC)-tetratricopeptide repeat (TPR) proteins (Prasad 2010, Pubmed ID: 20856808) with potential to interact with Hsp90/Hsp70 as co-chaperones. It also regulates tetrapyrrole biosynthesis through the accumulation of Mg-ProtoIX and acts as a negative regulator of photosynthesis associated nuclear gene expression during chloroplast biogenesis and development. | protein_coding | PROTEIN PHOSPHATASE 5 (PP5) | PROTEIN PHOSPHATASE 5 (PP5);PHYTOCHROME-ASSOCIATED PROTEIN PHOSPHATASE 5 (PAPP5);ARABIDOPSIS THALIANA PROTEIN PHOSPHATASE 5 (AtPP5);PROTEIN PHOSPHATASE 5.2 (PP5.2) |
| AT4G31850 | AT4G31850.1 | encodes a protein containing 27 pentatrico-peptide repeat (PPR) motifs. Functions in the stabilization of petL operon RNA and also in the translation of petL. | protein_coding | PROTON GRADIENT REGULATION 3 (PGR3) | PROTON GRADIENT REGULATION 3 (PGR3) |
| AT3G16810 | AT3G16810.1 | Encodes a member of the Arabidopsis Pumilio (APUM) proteins containing PUF domain (eight repeats of approximately 36 amino acids each). PUF proteins regulate both mRNA stability and translation through sequence-specific binding to the 3' UTR of target mRNA transcripts. | protein_coding | PUMILIO 24 (PUM24) | PUMILIO 24 (APUM24);PUMILIO 24 (PUM24) |
| AT2G24220 | AT2G24220.1 | Member of a family of proteins related to PUP1, a purine transporter. May be involved in the transport of purine and purine derivatives such as cytokinins, across the plasma membrane. | protein_coding | PURINE PERMEASE 5 (PUP5) | PURINE PERMEASE 5 (PUP5);PURINE PERMEASE 5 (ATPUP5) |
| AT5G57140 | AT5G57140.1 | purple acid phosphatase 28;(source:Araport11) | protein_coding | PURPLE ACID PHOSPHATASE 28 (PAP28) | PURPLE ACID PHOSPHATASE 28 (PAP28);PURPLE ACID PHOSPHATASE 28 (ATPAP28) |
| AT2G36570 | AT2G36570.1 | Leucine-rich repeat protein kinase family protein;(source:Araport11) | protein_coding | PXY/TDR-CORRELATED 1 (PXC1) | PXY/TDR-CORRELATED 1 (PXC1) |
| AT5G53580 | AT5G53580.1 | NAD(P)-linked oxidoreductase superfamily protein;(source:Araport11) | protein_coding | PYRIDOXAL REDUCTASE 1 (PLR1) | (AtPLR1);PYRIDOXAL REDUCTASE 1 (PLR1) |
| AT5G60540 | AT5G60540.1 | Encodes a protein predicted to function in tandem with PDX1 to form glutamine amidotransferase complex with involved in vitamin B6 biosynthesis. PDX2 is predicted to function as glutaminase within the complex. | protein_coding | PYRIDOXINE BIOSYNTHESIS 2 (PDX2) | EMBRYO DEFECTIVE 2407 (EMB2407);PYRIDOXINE BIOSYNTHESIS 2 (ATPDX2);PYRIDOXINE BIOSYNTHESIS 2 (PDX2) |
| AT3G17810 | AT3G17810.1 | Encodes a protein predicted to have dihydropyrimidine dehydrogenase activity. Its activity has not been demonstrated in vivo, but, it is required for efficient uracil catabolism in Arabidopsis. It localizes to the plastid. | protein_coding | PYRIMIDINE 1 (PYD1) | PYRIMIDINE 1 (PYD1) |

| | | | | | |
|---|---|---|---|---|---|
| AT1G01090 | AT1G01090.1 | pyruvate dehydrogenase E1 alpha subunit | protein_coding | PYRUVATE DEHYDROGENASE E1 ALPHA (PDH-E1 ALPHA) | PYRUVATE DEHYDROGENASE E1 ALPHA (PDH-E1 ALPHA) |
| AT5G55590 | AT5G55590.1 | Encodes a protein with pectin methylesterase activity. No change in activity were detected in mutants defective in this gene, which was interpreted as a result of redundancy of product function with other pectin methylesterases. The gene product is required for pollen separation during normal development. In qrt mutants, the outer walls of the four meiotic products of the pollen mother cell are fused, and pollen grains are released in tetrads.May be required for cell type-specific pectin degradation. | protein_coding | QUARTET 1 (QRT1) | QUARTET 1 (QRT1) |
| AT4G00740 | AT4G00740.1 | Encodes a Golgi-localized type II membrane pectin methyltransferase regulating cell wall biosynthesis in suspension cells. | protein_coding | QUASIMODO 3 (QUA3) | QUASIMODO 3 (QUA3) |
| AT2G01350 | AT2G01350.1 | At2g01350 encodes quinolinate phosphoribosyl transferase involved in NAD biosynthesis as shown by heterologous expression in E. coli. | protein_coding | QUINOLINATE PHOSHORIBOSYLTRANSFERASE (QPT) | QUINOLINATE PHOSHORIBOSYLTRANSFERASE (QPT) |
| AT5G08710 | AT5G08710.1 | Regulator of chromosome condensation (RCC1) family protein;(source:Araport11) | protein_coding | RCC1/UVR8/GEF-LIKE 1 (RUG1) | RCC1/UVR8/GEF-LIKE 1 (RUG1) |
| AT5G60870 | AT5G60870.1 | Encodes a mitochondrial protein RUG3 that is required for accumulation of mitochondrial respiratory chain complex I. RUG3 is related to human REGULATOR OF CHROMOSOME CONDENSATION 1 (RCC1) and Arabidopsis UV-B RESISTANCE 8 (UVR8). | protein_coding | RCC1/UVR8/GEF-LIKE 3 (RUG3) | RCC1/UVR8/GEF-LIKE 3 (RUG3) |
| AT3G20390 | AT3G20390.1 | Encodes a plastidial RidA (Reactive Intermediate Deaminase A) homolog that hydrolyzes the enamines/imines formed by Thr dehydratase from Ser or Thr. RidA accelerates the deamination of reactive enamine/imine intermediates produced by threonine dehydratase (At3g10050) with threonine or serine as substrates. In the absence of RidA, the serine-derived imine inactivates BCAT3 (At3g49680). RidA thus pre-empts damage to BCAT3 by hydrolyzing the reactive imine before it does damage. | protein_coding | REACTIVE INTERMEDIATE DEAMINASE A (RIDA) | REACTIVE INTERMEDIATE DEAMINASE A (RIDA) |
| AT1G28340 | AT1G28340.1 | receptor like protein 4;(source:Araport11) | protein_coding | RECEPTOR LIKE PROTEIN 4 (RLP4) | RECEPTOR LIKE PROTEIN 4 (RLP4);RECEPTOR LIKE PROTEIN 4 (AtRLP4) |
| AT2G26590 | AT2G26590.1 | regulatory particle non-ATPase 13;(source:Araport11) | protein_coding | REGULATORY PARTICLE NON-ATPASE 13 (RPN13) | REGULATORY PARTICLE NON-ATPASE 13 (RPN13) |
| AT1G77470 | AT1G77470.1 | Encodes a protein with high homology to the Replication Factor C, Subunit 3 (RFC3) of yeast and other eukaryotes. rfc3 mutants are hypersensitive to salicylic acid and exhibit enhanced induction of PR genes and resistance against virulent oomycete Hyaloperonospora arabidopsidis Noco2. The enhanced pathogen resistance in the mutant is NPR1-independent. | protein_coding | REPLICATION FACTOR C SUBUNIT 3 (RFC3) | REPLICATION FACTOR C SUBUNIT 3 (RFC3);EMBRYO DEFECTIVE 2810 (EMB2810);REPLICATION FACTOR C 5 (RFC5) |
| AT2G44420 | AT2G44420.1 | protein N-terminal asparagine amidohydrolase family protein;(source:Araport11) | protein_coding | RESIDUE-SPECIFIC N-TERMINAL AMIDASE 1 (NTAN1) | RESIDUE-SPECIFIC N-TERMINAL AMIDASE 1 (NTAN1) |
| AT1G69380 | AT1G69380.1 | Encodes a mitochondria-localized protein that is required for cell division in the root meristem. | protein_coding | RETARDED ROOT GROWTH (RRG) | RETARDED ROOT GROWTH (RRG) |
| AT4G09730 | AT4G09730.1 | Encodes RH39, a DEAD-box protein involved in the introduction of the hidden break into the 23S rRNA in the chloroplasts. Recombinant RH39 binds to the 23S rRNA in a segment adjacent to the stem-loop creating the hidden break target loop in a sequence-dependent manner. Has ATP-hydrolyzing activity at a Kcat of 5.3 /min in the presence of rRNA sequence. Mutants have drastically reduced level of | protein_coding | RH39 (RH39) | RH39 (RH39) |

| | | | | | |
|---|---|---|---|---|---|
| | | level of ribulose 1,5-bisphosphate carboxylase/oxygenase. The mRNA is cell-to-cell mobile. | | | |
| AT3G59520 | AT3G59520.1 | RHOMBOID-like protein 13;(source:Araport11) | protein_coding | RHOMBOID-LIKE PROTEIN 13 (RBL13) | ARABIDOPSIS RHOMBOID-LIKE PROTEIN 12 (ATRBL12);RHOMBOID-LIKE PROTEIN 13 (RBL13);RHOMBOID-LIKE PROTEIN 13 (ATRBL13);RHOMBOID-LIKE PROTEIN 12 (RBL12) |
| AT3G58460 | AT3G58460.2 | RHOMBOID-like protein 15;(source:Araport11) | protein_coding | RHOMBOID-LIKE PROTEIN 15 (RBL15) | ARABIDOPSIS RHOMBOID-LIKE PROTEIN 11 (ATRBL11);RHOMBOID-LIKE PROTEIN 11 (RBL11);RHOMBOID-LIKE PROTEIN 15 (RBL15);RHOMBOID-LIKE PROTEIN 15 (ATRBL15) |
| AT1G17160 | AT1G17160.1 | RBSK is a plastid localized ribokinase involved in nucleoside metabolism. It is the only member of this gene family in Arabidopsis. | protein_coding | RIBOKINASE (RBSK) | RIBOKINASE (RBSK) |
| AT1G60770 | AT1G60770.1 | Ribosomal pentatricopeptide repeat protein | protein_coding | RIBOSOMAL PENTATRICOPEPTIDE REPEAT PROTEIN 4 (RPPR4) | RIBOSOMAL PENTATRICOPEPTIDE REPEAT PROTEIN 4 (RPPR4) |
| AT2G37230 | AT2G37230.1 | Ribosomal pentatricopeptide repeat protein | protein_coding | RIBOSOMAL PENTATRICOPEPTIDE REPEAT PROTEIN 5 (RPPR5) | RIBOSOMAL PENTATRICOPEPTIDE REPEAT PROTEIN 5 (RPPR5) |
| AT5G30510 | AT5G30510.1 | ribosomal protein S1;(source:Araport11) | protein_coding | RIBOSOMAL PROTEIN S1 (RPS1) | PLASTID RIBOSOMAL PROTEIN S1 (PRPS1);RIBOSOMAL PROTEIN S1 (RPS1); (ARRPS1) |
| AT4G15850 | AT4G15850.1 | plant DEAD box-like RNA helicase. | protein_coding | RNA HELICASE 1 (RH1) | RNA HELICASE 1 (RH1);RNA HELICASE 1 (ATRH1) |
| AT1G16280 | AT1G16280.1 | Encodes a putative DEAD-box RNA helicase. Essential for female gametogenesis. | protein_coding | RNA HELICASE 36 (RH36) | RNA HELICASE 36 (RH36);SLOW WALKER 3 (SWA3);ARABIDOPSIS THALIANA RNA HELICASE 36 (AtRH36) |
| AT3G20420 | AT3G20420.1 | double-stranded RNA binding / ribonuclease III. Required for 3' external transcribed spacer (ETS) cleavage of the pre-rRNA in vivo. Localizes in the nucleus and cytoplasm. | protein_coding | RNASE THREE-LIKE PROTEIN 2 (RTL2) | RNASEIII-LIKE 2 (ATRTL2);RNASE THREE-LIKE PROTEIN 2 (RTL2) |
| AT1G13770 | AT1G13770.1 | root UVB sensitive protein (Protein of unknown function, DUF647);(source:Araport11) | protein_coding | ROOT UV-B SENSITIVE 3 (RUS3) | ROOT UV-B SENSITIVE 3 (RUS3) |
| AT5G01510 | AT5G01510.1 | root UVB sensitive protein (Protein of unknown function, DUF647);(source:Araport11) | protein_coding | ROOT UV-B SENSITIVE 5 (RUS5) | ROOT UV-B SENSITIVE 5 (RUS5) |
| AT2G41530 | AT2G41530.1 | Encodes a protein with S-formylglutathione hydrolase activity. | protein_coding | S-FORMYLGLUTATHIONE HYDROLASE (SFGH) | S-FORMYLGLUTATHIONE HYDROLASE (SFGH);ARABIDOPSIS THALIANA S-FORMYLGLUTATHIONE HYDROLASE (AtSFGH) |
| AT3G51830 | AT3G51830.1 | putative transmembrane protein G5p (AtG5) mRNA, complete. autophagy-related (ATG) gene | protein_coding | SAC DOMAIN-CONTAINING PROTEIN 8 (SAC8) | SAC DOMAIN-CONTAINING PROTEIN 8 (SAC8); (ATG5) |
| AT5G37850 | AT5G37850.1 | Encodes a pyridoxal kinase required for root hair development. Mutants are hypersensitive to Na+, K+ and Li+. | protein_coding | SALT OVERLY SENSITIVE 4 (SOS4) | SALT OVERLY SENSITIVE 4 (ATSOS4);SALT OVERLY SENSITIVE 4 (SOS4) |
| AT3G07700 | AT3G07700.3 | ABC1K7 is a member of an atypical protein kinase family that is induced by salt stress. Loss of function mutations affect the metabolic profile of chloroplast lipids. It appears to function along with ABC1K8 in mediating lipid membrane changes in response to stress. | protein_coding | SALT-INDUCED ABC1 KINASE 1 (SIA1) | SALT-INDUCED ABC1 KINASE 1 (SIA1); (ABC1K7); (ATSIA1) |
| AT5G52810 | AT5G52810.1 | SAR-DEFICIENT4 (SARD4) alias ORNITHINE CYCLODEAMINASE/m-CRYSTALLIN (ORNCD1) is involved in the biosynthesis of pipecolic acid. The reductase converts dehydropipecolic acid intermediates generated from L-Lysine by AGD2-LIKE DEFENSE RESPONSE PROTEIN1 (ALD1) to pipecolic acid (PMID:28330936). | protein_coding | SAR DEFICIENT 4 (SARD4) | SAR DEFICIENT 4 (SARD4) |
| AT5G13030 | AT5G13030.1 | Chloroplast localized homolog of SELO. Loss of function mutants have reduced production of reactive oxygen species (ROS) and higher ROS scavenging. | protein_coding | SELENOPROTEIN O (SELO) | SELENOPROTEIN O (SELO) |
| AT3G06510 | AT3G06510.2 | Encodes a protein with beta-glucosidase and galactosyltransferase activity, mutants show increased | protein_coding | SENSITIVE TO FREEZING 2 (SFR2) | SENSITIVE TO FREEZING 2 (SFR2) |

| | | | | | |
|---|---|---|---|---|---|
| | | sensitivity to freezing. Though it is classified as a family I glycosyl hydrolase, it has no hydrolase activity in vitro. | | | |
| AT4G30810 | AT4G30810.1 | serine carboxypeptidase-like 29;(source:Araport11) | protein_coding | SERINE CARBOXYPEPTIDASE-LIKE 29 (scpl29) | SERINE CARBOXYPEPTIDASE-LIKE 29 (scpl29) |
| AT5G37055 | AT5G37055.1 | Encodes SERRATED LEAVES AND EARLY FLOWERING (SEF), an Arabidopsis homolog of the yeast SWC6 protein, a conserved subunit of the SWR1/SRCAP complex. SEF loss-of-function mutants have a pleiotropic phenotype characterized by serrated leaves, frequent absence of inflorescence internodes, bushy aspect, and flowers with altered number and size of organs. sef plants flower earlier than wild-type plants both under inductive and non-inductive photoperiods. SEF, ARP6 and PIE1 might form a molecular complex in Arabidopsis related to the SWR1/SRCAP complex identified in other eukaryotes. | protein_coding | SERRATED LEAVES AND EARLY FLOWERING (SEF) | (AtSWC6);SERRATED LEAVES AND EARLY FLOWERING (SEF) |
| AT2G17900 | AT2G17900.1 | Homology Subgroup S-ET - Protein containing an interrupted SET domain. | protein_coding | SET DOMAIN GROUP 37 (SDG37) | ASH1-RELATED 1 (ASHR1);SET DOMAIN GROUP 37 (SDG37) |
| AT1G07010 | AT1G07010.3 | Calcineurin-like metallo-phosphoesterase superfamily protein;(source:Araport11) | protein_coding | SHEWENELLA-LIKE PROTEIN PHOSPHATASE 1 (SLP1) | SHEWENELLA-LIKE PROTEIN PHOSPHATASE 1 (SLP1); (ATSLP1) |
| AT1G31480 | AT1G31480.1 | encodes a novel protein that may be part of a gene family represented by bovine phosphatidic acid-preferring phospholipase A1 (PA-PLA1)containing a putative transmembrane domain. SGR2 is involved in the formation and function of the vacuole. | protein_coding | SHOOT GRAVITROPISM 2 (SGR2) | SHOOT GRAVITROPISM 2 (SGR2) |
| AT3G48820 | AT3G48820.1 | Encodes a homolog of the animal sialyltransferases but sialyltransferase activity was not detected (Plant Biology 2009, 11:284). Located in the Golgi apparatus. | protein_coding | SIALYLTRANSFERASE-LIKE 2 (SIA2) | SIALYLTRANSFERASE-LIKE 2 (SIA2) |
| AT5G24120 | AT5G24120.1 | Encodes a specialized sigma factor that functions in regulation of plastid genes and is responsible for the light-dependent transcription at the psbD LRP. Activation of SIG5 is dependent upon blue light and mediated by cryptochromes. | protein_coding | SIGMA FACTOR E (SIGE) | SIGMA FACTOR 5 (ATSIG5) |
| AT1G73990 | AT1G73990.1 | Encodes a putative protease SppA (SppA). | protein_coding | SIGNAL PEPTIDE PEPTIDASE (SPPA) | (SPPA1);SIGNAL PEPTIDE PEPTIDASE (SPPA) |
| AT3G61350 | AT3G61350.1 | Encodes an SKP1 interacting partner (SKIP4). | protein_coding | SKP1 INTERACTING PARTNER 4 (SKIP4) | SKP1 INTERACTING PARTNER 4 (SKIP4) |
| AT3G56950 | AT3G56950.2 | One of the Major Intrinsic Proteins(MIPs) which facilitate the passive transport of small molecules across membranes.Belongs to a family of plant aquaporins.Similar to yeast and radish aquaporins. Located on ER. Probably involved in the alleviation of ER stress; the lack of SIP2;1 reduces both pollen germination and pollen tube elongation. | protein_coding | SMALL AND BASIC INTRINSIC PROTEIN 2;1 (SIP2;1) | SMALL AND BASIC INTRINSIC PROTEIN 2;1 (SIP2;1) |
| AT5G06680 | AT5G06680.1 | Encodes protein similar to yeast SCP98. Yeast SCP98 is essential for the microtubule nucleation activity of the gamma-tubulin ring complexes. Enriched at the post-cytokinetic cell edges in leaves and roots. The mRNA is cell-to-cell mobile. | protein_coding | SPINDLE POLE BODY COMPONENT 98 (SPC98) | SPINDLE POLE BODY COMPONENT 98 (SPC98);GAMMA TUBULIN COMPLEX PROTEIN 3 (GCP3);SPINDLE POLE BODY COMPONENT 98 (ATSPC98);ARABIDOPSIS THALIANA GAMMA TUBULIN COMPLEX PROTEIN 3 (ATGCP3) |
| AT2G47580 | AT2G47580.1 | encodes spliceosomal protein U1A | protein_coding | SPLICEOSOMAL PROTEIN U1A (U1A) | SPLICEOSOMAL PROTEIN U1A (U1A) |
| AT2G17975 | AT2G17975.1 | SRP1 is a C2C2 type zinc finger protein that binds RNA. It has a role in response to ABA.. It can bind the 3'UTR of ABI2 and appears to be involved in RNA turnover. | protein_coding | STRESS ASSOCIATED RNA-BINDING PROTEIN 1 (SRP1) | STRESS ASSOCIATED RNA-BINDING PROTEIN 1 (SRP1) |
| AT1G31970 | AT1G31970.1 | DEA(D/H)-box RNA helicase family protein;(source:Araport11) | protein_coding | STRESS RESPONSE SUPPRESSOR 1 (STRS1) | (RH5);STRESS RESPONSE SUPPRESSOR 1 (STRS1) |
| AT1G68830 | AT1G68830.1 | STN7 protein kinase; required for state transitions, phosphorylation of the major antenna complex (LHCII) | protein_coding | STT7 HOMOLOG STN7 (STN7) | STT7 HOMOLOG STN7 (STN7) |

| | | | | | |
|---|---|---|---|---|---|
| | | between PSII and PSI, and light adaptation. STN7 is involved in state transitions. | | | |
| AT2G02860 | AT2G02860.1 | encodes a sucrose transporter in sieve elements and a number of sink tissues and cell types. Gene expression is induced by wounding. | protein_coding | SUCROSE TRANSPORTER 2 (SUT2) | SUCROSE TRANSPORTER 2 (SUT2);ARABIDOPSIS THALIANA SUCROSE TRANSPORTER 3 (ATSUC3);SUCROSE TRANSPORTER 3 (SUC3); (ATSUT2) |
| AT3G01910 | AT3G01910.1 | Encodes a homodimeric Mo-enzyme with molybdopterin as organic component of the molybdenum cofactor. It lacks the heme domain that other eukaryotic Mo-enzymes possess and has no redox-active centers other than the molybdenum. SO protein has been found in all parts of the plant. The plant SO combines its enzymatic sulfite oxidation with a subsequent nonenzymatic step using its reaction product H2O2 as intermediate for oxidizing another molecule of sulfite. | protein_coding | SULFITE OXIDASE (SOX) | (AtSO);SULFITE OXIDASE (SOX); (AT-SO) |
| AT3G59770 | AT3G59770.3 | Encodes a phosphoinositide phosphatase. The sac9 null mutant accumulates elevated levels of PtdIns(4,5)P2 and Ins(1,4,5)P3. The mutant plants have characteristics of constitutive stress responses. | protein_coding | SUPPRESSOR OF ACTIN 9 (SAC9) | ARABIDOPSIS THALIANA SUPPRESSOR OF ACTIN 9 (AtSAC9);SUPPRESSOR OF ACTIN 9 (SAC9) |
| AT2G31880 | AT2G31880.1 | Encodes a putative leucine rich repeat transmembrane protein that is expressed in response to Pseudomonas syringae. Expression of SRRLK may be required for silencing via lsiRNAs. Regulates cell death and innate immunity. | protein_coding | SUPPRESSOR OF BIR1 1 (SOBIR1) | EVERSHED (EVR);SUPPRESSOR OF BIR1 1 (SOBIR1) |
| AT1G02100 | AT1G02100.3 | Leucine carboxyl methyltransferase;(source:Araport11) | protein_coding | SUPPRESSOR OF BRI1 (SBI1) | LEUCINE CARBOXYL METHYL TRANSFERASE 1 (LCMT1);SUPPRESSOR OF BRI1 (SBI1) |
| AT5G46580 | AT5G46580.1 | pentatricopeptide (PPR) repeat-containing protein;(source:Araport11) | protein_coding | SUPPRESSOR OF THYLAKOID FORMATION 1 (SOT1) | SUPPRESSOR OF THYLAKOID FORMATION 1 (SOT1) |
| AT5G13650 | AT5G13650.2 | Encodes SVR3, a putative chloroplast TypA translation elongation GTPase. Loss of SVR3 suppresses variegation mediated by var2. SVR3 is essential for plants? ability to develop functional chloroplasts under chilling stress (8C), but not at normal temperature (22C). | protein_coding | SUPPRESSOR OF VARIEGATION 3 (SVR3) | SUPPRESSOR OF VARIEGATION 3 (SVR3) |
| AT4G16390 | AT4G16390.1 | Encodes a pentatricopeptide repeat protein, SVR7 (SUPPRESSOR OF VARIEGATION7), required for FtsH-mediated chloroplast biogenesis. It is involved in accumulation and translation of chloroplast ATP synthase subunits. | protein_coding | SUPPRESSOR OF VARIEGATION 7 (SVR7) | SUPPRESSOR OF VARIEGATION 7 (SVR7) |
| AT1G73720 | AT1G73720.1 | Encodes SMU1, a protein involved in RNA splicing. | protein_coding | SUPPRESSORS OF MEC-8 AND UNC-52 1 (SMU1) | SUPPRESSORS OF MEC-8 AND UNC-52 1 (SMU1) |
| AT2G47210 | AT2G47210.1 | myb-like transcription factor family protein;(source:Araport11) | protein_coding | SWR1 COMPLEX 4 (SWC4) | SWR1 COMPLEX 4 (SWC4); (ATSWC4) |
| AT4G34270 | AT4G34270.1 | TOR signaling pathway protein. | protein_coding | TAP42 INTERACTING PROTEIN OF 41 KDA (TIP41) | TAP42 INTERACTING PROTEIN OF 41 KDA (TIP41) |
| AT5G25150 | AT5G25150.1 | Encodes a putative TATA-binding-protein associated factor TAF5. TAFs are subunits of the general transcription factor IID (TFIID). | protein_coding | TBP-ASSOCIATED FACTOR 5 (TAF5) | TBP-ASSOCIATED FACTOR 5 (TAF5) |
| AT1G04130 | AT1G04130.1 | Encodes one of the 36 carboxylate clamp (CC)-tetratricopeptide repeat (TPR) proteins (Prasad 2010, Pubmed ID: 20856808). Interacts with Hsp90/Hsp70 as co-chaperone. | protein_coding | TETRATRICOPEPTIDE REPEAT 2 (TPR2) | TETRATRICOPEPTIDE REPEAT 2 (TPR2); (AtTPR2) |
| AT4G27800 | AT4G27800.1 | Choroplast protein phosphatase TAP38/PPH1 is required for efficient dephosphorylation of the LHCII anthena and state transition from state 2 to state 1. | protein_coding | THYLAKOID-ASSOCIATED PHOSPHATASE 38 (TAP38) | THYLAKOID-ASSOCIATED PHOSPHATASE 38 (TAP38);PROTEIN PHOSPHATASE 1 (PPH1) |
| AT2G24820 | AT2G24820.1 | translocon at the inner envelope membrane of chloroplasts 55-II;(source:Araport11) | protein_coding | TRANSLOCON AT THE INNER ENVELOPE MEMBRANE OF CHLOROPLASTS 55-II (TIC55-II) | TRANSLOCON AT THE INNER ENVELOPE MEMBRANE OF CHLOROPLASTS 55-II (TIC55-II);TRANSLOCON AT THE INNER ENVELOPE MEMBRANE OF |

| | | | | | |
|---|---|---|---|---|---|
| | | | | | CHLOROPLASTS 55 (AtTic55);TRANSLOCON AT THE INNER ENVELOPE MEMBRANE OF CHLOROPLASTS 55 (Tic55) |
| AT3G21300 | AT3G21300.1 | RNA methyltransferase family protein;(source:Araport11) | protein_coding | TRNA METHYLTRANSFERASE 2A (TRM2A) | (ATTRM2A);TRNA METHYLTRANSFERASE 2A (TRM2A) |
| AT4G27340 | AT4G27340.1 | Met-10+ like family protein;(source:Araport11) | protein_coding | TRNA METHYLTRANSFERASE 5B (TRM5B) | (ATTRM5B);TRNA METHYLTRANSFERASE 5B (TRM5B) |
| AT5G14600 | AT5G14600.1 | S-adenosyl-L-methionine-dependent methyltransferases superfamily protein;(source:Araport11) | protein_coding | TRNA METHYLTRANSFERASE 61 (TRM61) | (ATTRM61);TRNA METHYLTRANSFERASE 61 (TRM61) |
| AT5G24840 | AT5G24840.1 | tRNA (guanine-N-7) methyltransferase;(source:Araport11) | protein_coding | TRNA METHYLTRANSFERASE 8A (TRM8A) | (ATTRM8A);TRNA METHYLTRANSFERASE 8A (TRM8A) |
| AT3G26410 | AT3G26410.1 | Encodes a protein involved in modification of nucleosides in tRNA. Mutants have only 7.3% 2-methylguanosine levels of wild type counterparts. | protein_coding | TRNA MODIFICATION 11 (TRM11) | (AtTRM11);TRNA MODIFICATION 11 (TRM11) |
| AT1G03110 | AT1G03110.1 | Encodes a gene involved in the modification of nucleosides in tRNA. Mutants have no 7-methylguanosine. | protein_coding | TRNA MODIFICATION 82 (TRM82) | (AtTRM82);TRNA MODIFICATION 82 (TRM82) |
| AT2G27760 | AT2G27760.1 | Encodes tRNA isopentenyltransferase, similar to yeast MOD5. | protein_coding | TRNAISOPENTENYLTRANSFERASE 2 (IPT2) | TRNAISOPENTENYLTRANSFERASE 2 (ATIPT2);TRNAISOPENTENYLTRANSFERASE 2 (IPT2); (IPPT) |
| AT5G38530 | AT5G38530.1 | TSBtype2 encodes a type 2 tryptophan synthase beta subunit that catalyzes a condensation reaction between serine and indole to generate tryptophan.It appears to form a homodimer. Its biological role has not yet been determined, but it has a very high affinity for indole which may be involved in allowing TSBtype2 to carefully limit free indole build-up. But, to date no overall change in plant morphology or seedling root growth have been observed in tsbtype2 mutants, indicating that this gene is not essential under optimum conditions. n most organs, TSBtype2 is transcripts are expressed at a lower level than TSB1 but in dry seeds they are expressed at comparable levels. | protein_coding | TRYPTOPHAN SYNTHASE BETA TYPE 2 (TSBtype2) | TRYPTOPHAN SYNTHASE BETA TYPE 2 (TSBtype2) |
| AT4G03560 | AT4G03560.1 | Encodes a depolarization-activated Ca(2+) channel. Anti-sense experiments with this gene as well as Sucrose-H(+) symporters and complementation of yeast sucrose uptake mutant cch1 suggest that this protein mediates a voltage-activated Ca(2+ )influx. Mutants lack detectable SV channel activity suggesting TPC1 is essential component of the SV channel. Patch clamp analysis of loss of function mutation indicates TPC1 does not affect Ca2+ signaling in response to abiotic and biotic stress. | protein_coding | TWO-PORE CHANNEL 1 (TPC1) | TWO-PORE CHANNEL 1 (ATTPC1); (TPC1);TWO-PORE CHANNEL 1 (TPC1);FATTY ACID OXYGENATION UPREGULATED 2 (FOU2);CALCIUM CHANNEL 1 (ATCCH1) |
| AT1G09760 | AT1G09760.1 | U2 small nuclear ribonucleoprotein A;(source:Araport11) | protein_coding | U2 SMALL NUCLEAR RIBONUCLEOPROTEIN A (U2A') | U2 SMALL NUCLEAR RIBONUCLEOPROTEIN A (U2A') |
| AT3G17205 | AT3G17205.1 | ubiquitin protein ligase 6;(source:Araport11) | protein_coding | UBIQUITIN PROTEIN LIGASE 6 (UPL6) | UBIQUITIN PROTEIN LIGASE 6 (UPL6) |
| AT4G31600 | AT4G31600.1 | Encodes a Golgi-localized UDP?glucose/UDP?galactose transporter that affects lateral root emergence. | protein_coding | UDP-GALACTOSE TRANSPORTER 7 (UTR7) | UDP-GALACTOSE TRANSPORTER 7 (UTR7) |
| AT5G41150 | AT5G41150.1 | Confers resistance to UV radiation. Homolog of the human xeroderma pigmentosum group F DNA repair and yeast Rad1 proteins | protein_coding | ULTRAVIOLET HYPERSENSITIVE 1 (UVH1) | ULTRAVIOLET HYPERSENSITIVE 1 (UVH1); (RAD1); (ATRAD1) |
| AT1G14140 | AT1G14140.1 | Mitochondrial substrate carrier family protein;(source:Araport11) | protein_coding | UNCOUPLING PROTEIN 3 (UCP3) | UNCOUPLING PROTEIN 3 (UCP3) |
| AT4G02030 | AT4G02030.2 | Vps51/Vps67 family (components of vesicular transport) protein;(source:Araport11) | protein_coding | UNHINGED (UNH) | UNHINGED (UNH);VACUOLAR PROTEIN SORTING 51 (VPS51) |
| AT3G18630 | AT3G18630.1 | Encodes a uracil-DNA glycosylase (UDG) involved in a base excision DNA repair pathway in mitochondria. | protein_coding | URACIL DNA GLYCOSYLASE (UNG) | URACIL DNA GLYCOSYLASE (UNG);URACIL DNA GLYCOSYLASE (ATUNG) |

| | | | | | |
|---|---|---|---|---|---|
| AT2G26230 | AT2G26230.1 | Encodes a urate oxidase that is involved in peroxisome maintenance. | protein_coding | URATE OXIDASE (UOX) | URATE OXIDASE (UOX) |
| AT2G34470 | AT2G34470.2 | Encodes a urease accessory protein which is essential for the activation of plant urease. | protein_coding | UREASE ACCESSORY PROTEIN G (UREG) | (PSKF109);UREASE ACCESSORY PROTEIN G (UREG) |
| AT5G43600 | AT5G43600.1 | Encodes a protein with ureidoglycolate amidohydrolase activity in vitro. It is 27% identical and 43% similar to the E. coli allantoate amidohydrolase (AAH), but, in vitro assays with purified protein and allantoate as a substrate do not show any increase in ammonium concentration, indicating that there this enzyme has no AAH activity. The mRNA is cell-to-cell mobile. | protein_coding | UREIDOGLYCOLATE AMIDOHYDROLASE (UAH) | ARABIDOPSIS THALIANA ALLANTOATE AMIDOHYDROLASE 2 (ATAAH-2);UREIDOGLYCOLATE AMIDOHYDROLASE (UAH) |
| AT1G05620 | AT1G05620.1 | Encodes a cytosolic inosine nucleoside hydrolase. It forms a heterocomplex with NSH1 with almost two orders of magnitude higher catalytic efficiency for xanthosine hydrolysis than observed for NSH1 alone. Transcript levels for this gene are elevated in older leaves suggesting that it may play a role in purine catabolism during senescence. | protein_coding | URIDINE-RIBOHYDROLASE 2 (URH2) | NUCLEOSIDE HYDROLASE 2 (NSH2);URIDINE-RIBOHYDROLASE 2 (URH2) |
| AT2G36310 | AT2G36310.1 | Encodes a cytoplasmic nucleoside hydrolase. It has the highest levels of activity with uridine followed by xanthosine. It shows little activity with inosine and none with cytidine. Mutant analyses indicate that it plays a role in purine and pyrimidine catabolism. | protein_coding | URIDINE-RIBOHYDROLASE 1 (URH1) | NUCLEOSIDE HYDROLASE 1 (NSH1);URIDINE-RIBOHYDROLASE 1 (URH1) |
| AT1G09380 | AT1G09380.1 | nodulin MtN21-like transporter family protein | protein_coding | USUALLY MULTIPLE ACIDS MOVE IN AND OUT TRANSPORTERS 25 (UMAMIT25) | USUALLY MULTIPLE ACIDS MOVE IN AND OUT TRANSPORTERS 25 (UMAMIT25) |
| AT2G05170 | AT2G05170.1 | Homologous to yeast VPS11. Forms a complex with VCL1 and AtVPS33. Involved in vacuolar biogenesis. The mRNA is cell-to-cell mobile. | protein_coding | VACUOLAR PROTEIN SORTING 11 (VPS11) | VACUOLAR PROTEIN SORTING 11 (VPS11);VACUOLAR PROTEIN SORTING 11 (ATVPS11) |
| AT1G12470 | AT1G12470.1 | zinc ion binding protein;(source:Araport11) | protein_coding | VACUOLAR PROTEIN SORTING 18 (VPS18) | VACUOLAR PROTEIN SORTING 18 (VPS18) |
| AT3G61770 | AT3G61770.1 | Acid phosphatase/vanadium-dependent haloperoxidase-related protein;(source:Araport11) | protein_coding | VACUOLAR PROTEIN SORTING 30 (VPS30) | VACUOLAR PROTEIN SORTING 30 (VPS30) |
| AT2G38020 | AT2G38020.1 | necessary for proper vacuole formation and morphogenesis in Arabidopsis | protein_coding | VACUOLELESS 1 (VCL1) | VACUOLELESS 1 (VCL1); (EMB258);MANGLED (MAN) |
| AT4G29830 | AT4G29830.1 | VIP3 protein is composed of repeats of WD motif which is involved in protein complex formation. The gene is involved in flower timing and flower development. This gene is predicted to encode a protein with a DWD motif. It can bind to DDB1a in Y2H assays, and DDB1b in co-IP assays, and may be involved in the formation of a CUL4-based E3 ubiquitin ligase. Loss of gene function leads to a redistribution of H3K4me3 and K3K36me2 modifications within genes but not a change in the overall abundance of these modifications within chromatin. Also known as SKI8, a component of the SKI complex involved in exosome mediated RNA degredation. Member of PAF-C complex. | protein_coding | VERNALIZATION INDEPENDENCE 3 (VIP3) | A. THALIANA HOMOLOG OF YEAST SKI8 (SKI8);VERNALIZATION INDEPENDENCE 3 (VIP3) |
| AT1G56180 | AT1G56180.1 | VIR3 encodes a putative chloroplast metalloprotease that is localized to the thylakoid membrane. The vir3-1 mutant came out of a genetic suppressor screen of the Arabidopsis variegation mutant yellow variegated (var2). The suppressor displayed an additional virescent phenotype, i.e. the bases of young leaves were yellow, and leaf color gradually turned to green toward the leaf tips. | protein_coding | VIRESCENT3-1 (VIR3) | (VIR3);VIRESCENT3-1 (VIR3) |
| AT5G20520 | AT5G20520.1 | Encodes a Bem46-like protein. WAV2 negatively regulates root bending when roots alter their growth direction. It's not | protein_coding | WAVY GROWTH 2 (WAV2) | WAVY GROWTH 2 (WAV2) |

| | | | | | |
|---|---|---|---|---|---|
| | | involved in sensing environmental stimuli (e.g. gravity, light, water, touch). | | | |
| AT1G09850 | AT1G09850.1 | Arabidopsis thaliana papain-like cysteine peptidase | protein_coding | XYLEM BARK CYSTEINE PEPTIDASE 3 (XBCP3) | XYLEM BARK CYSTEINE PEPTIDASE 3 (XBCP3) |
| AT1G11545 | AT1G11545.1 | xyloglucan endotransglucosylase/hydrolase 8;(source:Araport11) | protein_coding | XYLOGLUCAN ENDOTRANSGLUCOSYLASE/HYDROLASE 8 (XTH8) | XYLOGLUCAN ENDOTRANSGLUCOSYLASE/HYDROLASE 8 (XTH8) |
| AT2G21370 | AT2G21370.1 | Although this gene has a sequence similar to xylulose kinases, several lines of experimental evidence suggest that it does not act on xylulose or deoxy-xylulose. | protein_coding | XYLULOSE KINASE-1 (XK-1) | XYLULOSE KINASE-1 (XK-1);XYLULOSE KINASE 1 (XK1) |
| AT3G04870 | AT3G04870.1 | Involved in the biosynthesis of carotenes and xanthophylls, reduces zeta-carotene to lycopene. | protein_coding | ZETA-CAROTENE DESATURASE (ZDS) | SPONTANEOUS CELL DEATH 1 (SPC1);ZETA-CAROTENE DESATURASE (ZDS);PIGMENT DEFECTIVE EMBRYO 181 (PDE181) |
