## Supplementary material for "Phylogenomics unravels speciation patterns in temperate-montane plant species: a case study on the recently radiating *Ranunculus auricomus* species complex": Table S3

**Table S3** Number of reads per sample, along with information about quality trimming and duplicate removal.

| **Sample no.** | **Taxon** | **Nr. of pairs** | **Nr. of reads** | **Both surviving** | **Forward only surviving** | **Reverse only surviving** | **Nr. reads after quality trimming** | **% quality trimmed reads** | **Nr. reads without duplicates** | **% duplicates** |
|---|---|---|---|---|---|---|---|---|---|---|
| DU003-1 | Ranunculus austroslovenicus | 434604 | 869208 | 418577 | 9204 | 4216 | 850574 | 2.15 | 788804 | 7.27 |
| LH012-8 | Ranunculus austroslovenicus | 614059 | 1228118 | 593017 | 11436 | 6398 | 1203868 | 1.98 | 1088796 | 9.56 |
| 2 | Ranunculus calapius | 591569 | 1183138 | 547175 | 33091 | 4235 | 1131676 | 4.35 | 1122814 | 0.79 |
| Du35351-15 | Ranunculus calapius | 878989 | 1757978 | 843735 | 20017 | 7997 | 1715484 | 2.42 | 1630888 | 4.94 |
| 9126-2 | Ranunculus carpaticola | 878231 | 1756462 | 836843 | 19697 | 9823 | 1703206 | 3.04 | 1545974 | 9.24 |
| DU013-1 | Ranunculus carpaticola | 791056 | 1582112 | 742839 | 36202 | 5318 | 1527198 | 3.48 | 1370336 | 10.28 |
| LH40-4 | Ranunculus carpaticola | 590893 | 1181786 | 552794 | 28544 | 4068 | 1138200 | 3.69 | 1070354 | 5.97 |
| LH006-17 | Ranunculus cassubicifolius | 715201 | 1430402 | 674038 | 29304 | 5430 | 1382810 | 3.33 | 1284166 | 7.14 |
| LH016-14 | Ranunculus cassubicifolius | 1089476 | 2178952 | 1051712 | 20883 | 10570 | 2134877 | 2.03 | 1928887 | 9.65 |
| LH008-7 | Ranunculus cassubicifolius | 742197 | 1484394 | 712605 | 17458 | 7015 | 1449683 | 2.34 | 1318205 | 9.07 |
| LH009-3 | Ranunculus cassubicifolius | 677510 | 1355020 | 589344 | 77486 | 2341 | 1258515 | 7.13 | 1193781 | 5.15 |
| Du33354-2 | Ranunculus cebennensis | 734001 | 1468002 | 694081 | 26409 | 6366 | 1420937 | 3.21 | 1360769 | 4.24 |
| DU018-1 | Ranunculus envalirensis | 471152 | 942304 | 452010 | 11863 | 4279 | 920162 | 2.35 | 834448 | 9.32 |
| Du019 | Ranunculus envalirensis | 618223 | 1236446 | 525997 | 81023 | 2217 | 1135234 | 8.19 | 1120850 | 1.27 |
| DU021-1 | Ranunculus flabellifolius | 796503 | 1593006 | 770210 | 13476 | 8247 | 1562143 | 1.94 | 1370079 | 12.3 |
| DU054-1 | Ranunculus flabellifolius | 563973 | 1127946 | 541083 | 14506 | 4726 | 1101398 | 2.36 | 979444 | 11.08 |
| LH2501 | Ranunculus flabellifolius | 533925 | 1067850 | 505220 | 20250 | 3864 | 1034554 | 3.12 | 956330 | 7.57 |
| LH017-1 | Ranunculus marsicus | 549225 | 1098450 | 527303 | 12906 | 5484 | 1072996 | 2.32 | 967954 | 9.79 |
| LH018-2 | Ranunculus marsicus | 715224 | 1430448 | 680873 | 22667 | 6395 | 1390808 | 2.78 | 1275636 | 8.29 |
| LH014-3 | Ranunculus mediocompositus | 690788 | 1381576 | 650077 | 30227 | 4615 | 1334996 | 3.38 | 1226172 | 8.16 |
| LH015-10 | Ranunculus mediocompositus | 758808 | 1517616 | 729505 | 17696 | 6762 | 1483468 | 2.26 | 1332162 | 10.2 |
| 10137-3 | Ranunculus notabilis | 684406 | 1368812 | 647306 | 26682 | 4787 | 1326081 | 3.13 | 1226859 | 7.49 |
| Hoe5615 | Ranunculus notabilis | 788655 | 1577310 | 653152 | 121175 | 2219 | 1429698 | 9.36 | 1359838 | 4.89 |
| LH010-7 | Ranunculus peracris | 433500 | 867000 | 416171 | 10395 | 4167 | 846904 | 2.32 | 794542 | 6.19 |
| LH011-1 | Ranunculus peracris | 680873 | 1361746 | 638035 | 32232 | 4542 | 1312844 | 3.6 | 1193156 | 9.12 |
| LH011-14 | Ranunculus peracris | 482678 | 965356 | 461588 | 13306 | 4441 | 940923 | 2.54 | 883035 | 6.16 |
| LG09 | Ranunculus pygmaeus | 908098 | 1816196 | 875316 | 19488 | 8040 | 1778160 | 2.1 | 1619494 | 8.93 |
| 104263 | Ranunculus sceleratus | 1047365 | 2094730 | 991745 | 40081 | 7439 | 2031010 | 3.05 | 1886966 | 7.1 |
| DU049-1 | Ranunculus subcarniolicus | 656379 | 1312758 | 631484 | 14967 | 6148 | 1284083 | 2.19 | 1183105 | 7.87 |
| LH013-3 | Ranunculus subcarniolicus | 479816 | 959632 | 463766 | 8596 | 4762 | 940890 | 1.96 | 876464 | 6.85 |
