## Supplementary material for "Phylogenomics unravels speciation patterns in temperate-montane plant species: a case study on the recently radiating *Ranunculus auricomus* species complex": Table S4

**Table S4** Number and percentage of mapped read per sample.

| Sample no. | Taxon | Total nr. reads | Nr. paired reads | Nr. forward unpaired reads | Nr. reverse unpaired reads | Nr. mapped reads | Percentage of mapped reads |
|---|---|---|---|---|---|---|---|
| DU003-1 | Ranunculus austroslovenicus | 828673 | 387692 | 9204 | 4216 | 662966 | 80.003 |
| LH012-8 | Ranunculus austroslovenicus | 1144964 | 535481 | 11436 | 6398 | 935564 | 81.711 |
| 2 | Ranunculus calapius | 1136354 | 542744 | 33091 | 4235 | 305569 | 26.89 |
| Du35351-15 | Ranunculus calapius | 1658283 | 801437 | 20017 | 7997 | 974768 | 58.781 |
| 9126-2 | Ranunculus carpaticola | 1555793 | 758227 | 19697 | 9823 | 857218 | 55.098 |
| DU013-1 | Ranunculus carpaticola | 1449882 | 664408 | 36202 | 5318 | 1079453 | 74.451 |
| LH40-4 | Ranunculus carpaticola | 1100804 | 518871 | 28544 | 4068 | 717409 | 65.171 |
| LH006-17 | Ranunculus cassubicifolius | 1334773 | 624716 | 29304 | 5430 | 910910 | 68.244 |
| LH016-14 | Ranunculus cassubicifolius | 2015353 | 948717 | 20883 | 10570 | 1416897 | 70.305 |
| LH008-7 | Ranunculus cassubicifolius | 1363349 | 646866 | 17458 | 7015 | 927282 | 68.015 |
| LH009-3 | Ranunculus cassubicifolius | 1230345 | 556977 | 77486 | 2341 | 830803 | 67.526 |
| Du33354-2 | Ranunculus cebennensis | 1384000 | 663997 | 26409 | 6366 | 839598 | 60.664 |
| DU018-1 | Ranunculus envalirensis | 880508 | 409153 | 11863 | 4279 | 615685 | 69.923 |
| Du019 | Ranunculus envalirensis | 1147489 | 518805 | 81023 | 2217 | 611987 | 53.332 |
| DU021-1 | Ranunculus flabellifolius | 1424710 | 674178 | 13476 | 8247 | 1015231 | 71.258 |
| DU054-1 | Ranunculus flabellifolius | 1032953 | 480106 | 14506 | 4726 | 768506 | 74.398 |
| LH2501 | Ranunculus flabellifolius | 984472 | 466108 | 20250 | 3864 | 618623 | 62.838 |
| LH017-1 | Ranunculus marsicus | 1015923 | 474782 | 12906 | 5484 | 760511 | 74.859 |
| LH018-2 | Ranunculus marsicus | 1342779 | 623287 | 22667 | 6395 | 946497 | 70.487 |
| LH014-3 | Ranunculus mediocompositus | 1290181 | 595665 | 30227 | 4615 | 1041981 | 80.762 |
| LH015-10 | Ranunculus mediocompositus | 1410906 | 653852 | 17696 | 6762 | 1141346 | 80.894 |
| 10137-3 | Ranunculus notabilis | 1283419 | 597695 | 26682 | 4787 | 1008654 | 78.591 |
| Hoe5615 | Ranunculus notabilis | 1379116 | 618222 | 121175 | 2219 | 863888 | 62.64 |
| LH010-7 | Ranunculus peracris | 825825 | 389990 | 10395 | 4167 | 619412 | 75.005 |
| LH011-1 | Ranunculus peracris | 1254824 | 578191 | 32232 | 4542 | 935436 | 74.547 |
| LH011-14 | Ranunculus peracris | 929053 | 432644 | 13306 | 4441 | 702164 | 75.578 |
| LG09 | Ranunculus pygmaeus | 1715802 | 795983 | 19488 | 8040 | 1415286 | 82.485 |
| 104263 | Ranunculus sceleratus | 1986542 | 919723 | 40081 | 7439 | 1571110 | 79.087 |
| DU049-1 | Ranunculus subcarniolicus | 1254050 | 580995 | 14967 | 6148 | 994326 | 79.289 |
| LH013-3 | Ranunculus subcarniolicus | 920035 | 431553 | 8596 | 4762 | 747517 | 81.248 |
