## Supplementary material for "Phylogenomics unravels speciation patterns in temperate-montane plant species: a case study on the recently radiating *Ranunculus auricomus* species complex": Table S5

**Table S5** Tukey multiple comparisons of means. Significant p values are marked in bolt

| | diff | lwr | upr | p adj |
|---|---|---|---|---|
| AlleleAlign.ParalogFilt.PositionUnfilt.-AlleleAlign.ParalogFilt.PositionFilt. | 0.00098571 | -0.0178477 | 0.01981912 | 0.9999999 |
| AlleleAlign.ParalogUnfilt.PositionFilt.-AlleleAlign.ParalogFilt.PositionFilt. | 0.03172143 | 0.01288802 | 0.05055483 | **0.0000090** |
| AlleleAlign.ParalogUnfilt.PositionUnfilt.-AlleleAlign.ParalogFilt.PositionFilt. | 0.03446429 | 0.01563088 | 0.05329769 | **0.0000006** |
| ConsensusAlign.ParalogFilt.PositionFilt.-AlleleAlign.ParalogFilt.PositionFilt. | -0.0065917 | -0.0255987 | 0.01241532 | 0.9664137 |
| ConsensusAlign.ParalogFilt.PositionUnfilt.-AlleleAlign.ParalogFilt.PositionFilt. | -0.0074324 | -0.0264394 | 0.01157458 | 0.9363370 |
| ConsensusAlign.ParalogUnfilt.PositionFilt.-AlleleAlign.ParalogFilt.PositionFilt. | 0.04046389 | 0.0214569 | 0.05947088 | **0.0000000** |
| ConsensusAlign.ParalogUnfilt.PositionUnfilt.-AlleleAlign.ParalogFilt.PositionFilt. | 0.04115648 | 0.02214949 | 0.06016347 | **0.0000000** |
| AlleleAlign.ParalogUnfilt.PositionFilt.-AlleleAlign.ParalogFilt.PositionUnfilt. | 0.03073571 | 0.01190231 | 0.04956912 | 0.0000207 |
| AlleleAlign.ParalogUnfilt.PositionUnfilt.-AlleleAlign.ParalogFilt.PositionUnfilt. | 0.03347857 | 0.01464517 | 0.05231198 | 0.0000018 |
| ConsensusAlign.ParalogFilt.PositionFilt.-AlleleAlign.ParalogFilt.PositionUnfilt. | -0.0075774 | -0.0265844 | 0.01142961 | 0.9297382 |
| ConsensusAlign.ParalogFilt.PositionUnfilt.-AlleleAlign.ParalogFilt.PositionUnfilt. | -0.0084181 | -0.0274251 | 0.01058887 | 0.8825819 |
| ConsensusAlign.ParalogUnfilt.PositionFilt.-AlleleAlign.ParalogFilt.PositionUnfilt. | 0.03947817 | 0.02047119 | 0.05848516 | **0.0000000** |
| ConsensusAlign.ParalogUnfilt.PositionUnfilt.-AlleleAlign.ParalogFilt.PositionUnfilt. | 0.04017077 | 0.02116378 | 0.05917776 | **0.0000000** |
| AlleleAlign.ParalogUnfilt.PositionUnfilt.-AlleleAlign.ParalogUnfilt.PositionFilt. | 0.00274286 | -0.0160905 | 0.02157626 | 0.9998539 |
| ConsensusAlign.ParalogFilt.PositionFilt.-AlleleAlign.ParalogUnfilt.PositionFilt. | -0.0383131 | -0.0573201 | -0.0193061 | **0.0000000** |
| ConsensusAlign.ParalogFilt.PositionUnfilt.-AlleleAlign.ParalogUnfilt.PositionFilt. | -0.0391538 | -0.0581608 | -0.0201469 | **0.0000000** |
| ConsensusAlign.ParalogUnfilt.PositionFilt.-AlleleAlign.ParalogUnfilt.PositionFilt. | 0.00874246 | -0.0102645 | 0.02774945 | 0.8602767 |
| ConsensusAlign.ParalogUnfilt.PositionUnfilt.-AlleleAlign.ParalogUnfilt.PositionFilt. | 0.00943505 | -0.0095719 | 0.02844204 | 0.8052769 |
| ConsensusAlign.ParalogFilt.PositionFilt.-AlleleAlign.ParalogUnfilt.PositionUnfilt. | -0.041056 | -0.0600629 | -0.022049 | **0.0000000** |
| ConsensusAlign.ParalogFilt.PositionUnfilt.-AlleleAlign.ParalogUnfilt.PositionUnfilt. | -0.0418967 | -0.0609037 | -0.0228897 | **0.0000000** |
| ConsensusAlign.ParalogUnfilt.PositionFilt.-AlleleAlign.ParalogUnfilt.PositionUnfilt. | 0.0059996 | -0.0130074 | 0.02500659 | 0.9802237 |
| ConsensusAlign.ParalogUnfilt.PositionUnfilt.-AlleleAlign.ParalogUnfilt.PositionUnfilt. | 0.0066922 | -0.0123148 | 0.02569918 | 0.9635074 |
| ConsensusAlign.ParalogFilt.PositionUnfilt.-ConsensusAlign.ParalogFilt.PositionFilt. | -0.0008407 | -0.0200197 | 0.01833826 | 1.0000000 |
| ConsensusAlign.ParalogUnfilt.PositionFilt.-ConsensusAlign.ParalogFilt.PositionFilt. | 0.04705556 | 0.02787656 | 0.06623456 | **0.0000000** |
| ConsensusAlign.ParalogUnfilt.PositionUnfilt.-ConsensusAlign.ParalogFilt.PositionFilt. | 0.04774815 | 0.02856915 | 0.06692715 | **0.0000000** |
| ConsensusAlign.ParalogUnfilt.PositionFilt.-ConsensusAlign.ParalogFilt.PositionUnfilt. | 0.0478963 | 0.0287173 | 0.0670753 | **0.0000000** |
| ConsensusAlign.ParalogUnfilt.PositionUnfilt.-ConsensusAlign.ParalogFilt.PositionUnfilt. | 0.04858889 | 0.02940989 | 0.06776789 | **0.0000000** |
| ConsensusAlign.ParalogUnfilt.PositionUnfilt.-ConsensusAlign.ParalogUnfilt.PositionFilt. | 0.00069259 | -0.0184864 | 0.01987159 | 1.0000000 |
