## Supplementary material for "Phylogenomics unravels speciation patterns in temperate-montane plant species: a case study on the recently radiating *Ranunculus auricomus* species complex": Table S6

**Table S6** Results from the Likelihood Ratio Test (LRT) of different nested models.

| alt | null | LnLalt | LnLnull | DFalt | DFnull | DF | Dstatistic | pval | test | tail | AIC1 | AIC2 | AICwt1 | AICwt2 | AICweight ratio model1 | AICweight ratio model2 |
|---|---|---|---|---|---|---|---|---|---|---|---|---|---|---|---|---|
| DEC+x | DEC | -22.756713 | -26.228416 | 3 | 2 | 1 | 6.94340487 | 0.00841288 | chi-squared | one-tailed | 51.51343 | 56.45683 | 0.9221341 | 0.07786591 | 11.84259 | 0.08444098 |
| DEC+j | DEC | -24.273943 | -26.228416 | 3 | 2 | 1 | 3.90894538 | 0.04802973 | chi-squared | one-tailed | 54.54789 | 56.45683 | 0.7220138 | 0.2779862 | 2.597301 | 0.3850151 |
| DEC+x+j | DEC+x | -22.597258 | -22.756713 | 4 | 3 | 1 | 0.31891002 | 0.57226342 | chi-squared | one-tailed | 53.19452 | 51.51343 | 0.30142 | 0.69858 | 0.4314753 | 2.31763 |
| DIVA+x | DIVA | -18.306908 | -21.475135 | 3 | 2 | 1 | 6.3364542 | 0.01182811 | chi-squared | one-tailed | 42.61382 | 46.95027 | 0.8973598 | 0.1026402 | 8.74277 | 0.1143802 |
| DIVA+j | DIVA | -21.473241 | -21.475135 | 3 | 2 | 1 | 0.00378824 | 0.95092226 | chi-squared | one-tailed | 48.94648 | 46.95027 | 0.269314 | 0.730686 | 0.3685769 | 2.713138 |
| DIVA+x+j | DIVA+x | -17.718453 | -18.306908 | 4 | 3 | 1 | 1.17690952 | 0.27798605 | chi-squared | one-tailed | 43.43691 | 42.61382 | 0.3985417 | 0.6014583 | 0.6626255 | 1.509148 |
| BAYAREA+x | BAYAREA | -29.506118 | -31.871805 | 3 | 2 | 1 | 4.73137481 | 0.0296172 | chi-squared | one-tailed | 65.01224 | 67.74361 | 0.7966825 | 0.2033175 | 3.918416 | 0.2552052 |
| BAYAREA+j | BAYAREA | -26.981695 | -31.871805 | 3 | 2 | 1 | 9.78021918 | 0.00176399 | chi-squared | one-tailed | 59.96339 | 67.74361 | 0.9799664 | 0.02003356 | 48.91625 | 0.02044311 |
| BAYAREA+x+j | BAYAREA+x | -25.354431 | -29.506118 | 4 | 3 | 1 | 8.30337279 | 0.00395715 | chi-squared | one-tailed | 58.70886 | 65.01224 | 0.9589751 | 0.04102488 | 23.37545 | 0.04277992 |
