## Supplementary material for "Phylogenomics unravels speciation patterns in temperate-montane plant species: a case study on the recently radiating *Ranunculus auricomus* species complex": Table S7

**Table S7** Detailed information about positions (along with their estimated substitution rate [illegible]
position filtering for the different datasets

a) Consensus Alignments:

| Locus | position | rate |
|---|---|---|
| **Assembly_AT1G09280** | 1522 | 84.1606 |
| **Assembly_AT1G09280** | 1494 | 84.1606 |
| **Assembly_AT1G09280** | 1492 | 84.1606 |
| **Assembly_AT1G09280** | 1479 | 84.1606 |
| **Assembly_AT1G09280** | 1470 | 84.1606 |
| **Assembly_AT1G09280** | 1468 | 84.1606 |
| **Assembly_AT1G09280** | 1458 | 84.1606 |
| **Assembly_AT1G09280** | 1457 | 84.1606 |
| **Assembly_AT1G09280** | 1456 | 84.1606 |
| **Assembly_AT1G09280** | 1454 | 84.1606 |
| **Assembly_AT1G09280** | 1453 | 84.1606 |
| **Assembly_AT1G09280** | 1452 | 84.1606 |
| **Assembly_AT1G09280** | 1451 | 84.1606 |
| **Assembly_AT1G09280** | 1449 | 84.1606 |
| **Assembly_AT1G09280** | 1447 | 84.1606 |
| **Assembly_AT1G09280** | 1446 | 84.1606 |
| **Assembly_AT1G09280** | 1445 | 84.1606 |
| **Assembly_AT1G09280** | 1444 | 84.1606 |
| **Assembly_AT1G09850** | 344 | 230.3317 |
| **Assembly_AT1G21370** | 374 | 716.5574 |
| **Assembly_AT1G31970** | 279 | 84.1606 |
| **Assembly_AT1G31970** | 267 | 84.1606 |
| **Assembly_AT1G75210** | 662 | 84.1606 |
| **Assembly_AT1G75210** | 13 | 55.6806 |
| **Assembly_AT1G78280** | 2444 | 190.5341 |
| **Assembly_AT2G03430** | 394 | 55.0661 |
| **Assembly_AT2G38060** | 587 | 178.2413 |
| **Assembly_AT2G38060** | 584 | 440.6491 |
| **Assembly_AT2G38060** | 564 | 91.2023 |
| **Assembly_AT2G38060** | 563 | 81.385 |
| **Assembly_AT2G38060** | 476 | 1966.7312 |
| **Assembly_AT2G38060** | 475 | 1724.6578 |
| **Assembly_AT2G38060** | 466 | 220.3121 |
| **Assembly_AT3G11830** | 788 | 1217.649 |
| **Assembly_AT3G12940** | 202 | 84.1606 |
| **Assembly_AT3G55260** | 523 | 84.1606 |
| **Assembly_AT3G55260** | 462 | 84.1606 |
| **Assembly_AT3G55260** | 461 | 84.1606 |
| **Assembly_AT3G55260** | 446 | 178.3747 |
| **Assembly_AT3G55260** | 440 | 84.1606 |
| **Assembly_AT3G55260** | 414 | 84.1606 |
| **Assembly_AT3G55260** | 407 | 84.1606 |
| **Assembly_AT3G55260** | 404 | 84.1606 |
| **Assembly_AT4G13590** | 594 | 236.9334 |
| **Assembly_AT4G13590** | 593 | 265.9859 |
| **Assembly_AT4G13590** | 592 | 236.9334 |
| **Assembly_AT4G13590** | 591 | 236.9334 |
| **Assembly_AT4G13590** | 590 | 197.9026 |
| **Assembly_AT4G13590** | 589 | 236.9334 |
| **Assembly_AT4G13590** | 588 | 197.9026 |
| **Assembly_AT4G13590** | 587 | 265.9859 |
| **Assembly_AT4G13590** | 586 | 236.9334 |
| **Assembly_AT4G13590** | 584 | 197.9026 |
| **Assembly_AT4G13590** | 583 | 197.9026 |
| **Assembly_AT4G13590** | 582 | 236.9334 |

| | | |
|---|---|---|
| **Assembly_AT4G13590** | 581 | 265.9859 |
| **Assembly_AT4G13590** | 580 | 265.9859 |
| **Assembly_AT4G20325** | 378 | 229.9387 |
| **Assembly_AT4G20325** | 377 | 223.273 |
| **Assembly_AT4G30310** | 882 | 101.4457 |
| **Assembly_AT5G08720** | 289 | 84.1606 |
| **Assembly_AT5G08720** | 244 | 84.1606 |
| **Assembly_AT5G08720** | 234 | 84.1606 |
| **Assembly_AT5G20040** | 522 | 1762.6633 |
| **Assembly_AT5G21070** | 1152 | 84.1606 |
| **Assembly_AT5G21070** | 1134 | 84.1606 |
| **Assembly_AT5G48520** | 737 | 84.1606 |
| **Assembly_AT5G48520** | 736 | 84.1606 |
| **Assembly_AT5G48520** | 735 | 84.1606 |
| **Assembly_AT5G48520** | 734 | 84.1606 |
| **Assembly_AT5G48520** | 733 | 84.1606 |
| **Assembly_AT5G48520** | 731 | 84.1606 |
| **Assembly_AT5G48520** | 730 | 84.1606 |
| **Assembly_AT5G64940** | 828 | 84.1606 |
| **Assembly_AT5G64940** | 795 | 84.1606 |
| **Assembly_AT5G64940** | 637 | 84.1606 |
| **Assembly_AT5G66120** | 405 | 884.0018 |

b) Consensus alignments, paralog filtering

| Locus | position | rate |
|---|---|---|
| **Assembly_AT1G31970** | 279 | 84.1606 |
| **Assembly_AT1G31970** | 267 | 84.1606 |
| **Assembly_AT1G78280** | 2444 | 190.5341 |
| **Assembly_AT3G55260** | 523 | 84.1606 |
| **Assembly_AT3G55260** | 462 | 84.1606 |
| **Assembly_AT3G55260** | 461 | 84.1606 |
| **Assembly_AT3G55260** | 446 | 178.3747 |
| **Assembly_AT3G55260** | 440 | 84.1606 |
| **Assembly_AT3G55260** | 414 | 84.1606 |
| **Assembly_AT3G55260** | 407 | 84.1606 |
| **Assembly_AT3G55260** | 404 | 84.1606 |
| **Assembly_AT4G20325** | 378 | 229.9387 |
| **Assembly_AT4G20325** | 377 | 223.273 |
| **Assembly_AT5G21070** | 1152 | 84.1606 |
| **Assembly_AT5G21070** | 1134 | 84.1606 |
| **Assembly_AT5G66120** | 405 | 884.0018 |

c) Allele alignments

| Exons | position | rate |
|---|---|---|
| **Assembly_AT1G21370_13** | 78 | 110.5288 |
| **Assembly_AT2G18940_1** | 409 | 62.854 |
| **Assembly_AT2G18940_1** | 349 | 86.7073 |
| **Assembly_AT2G18940_49** | 409 | 62.854 |
| **Assembly_AT2G18940_49** | 349 | 86.7073 |
| **Assembly_AT3G04870_2376** | 6 | 48.9575 |
| **Assembly_AT4G17300_3349** | 293 | 1572.8704 |
| **Assembly_AT4G17300_3349** | 284 | 786.4385 |
| **Assembly_AT4G33060_1179** | 4 | 75.0819 |
| **Assembly_AT5G01010_17** | 93 | 221.1661 |
| **Assembly_AT5G01010_17** | 89 | 263.8743 |
| **Assembly_AT5G14600_85** | 36 | 98.6839 |
| **Assembly_AT5G64050_4** | 287 | 101.2217 |

d) Allele alignments, paralog filtering

| Exons | position | rate |
|---|---|---|
| **Assembly_AT2G18940_1** | 409 | 262.9397 |
| **Assembly_AT2G18940_1** | 349 | 443.5075 |
| **Assembly_AT2G18940_49** | 409 | 262.9397 |
| **Assembly_AT2G18940_49** | 349 | 443.5075 |
| **Assembly_AT3G04870_2376** | 6 | 76.6957 |
| **Assembly_AT4G17300_3349** | 293 | 179.3766 |
| **Assembly_AT4G17300_3349** | 284 | 229.6578 |
| **Assembly_AT4G33060_1179** | 4 | 118.3271 |
| **Assembly_AT5G19850_953** | 37 | 22.0063 |
