## Supplementary material for "Phylogenomics unravels speciation patterns in temperate-montane plant species: a case study on the recently radiating *Ranunculus auricomus* species complex": Figure S1

**Firure S1** Boxplot showing performance (mean Local Posterior Probability of the bootstrap ASTRAL trees) of the different phasing and filtering schemes.

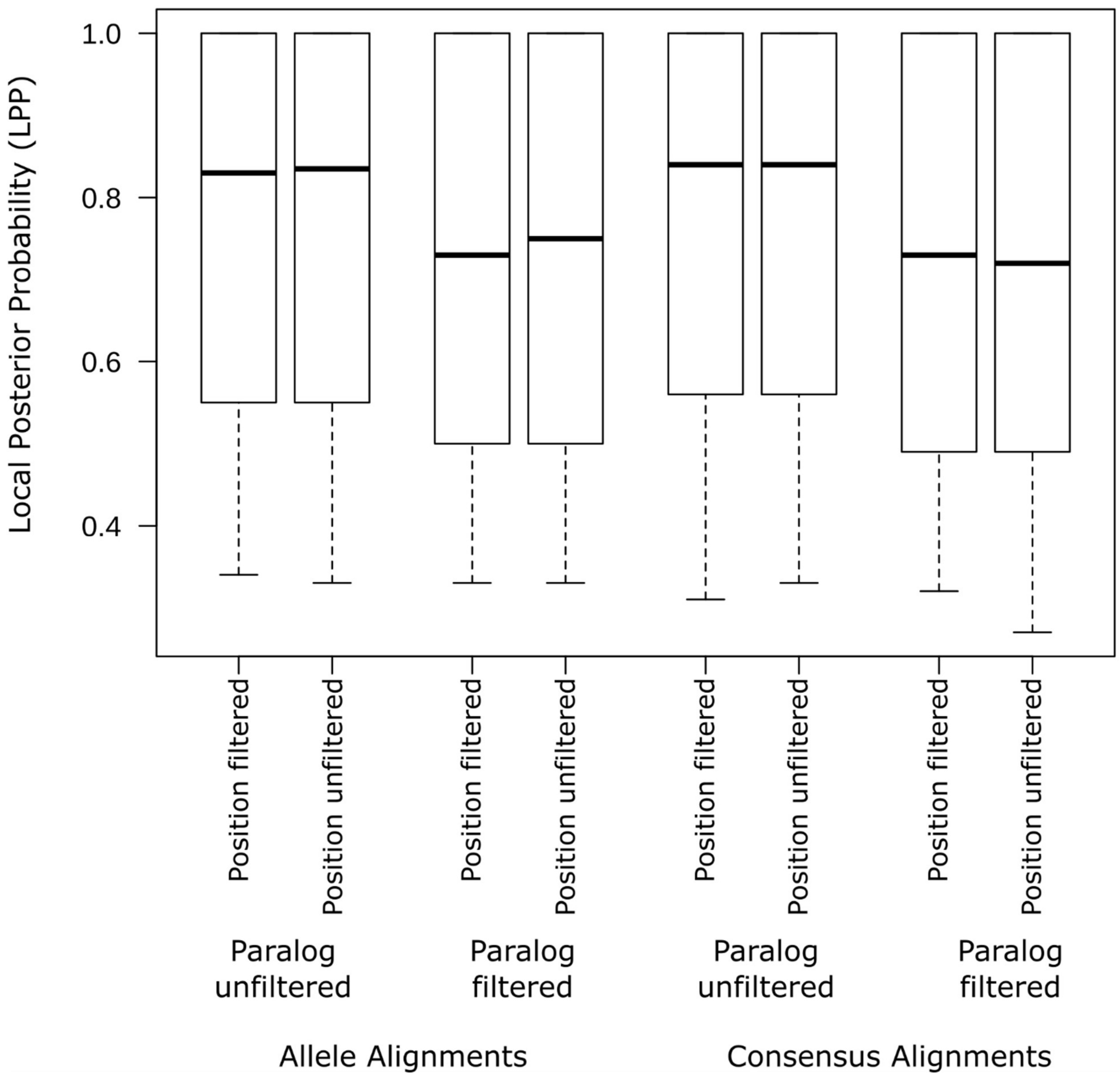
