## Supplementary material for "Phylogenomics unravels speciation patterns in temperate-montane plant species: a case study on the recently radiating *Ranunculus auricomus* species complex": Figures S2-S7

**Figure S2** Phylogenetic trees drawn opposite to each other (tanglegram) from the coalescent-based Astral species tree (left) and concatenated maximum likelihood analysis (right), based on the consensus dataset after applying position and paralog filtering. Continuous lines indicate accession with congruent position, whereas dashed lines indicate incongruence. Circles on the nodes of the trees refer to the bootstrap support values. Blue bigger circles are for support values above 95; purples medium-sized circles are for values between 75 and 95; small red circles are for bootstraps below 75. Numbers in proximity of nodes indicate ASTRAL quartet supports (left) and results from quartet sampling on the maximum likelihood tree (right).

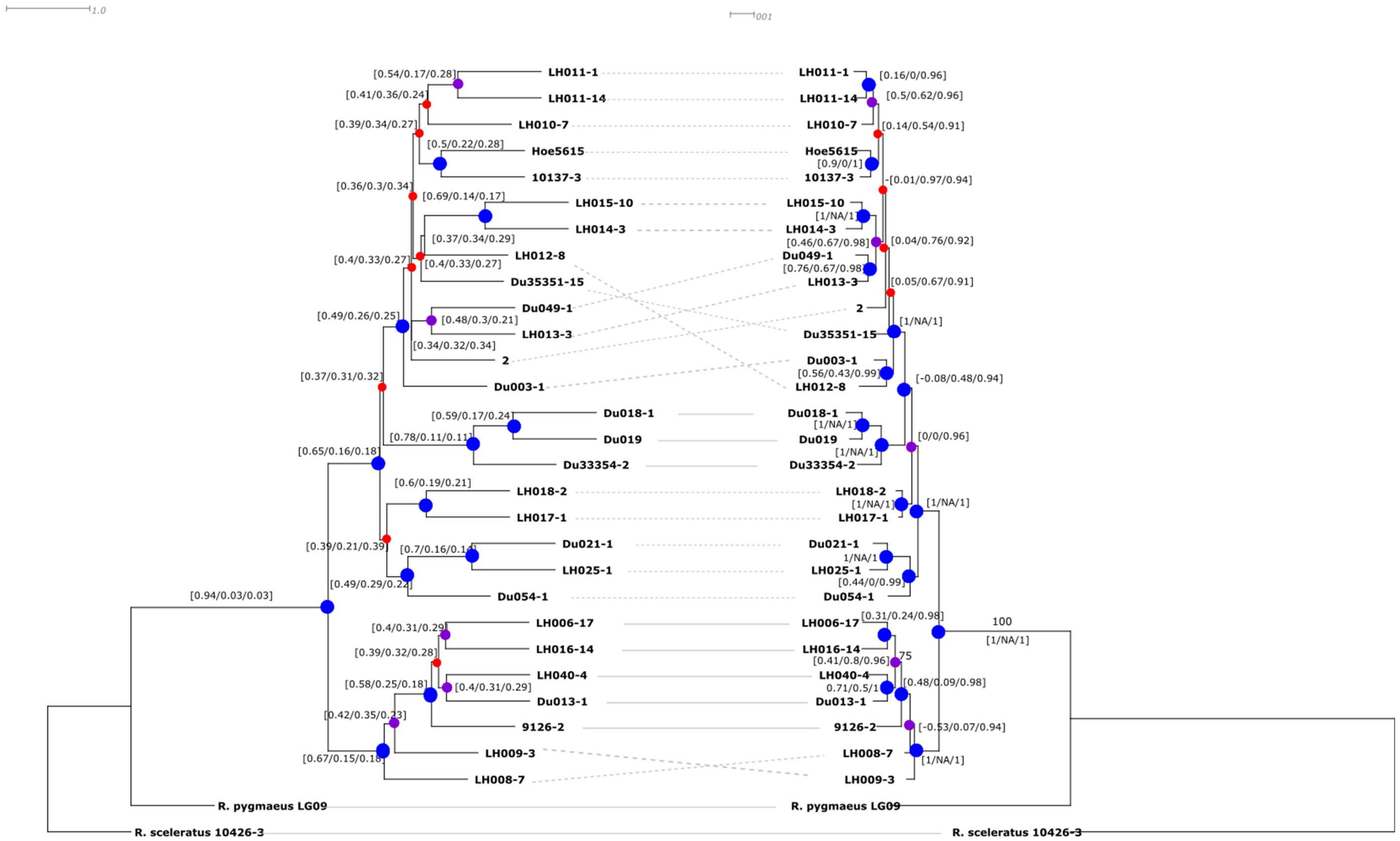

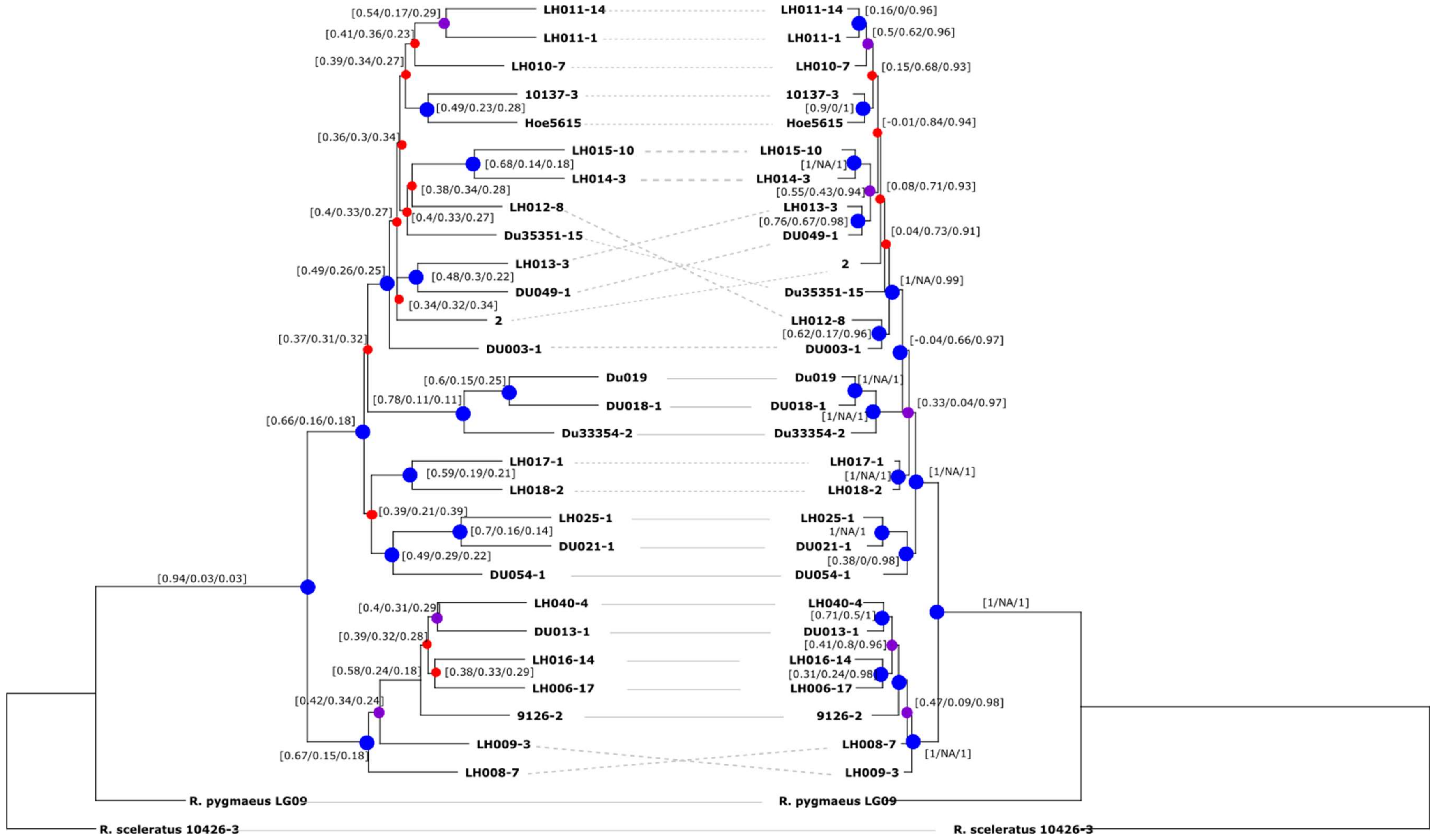


**Figure S3** Phylogenetic trees drawn opposite to each other (tanglegram) from the coalescent-based Astral species tree (left) and concatenated maximum likelihood analysis (right), based on the consensus dataset after applying and paralog filtering. Continuous lines indicate accession with congruent position, whereas dashed lines indicate incongruence. Circles on the nodes of the trees refer to the bootstrap support values. Blue bigger circles are for support values above 95; purples medium -ized circles are for values between 75 and 95; small red circles are for bootstraps below 75. Numbers in proximity of nodes indicate ASTRAL quartet supports (left) and results from quartet sampling on the maximum likelihood tree (right).

**Figure S4** Phylogenetic trees drawn opposite to each other (tanglegram) from the coalescent-based Astral species tree (left) and concatenated maximum likelihood analysis (right), based on the consensus dataset after applying position filtering and without paralog filtering. Continuous lines indicate accession with congruent position, whereas dashed lines indicate incongruence. Circles on the nodes of the trees refer to the bootstrap support values. Blue bigger circles are for support values above 95; purples medium-sized circles are for values between 75 and 95; small red circles are for bootstraps below 75. Numbers in proximity of nodes indicate ASTRAL quartet supports (left) and results from quartet sampling on the maximum likelihood tree (right).

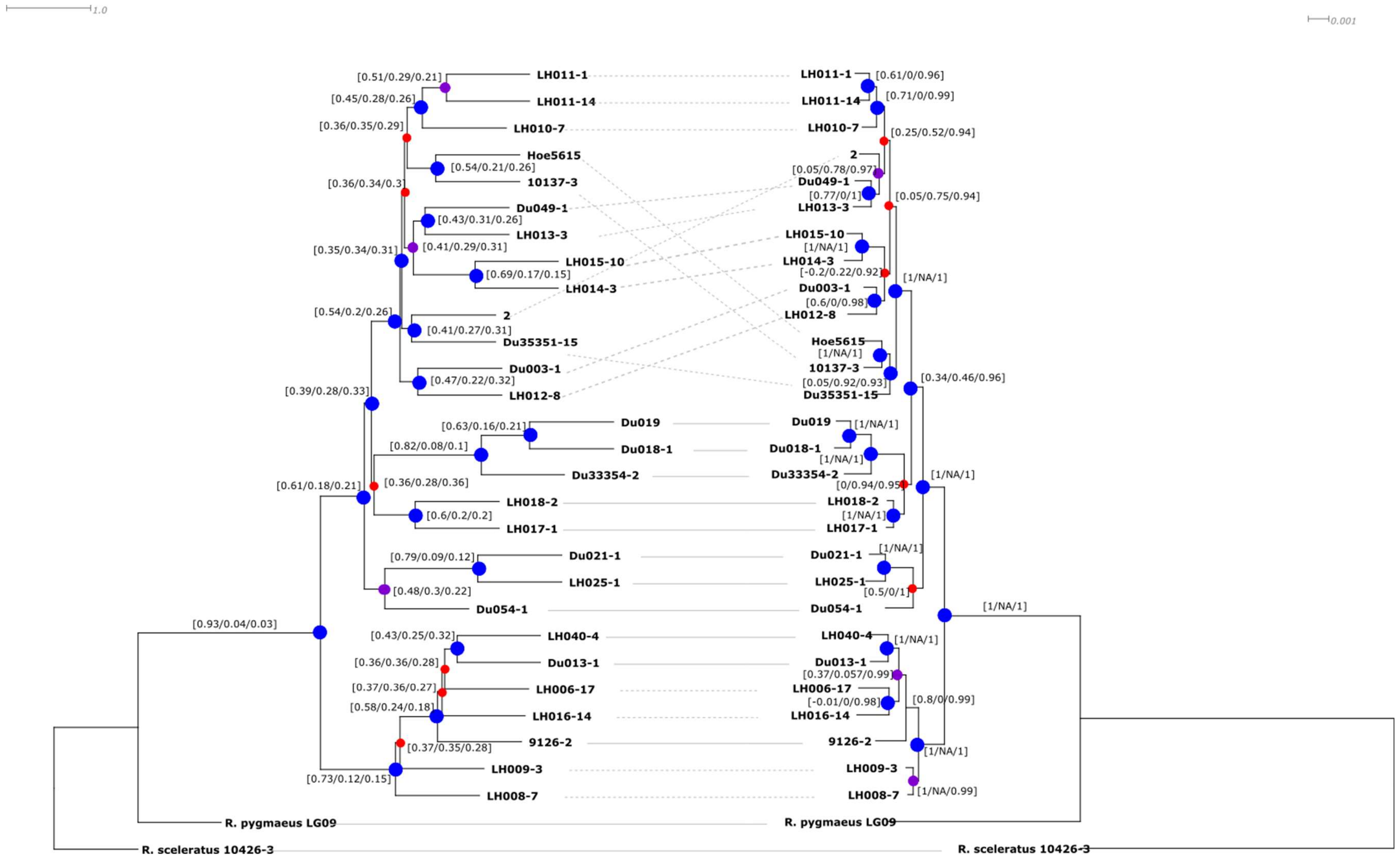

**Figure S5** Astral species tree based on the allele dataset after applying position and paralog filtering. Numbers above branches are bootstrap values (only above 75 are shown). Numbers in square brackets in proximity of nodes indicate ASTRAL quartet supports for the main/first alternative/second alternative topologies.

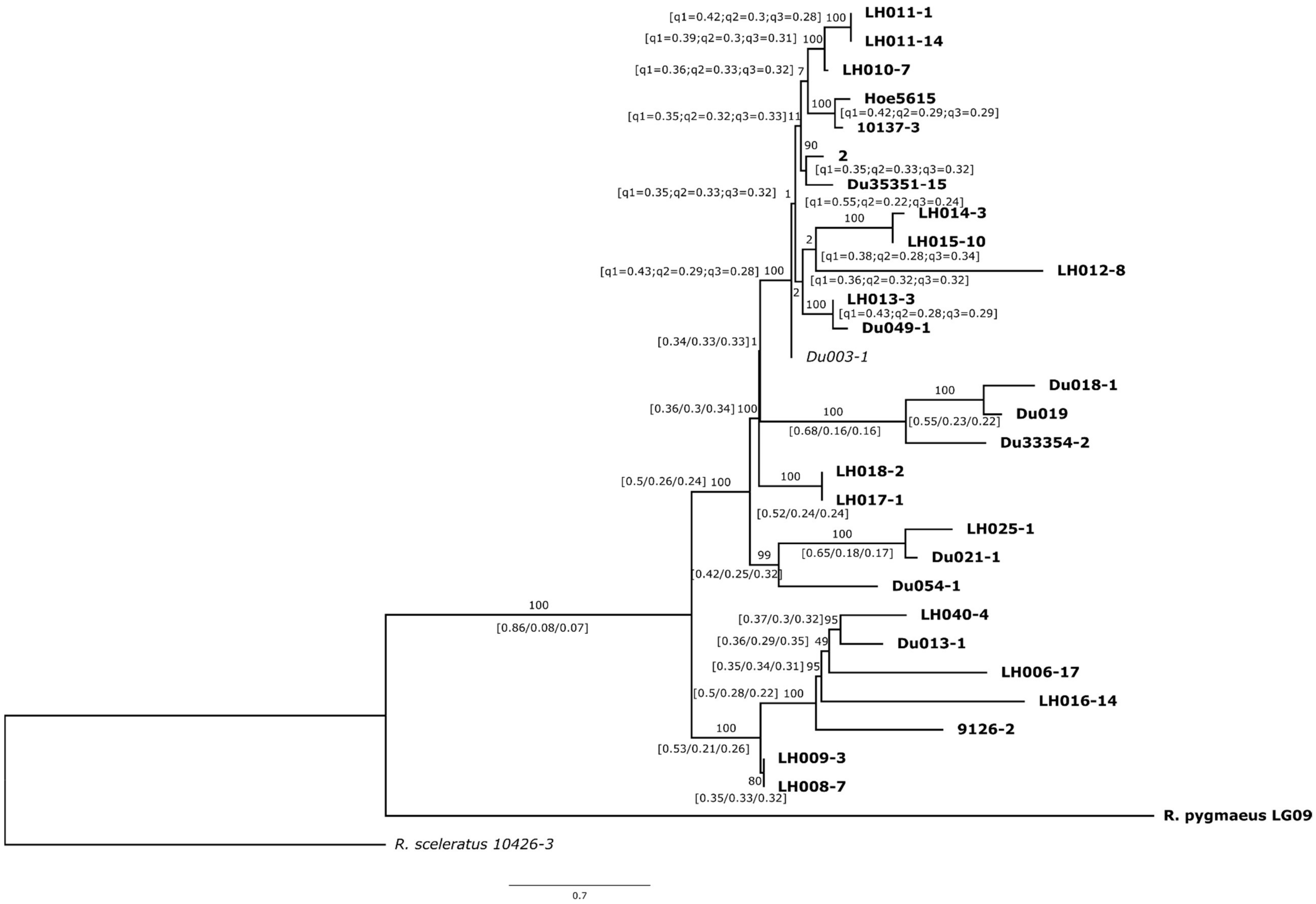

**Figure S6** Astral species tree based on the allele dataset after applying paralog filtering. Numbers above branches are bootstrap values (only above 75 are shown). Numbers in square brackets in proximity of nodes indicate ASTRAL quartet supports for the main/first alternative/second alternative topologies.

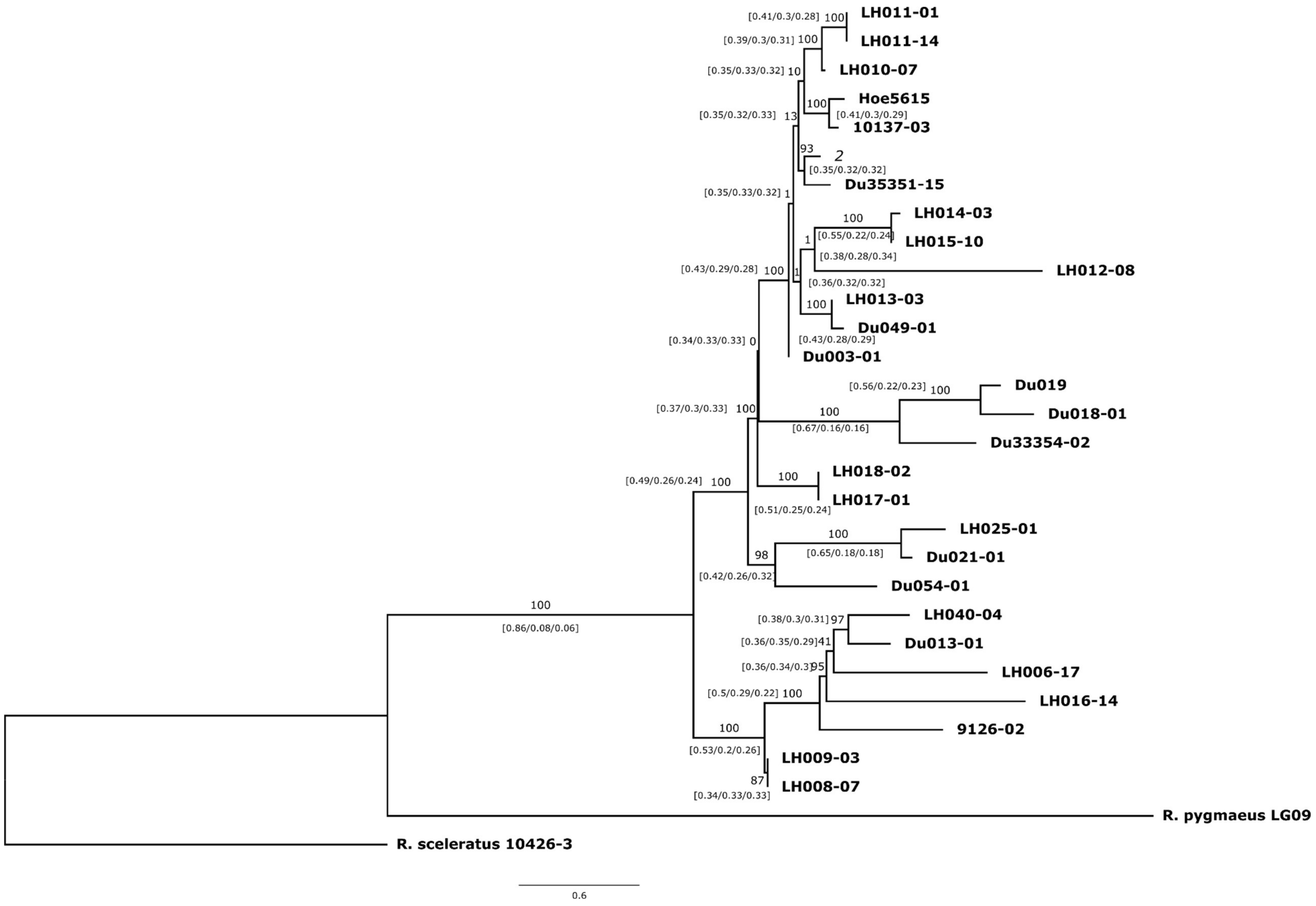

**Figure S7** Astral species tree based on the allele dataset after applying position filtering and without paralog filtering. Numbers above branches are bootstrap values (only above 75 are shown). Numbers in square brackets in proximity of nodes indicate ASTRAL quartet supports for the main/first alternative/second alternative topologies.

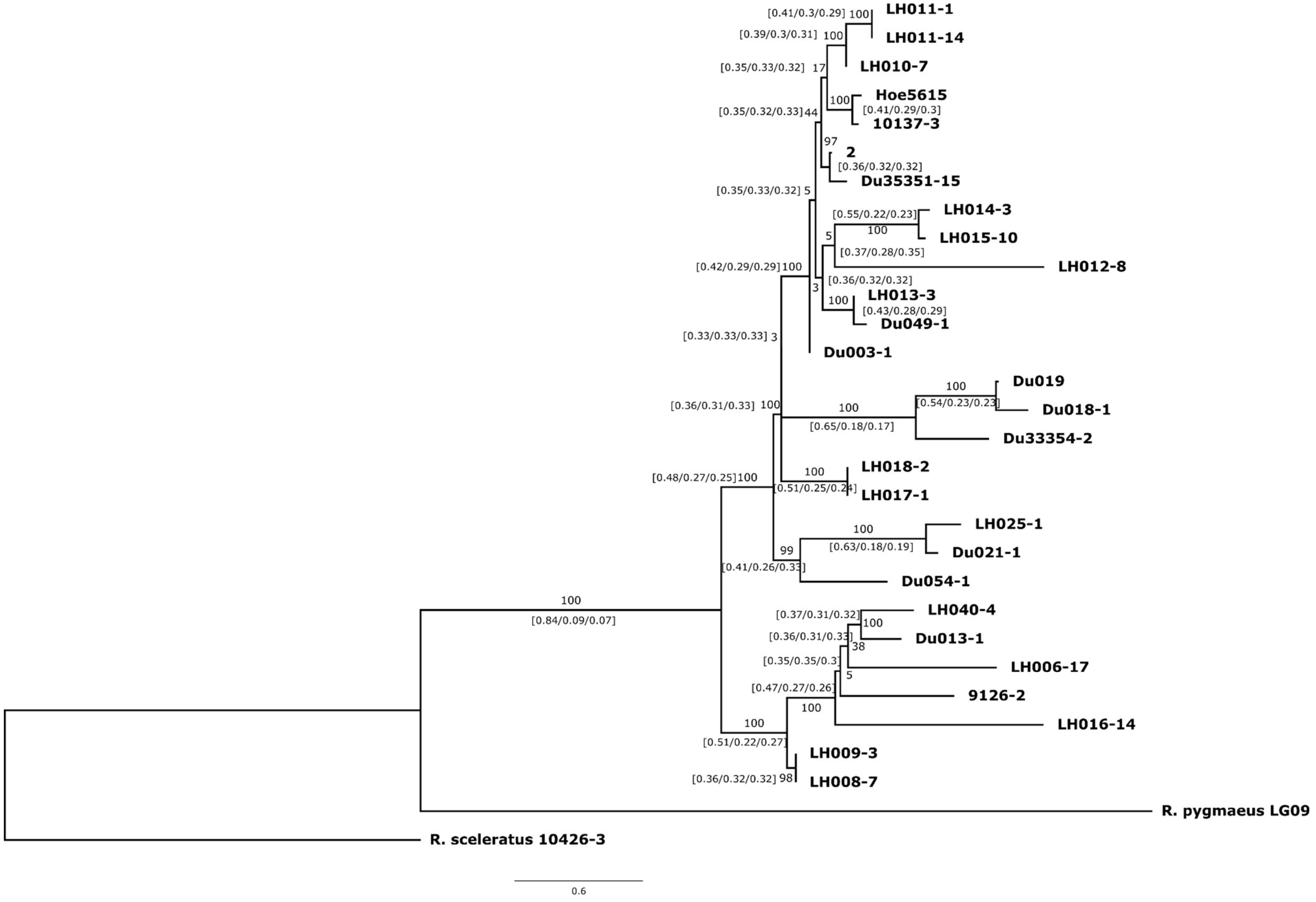
