## Supplementary material for "Phylogenomics unravels speciation patterns in temperate-montane plant species: a case study on the recently radiating *Ranunculus auricomus* species complex": Table S2

**Table S2** The 50 loci with the highest number of Parsimony Informative sites along with their best-fitting substitution model and characteristics.

| | | Model | f(a) | f(c) | f(g) | f(t) | kappa | titv | Ra | Rb | Rc | Rd | Re | Rf | pInv | gamma |
|---|---|---|---|---|---|---|---|---|---|---|---|---|---|---|---|---|
| AT1G01090 | BIC | HKY+I | 0.26 | 0.14 | 0.28 | 0.32 | 10.75 | 5.1 | 1 | 10.745 | 1 | 1 | 10.745 | 1 | 0.87 | N/A |
| AT1G02910 | BIC | HKY+I | 0.28 | 0.22 | 0.21 | 0.29 | 4.33 | 2.12 | 1 | 4.329 | 1 | 1 | 4.329 | 1 | 0.9 | N/A |
| AT1G06560 | BIC | TrN+I | 0.29 | 0.19 | 0.26 | 0.26 | 0 | 0 | 1 | 1.819 | 1 | 1 | 4.161 | 1 | 0.82 | N/A |
| AT1G08660 | BIC | TrN+I | 0.3 | 0.2 | 0.22 | 0.28 | 0 | 0 | 1 | 1.564 | 1 | 1 | 3.843 | 1 | 0.86 | N/A |
| AT1G23400 | BIC | HKY+I+G | 0.31 | 0.21 | 0.21 | 0.28 | 3.77 | 1.83 | 1 | 3.772 | 1 | 1 | 3.772 | 1 | 0.81 | 0.8 |
| AT1G28680 | BIC | K80+I+G | 0.25 | 0.25 | 0.25 | 0.25 | 4.26 | 2.13 | 1 | 4.26 | 1 | 1 | 4.26 | 1 | 0.85 | 0.65 |
| AT1G30360 | BIC | HKY+I | 0.28 | 0.21 | 0.21 | 0.3 | 2.43 | 1.18 | 1 | 2.425 | 1 | 1 | 2.425 | 1 | 0.84 | N/A |
| AT1G31480 | BIC | HKY+I | 0.3 | 0.17 | 0.24 | 0.29 | 4.03 | 1.96 | 1 | 4.027 | 1 | 1 | 4.027 | 1 | 0.9 | N/A |
| AT1G32080 | BIC | TPM3uf+I | 0.25 | 0.22 | 0.21 | 0.31 | 0 | 0 | 4.094 | 5.241 | 1 | 4.094 | 5.241 | 1 | 0.86 | N/A |
| AT1G57770 | BIC | HKY+I | 0.27 | 0.19 | 0.23 | 0.31 | 4.94 | 2.4 | 1 | 4.943 | 1 | 1 | 4.943 | 1 | 0.94 | N/A |
| AT1G68100 | BIC | TrN+G | 0.29 | 0.2 | 0.21 | 0.3 | 0 | 0 | 1 | 2.58 | 1 | 1 | 5.425 | 1 | N/A | 0.02 |
| AT1G69500 | BIC | HKY+I | 0.32 | 0.2 | 0.2 | 0.28 | 3.66 | 1.75 | 1 | 3.659 | 1 | 1 | 3.659 | 1 | 0.84 | N/A |
| AT1G74640 | BIC | TPM2uf+I | 0.29 | 0.21 | 0.22 | 0.28 | 0 | 0 | 4.181 | 10.045 | 4.181 | 1 | 10.045 | 1 | 0.87 | N/A |
| AT1G78280 | BIC | HKY+I | 0.3 | 0.19 | 0.23 | 0.29 | 2.7 | 1.32 | 1 | 2.699 | 1 | 1 | 2.699 | 1 | 0.85 | N/A |
| AT2G13100 | BIC | HKY+G | 0.23 | 0.14 | 0.27 | 0.36 | 2.44 | 1.1 | 1 | 2.438 | 1 | 1 | 2.438 | 1 | N/A | 0.02 |
| AT2G21370 | BIC | K80+I | 0.25 | 0.25 | 0.25 | 0.25 | 4.27 | 2.14 | 1 | 4.273 | 1 | 1 | 4.273 | 1 | 0.84 | N/A |
| AT2G21720 | BIC | HKY+I | 0.32 | 0.18 | 0.23 | 0.27 | 4.58 | 2.26 | 1 | 4.583 | 1 | 1 | 4.583 | 1 | 0.84 | N/A |
| AT2G28070 | BIC | HKY+I | 0.27 | 0.21 | 0.21 | 0.31 | 3.05 | 1.49 | 1 | 3.051 | 1 | 1 | 3.051 | 1 | 0.89 | N/A |
| AT2G34640 | BIC | HKY+I+G | 0.31 | 0.2 | 0.25 | 0.24 | 2.24 | 1.14 | 1 | 2.242 | 1 | 1 | 2.242 | 1 | 0.7 | 0.9 |
| AT2G40190 | BIC | HKY+I | 0.3 | 0.19 | 0.22 | 0.3 | 3.83 | 1.85 | 1 | 3.83 | 1 | 1 | 3.83 | 1 | 0.9 | N/A |
| AT2G45500 | BIC | HKY+I | 0.33 | 0.19 | 0.19 | 0.28 | 3.47 | 1.65 | 1 | 3.466 | 1 | 1 | 3.466 | 1 | 0.87 | N/A |
| AT3G04480 | BIC | HKY+I | 0.29 | 0.21 | 0.19 | 0.31 | 5.28 | 2.53 | 1 | 5.278 | 1 | 1 | 5.278 | 1 | 0.86 | N/A |
| AT3G07720 | BIC | TrNef+I | 0.25 | 0.25 | 0.25 | 0.25 | 0 | 0 | 1 | 1.995 | 1 | 1 | 5.547 | 1 | 0.67 | N/A |
| AT3G23940 | BIC | TrN+I+G | 0.31 | 0.17 | 0.24 | 0.28 | 0 | 0 | 1 | 1.439 | 1 | 1 | 5.935 | 1 | 0.75 | 0.6 |
| AT3G27530 | BIC | HKY+I | 0.32 | 0.18 | 0.24 | 0.27 | 3.75 | 1.85 | 1 | 3.746 | 1 | 1 | 3.746 | 1 | 0.89 | N/A |
| AT3G44880 | BIC | TPM3+I | 0.25 | 0.25 | 0.25 | 0.25 | 0 | 0 | 2.556 | 5.89 | 1 | 2.556 | 5.89 | 1 | 0.88 | N/A |
| AT3G54090 | BIC | HKY+I | 0.3 | 0.2 | 0.22 | 0.28 | 3.13 | 1.53 | 1 | 3.129 | 1 | 1 | 3.129 | 1 | 0.86 | N/A |
| AT3G54510 | BIC | HKY | 0.27 | 0.19 | 0.21 | 0.34 | 4.07 | 1.94 | 1 | 4.074 | 1 | 1 | 4.074 | 1 | N/A | N/A |
| AT3G58470 | BIC | HKY+I | 0.3 | 0.18 | 0.24 | 0.28 | 5.47 | 2.69 | 1 | 5.472 | 1 | 1 | 5.472 | 1 | 0.88 | N/A |
| AT3G59040 | BIC | HKY+I | 0.33 | 0.17 | 0.23 | 0.28 | 3.69 | 1.81 | 1 | 3.69 | 1 | 1 | 3.69 | 1 | 0.87 | N/A |
| AT3G59770 | BIC | HKY+G | 0.26 | 0.21 | 0.23 | 0.29 | 3.89 | 1.92 | 1 | 3.892 | 1 | 1 | 3.892 | 1 | N/A | 0.02 |
| AT3G61320 | BIC | K80+I+G | 0.25 | 0.25 | 0.25 | 0.25 | 2.19 | 1.09 | 1 | 2.187 | 1 | 1 | 2.187 | 1 | 0.83 | 0.88 |
| AT4G13250 | BIC | HKY+I | 0.31 | 0.19 | 0.23 | 0.27 | 3.56 | 1.76 | 1 | 3.564 | 1 | 1 | 3.564 | 1 | 0.93 | N/A |
| AT4G15850 | BIC | HKY | 0.27 | 0.19 | 0.23 | 0.31 | 2.51 | 1.22 | 1 | 2.51 | 1 | 1 | 2.51 | 1 | N/A | N/A |
| AT4G19860 | BIC | HKY+I | 0.31 | 0.17 | 0.24 | 0.28 | 3.2 | 1.58 | 1 | 3.201 | 1 | 1 | 3.201 | 1 | 0.79 | N/A |
| AT4G27340 | BIC | TPM2uf+I | 0.29 | 0.14 | 0.27 | 0.3 | 0 | 0 | 0.463 | 1.697 | 0.463 | 1 | 1.697 | 1 | 0.86 | N/A |
| AT4G29310 | BIC | HKY+I | 0.24 | 0.22 | 0.23 | 0.31 | 2.39 | 1.18 | 1 | 2.394 | 1 | 1 | 2.394 | 1 | 0.85 | N/A |
| AT4G33440 | BIC | TrN+I | 0.31 | 0.17 | 0.25 | 0.27 | 0 | 0 | 1 | 1.726 | 1 | 1 | 8.945 | 1 | 0.91 | N/A |
| AT5G06580 | BIC | TPM3uf+I | 0.28 | 0.27 | 0.15 | 0.29 | 0 | 0 | 3.294 | 6.95 | 1 | 3.294 | 6.95 | 1 | 0.85 | N/A |
| AT5G06830 | BIC | HKY+I | 0.32 | 0.18 | 0.23 | 0.27 | 5.45 | 2.69 | 1 | 5.445 | 1 | 1 | 5.445 | 1 | 0.82 | N/A |
| AT5G10910 | BIC | TPM3uf+I | 0.3 | 0.19 | 0.24 | 0.26 | 0 | 0 | 2.735 | 4.181 | 1 | 2.735 | 4.181 | 1 | 0.85 | N/A |
| AT5G10920 | BIC | TIM2+I | 0.3 | 0.2 | 0.22 | 0.28 | 0 | 0 | 3.081 | 1.847 | 3.081 | 1 | 5.983 | 1 | 0.89 | N/A |
| AT5G27950 | BIC | HKY+I | 0.32 | 0.18 | 0.24 | 0.27 | 6.11 | 3.03 | 1 | 6.114 | 1 | 1 | 6.114 | 1 | 0.8 | N/A |
| AT5G41150 | BIC | HKY+I | 0.29 | 0.2 | 0.23 | 0.28 | 3.74 | 1.84 | 1 | 3.743 | 1 | 1 | 3.743 | 1 | 0.84 | N/A |
| AT5G45900 | BIC | TPM2uf+I | 0.25 | 0.21 | 0.24 | 0.3 | 0 | 0 | 0.489 | 1.367 | 0.489 | 1 | 1.367 | 1 | 0.91 | N/A |
| AT5G47090 | BIC | TPM1uf+I+G | 0.33 | 0.16 | 0.28 | 0.23 | 0 | 0 | 1 | 7.505 | 2.516 | 2.516 | 7.505 | 1 | 0.71 | 0.64 |
| AT5G54910 | BIC | HKY+G | 0.33 | 0.19 | 0.22 | 0.27 | 2.25 | 1.11 | 1 | 2.254 | 1 | 1 | 2.254 | 1 | N/A | 0.02 |
| AT5G57590 | BIC | HKY+I | 0.28 | 0.17 | 0.25 | 0.3 | 3.47 | 1.7 | 1 | 3.473 | 1 | 1 | 3.473 | 1 | 0.89 | N/A |
| AT5G64050 | BIC | HKY+I+G | 0.28 | 0.2 | 0.21 | 0.31 | 2.51 | 1.21 | 1 | 2.511 | 1 | 1 | 2.511 | 1 | 0.79 | 0.89 |
| AT5G64370 | BIC | HKY+I | 0.28 | 0.21 | 0.23 | 0.28 | 3.93 | 1.93 | 1 | 3.925 | 1 | 1 | 3.925 | 1 | 0.85 | N/A |
